# *Enterococcus faecalis*-derived lactic acid suppresses macrophage activation to facilitate persistent and polymicrobial wound infections

**DOI:** 10.1101/2025.01.31.635924

**Authors:** Ronni A. G. da Silva, Brenda Yin Qi Tien, Patrick Hsien Neng Kao, Haris Antypas, Cenk Celik, Ai Zhu Casandra Tan, Muhammad Hafiz Ismail, Guangan Hu, Kelvin Kian Long Chong, Guillaume Thibault, Jianzhu Chen, Kimberly A. Kline

## Abstract

Macrophage activation is essential for innate immunity, and its suppression enables pathogen persistence. We show that *Enterococcus faecalis* suppresses macrophage activation through lactic acid-mediated extracellular acidification. Mutants lacking lactate dehydrogenase (*ldh*), and thus unable to acidify the environment, fail to inhibit NF-κB. *E. faecalis-*derived lactic acid acts via MCT-1 and GPR81 through two distinct but complementary mechanisms that culminate in the reduction of NF-κB activity. Lactic acid acts through MCT-1 to inhibit ERK and STAT3 phosphorylation, leading to reduced STAT3 binding to the *Myd88* promoter, and reduced MyD88 protein levels. Lactic acid signaling to GPR81 mediates the phosphorylation of the transcriptional factor YAP, ultimately attenuating NF-κB signaling. In a murine wound infection model, this lactic acid-driven immunosuppressive niche enables prolonged *E. faecalis* persistence and enhances the fitness of co-infecting bacteria such as *Escherichia coli.* These findings reveal how bacterial lactic acid subverts innate immunity to support chronic and polymicrobial infections.

**Graphical Abstract:** 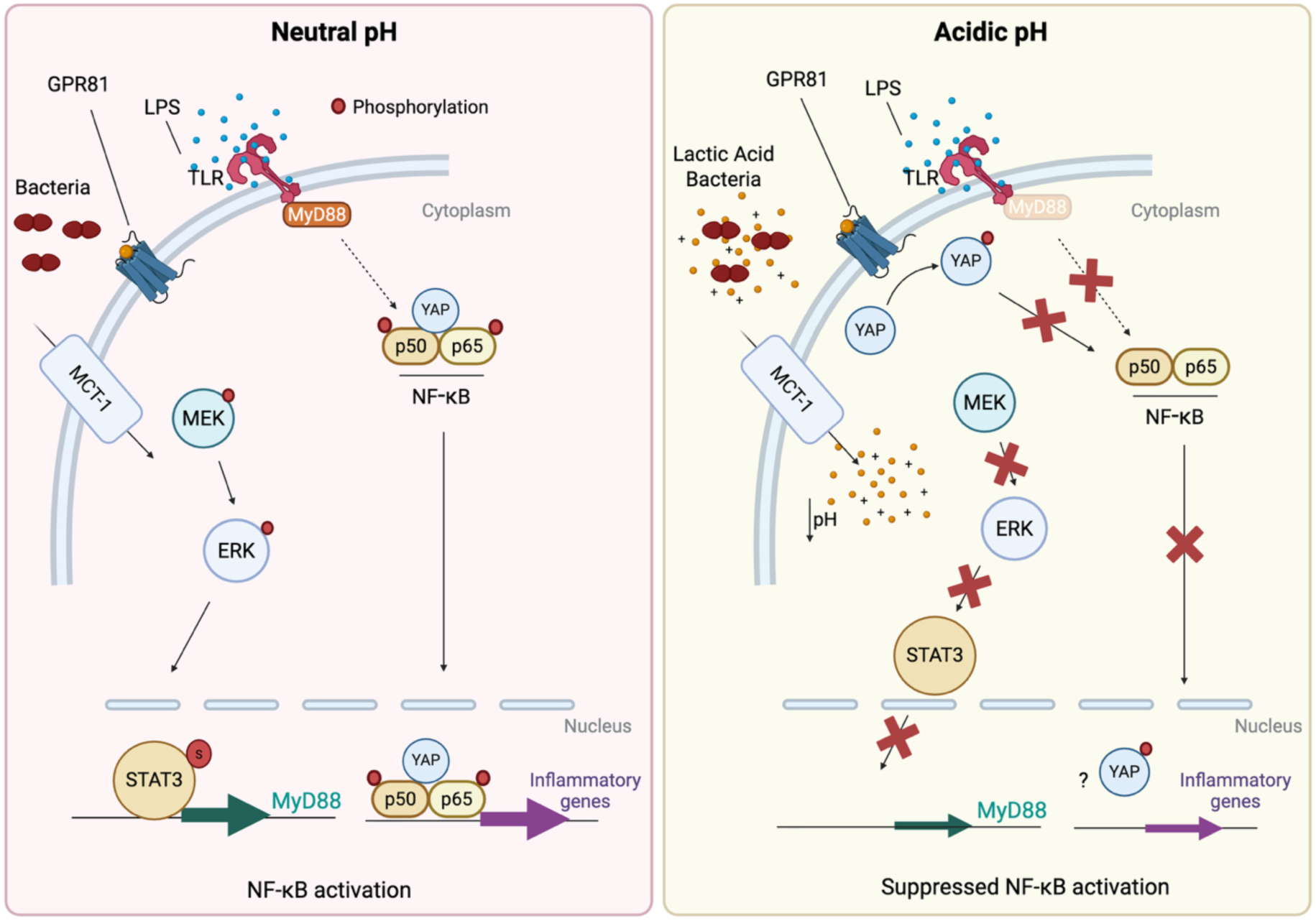

**Highlights:**

– *E. faecalis* lactate dehydrogenase mutants fail to acidify the environment and cannot suppress NF-κB signaling.
– Lactic acid is necessary and sufficient to drive NF-κB suppression in macrophages.
– Lactic acid NF-κB suppression is relayed through two complementary routes: MCT- 1 transport and the lactate sensor GPR81.
– MCT-1-dependent signaling blunts ERK/STAT3 phosphorylation, lowering MyD88 levels; whereas GPR81 drives YAP phosphorylation, reducing NF-κB activation.
– *E. faecalis*-derived lactic acid drives immunosuppression, potentiates persistence, and promotes multi-species wound infection *in vivo*.

## Introduction

*Enterococcus faecalis* are commensal, lactic acid producing, Gram-positive bacteria that are ubiquitous in the healthy human gastrointestinal tract [1]. *E. faecalis* are also opportunistic pathogens in the nosocomial environment, particularly among immunocompromised individuals, contributing to considerable morbidity and mortality in susceptible populations [2, 3]. *E. faecalis* infections are often polymicrobial in nature with two or more species present at an infection site, especially in catheter associated urinary tract infections (CAUTI) and wound infections [4–6]. Treatment of *E. faecalis* infections has become increasingly difficult due to their acquired antibiotic resistance, antibiotic tolerance associated with their ability to form biofilm in the host, and intracellular survival [4, 7–9].

The nuclear factor kappa-light-chain-enhancer of activated B cells (NF-κB) pathway is a central regulator of the immune response to bacterial and viral infections [10–12]. Stimuli, such as lipopolysaccharide (LPS) or lipoteichoic acid (LTA), are recognized by co-receptors, including CD14 and pattern recognition Toll-like receptors (TLRs), inducing TLR dimerization and cargo internalization. This process, mediated by downstream adaptor proteins such as MyD88, initiates a signaling cascade that ultimately activates the IκB kinase (IKK) complex [13]. The IKK complex phosphorylates IκB, leading to its ubiquitin-dependent degradation and release of the NF-κB transcription factor (typically a p65/p50 dimer), which then translocates to the nucleus and subsequently drives the expression of genes involved in immunity, inflammation, and cell survival [10].

Recent work highlights the importance of host metabolic and epigenetic regulators in controlling inflammatory signaling [14, 15]. The monocarboxylate transporter 1 (MCT- 1, encoded by *SLC16A1*) facilitates proton-coupled lactate transport, thereby influencing intracellular pH and redox balance [16]. MCT-1 activity can alter macrophage transcription and inflammatory responses [17]. Similarly, the G-protein-coupled lactate receptor GPR81 (also known as hydroxycarboxylic acid receptor 1, HCAR1) senses extracellular lactate to regulate AMPK–YAP signaling, which can suppress NF-κB-driven cytokine expression and dampen inflammation [18, 19]. In addition, histone deacetylases (HDACs) fine-tune NF-κB responses by deacetylating histones and NF-κB subunits such as p65/RelA, thereby influencing chromatin accessibility and transcriptional activity [20, 21]. Together, these systems link metabolic cues, pH homeostasis, and epigenetic regulation to immune signaling control in macrophages.

To colonize and persist within the human host, *E. faecalis* deploys multiple strategies to evade immune clearance, including biofilm formation to hinder phagocytosis by immune cells, and prolonged survival within macrophages and neutrophils [8, 9, 22–28]. *E. faecalis* strains isolated from the gastrointestinal tract of healthy human infants can suppress mitogen-activated protein kinase (MAPK) and NF-κB signaling, as well as interleukin 8 (IL-8) expression in intestinal epithelial cells *in vitro* [29, 30]. We previously demonstrated that *E. faecalis* infection suppresses NF-κB signaling in RAW 264.7 macrophages at high multiplicities of infection (MOI 100) [31]. In addition, *E. faecalis* suppresses lipopolysaccharide (LPS) or lipoteichoic acid (LTA)-mediated NF-κB activation in the same cell line in a dose-dependent manner [31]. However, the *E. faecalis* and host factors involved in NF-κB suppression in macrophages were not reported.

In this study, we identified *E. faecalis* lactate dehydrogenase and its lactic acid product as critical factors in macrophage suppression. *E. faecalis*-derived lactic acid lowers the extracellular pH [39]. Lower extracellular pH increases the activity of MCT-1 to transport protons and lactate into the intracellular environment driving down the intracellular pH [16, 32, 33] which impairs STAT3-dependent transcription of *Myd88*, ultimately preventing signaling transduction leading to NF-κB activation. In parallel, low extracellular pH likely enhances GPR81 affinity for extracellular lactate, in turn triggering AMPK-LATS signaling, increasing inhibitory YAP phosphorylation [19, 34], and limiting NF-κB activation. In addition, lactic acid-mediated immune suppression promotes persistent *E. faecalis* wound infection and promotes *E. coli* growth during mixed species wound infection. These findings advance our understanding of *E. faecalis* pathogenesis and identify potential targets for immunotherapeutic strategies aimed at improving treatment and prevention of recalcitrant infections.

## Results

### *E. faecalis* infection suppresses the antibacterial response of macrophages

We previously demonstrated that *E. faecalis* can suppress LPS- and LTA-mediated NF-κB activation in RAW 264.7 macrophages in a dose-dependent manner without causing host cell cytotoxicity [22]. To further investigate this immunosuppression, we infected RAW 264.7 macrophages for 6 h with *E. faecalis* strain OG1RF in the presence or absence of LPS followed by RNA sequencing. Gene set enrichment analysis (GSEA) revealed that while *E. faecalis* infection at MOI 100 enriched for pathways linked to the immune response to bacteria, *E. faecalis* suppressed pathways associated with positive immune regulation and defense responses in LPS-treated cells, **(Fig. 1A, SI 1-3)**. To explore how *E. faecalis* interferes with NF-κB signaling, we examined transcripts of key genes in this pathway. LPS alone increased transcripts that encode the LPS co-receptor CD14 [35] and the transcription factor p50 precursors p100 and p105 **(Fig. 1B, Table S1, SI 4-6)**. *E. faecalis* significantly suppressed some of these LPS-induced transcripts, with *Myd88* expression reduced by ∼2-fold in MOI 100-infected LPS-untreated macrophages and nearly 4-fold in MOI 100-infected LPS-treated macrophages **(Fig. 1B, Table S1)**. This suppression may explain why NF-κB is inactive following MOI 100 infection, despite higher transcript levels of genes encoding p100 and p105 compared to uninfected cells **(Fig. 1B, Table S1, SI 4-6)**.

**Figure 1.**
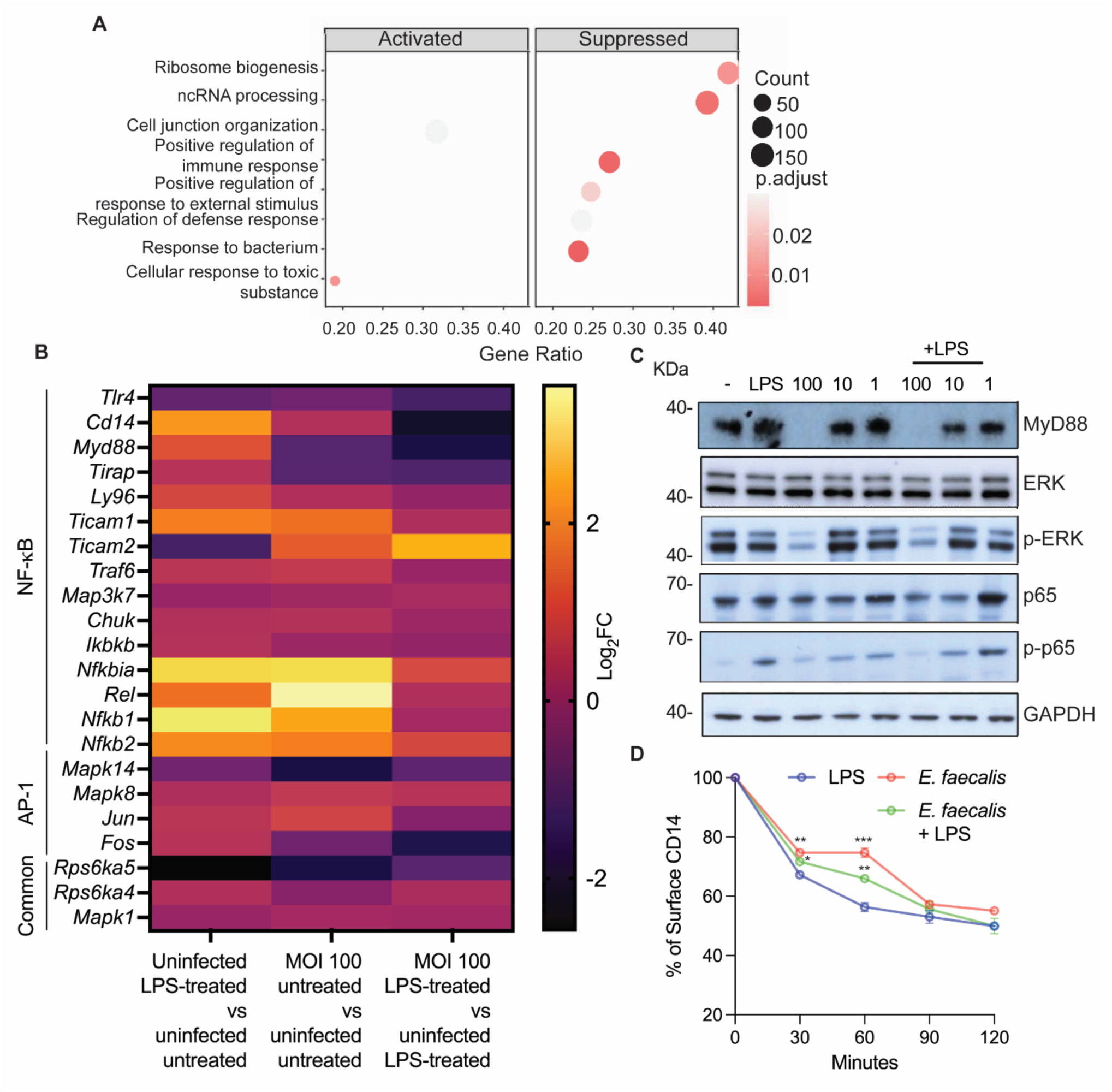
*E. faecalis* modulates the macrophage response to bacteria by reducing MyD88 protein levels and preventing ERK and p65 phosphorylation. **(A)** GSEA using gene ontology biological process gene sets showing significantly enriched pathways in RAW 264.7 macrophages infected with *E. faecalis* OG1RF for 6 h at MOI 100 in the presence of LPS (100 ng/mL) compared to uninfected LPS-treated macrophages. Activated and suppressed pathways were determined based on genes that are over and under expressed in the analyzed pathways. The size of the circle indicates the number of genes in each pathway gene set that are up- or down- regulated. **(B)** Heatmap of differentially expressed NF-κB/ AP-1-related genes. **(C)** Immunoblots performed on protein extracted from RAW 264.7 macrophages infected with *E. faecalis* for 6 h at the indicated MOI (100, 10, 1) with or without LPS and probed with antibodies specific for MyD88, ERK, p-ERK, p65, p-p65 and GAPDH. The two bands for ERK and p-ERK immunoblots represent ERK isoforms 1 and 2. Data shown are representative of at least 3 independent experiments. **(D)** RAW 264.7 cells were either infected or treated with LPS alone or in combination for the times indicated before CD14 endocytosis was examined by flow cytometry. Data (mean ± SEM) are a summary of three independent experiments and calculated relative to uninfected untreated cells which were set as 100% levels for the examined receptor. Statistical analysis was performed using ordinary one-way ANOVA followed by Dunnett’s multiple comparison test *p<0.05, **p<0.01 and ***p<0.001.

Consistently, the levels of MyD88 protein were lower in MOI 100-infected macrophages with or without LPS, as well as the downstream phosphorylated TAK1, IKK, IκBα, and p65 but not the levels of the unphosphorylated TAK1, IKKα and p65 **(Fig. 1C and Fig. S1A)**.

Since MyD88 also activates AP-1 **(Fig. S1B)**, we investigated components of this pathway, including ERK, c-Jun N-terminal kinases (JNK), p38, and AP-1 (c-Jun and c-Fos). While LPS did not affect AP-1 pathway transcript levels **(Fig. 1B, Table S1)**, MOI 100 infection reduced p38 transcripts compared to uninfected macrophages. In LPS-treated cells, MOI 100 infection decreased *Fos* (encoding c-Fos) transcripts relative to controls. Protein analysis showed significantly reduced phosphorylated ERK at MOI 100, regardless of LPS presence **(Fig. 1C)**, while total and phosphorylated levels of p38 and JNK, as well as total ERK levels were unaffected **(Fig. 1C and S1C)**. Minimal changes in phosphorylated c-Jun and c-Fos at MOI 100 suggest that *E. faecalis* likely interferes with AP-1 activation to a lesser degree than with NF-κB **(Fig. S1C)**.

To determine if *E. faecalis* affects other aspects of MyD88-related signaling, beyond suppressing *Myd88* expression, we analyzed the RNA-seq data to include a wider range of immune signaling genes. While *Tlr4* transcript levels remained unchanged by *E. faecalis* infection, expression of *Cd14* was reduced following 6 h of infection of LPS-treated cells, indicating that *E. faecalis* might also impact cargo internalization as CD14 mediates the LPS-induced internalization of extracellular molecules [23] **(Fig. 1B and Table S1)**. Although there was a significant delay in CD14 internalization at early times in *E. faecalis* infected cells, there was no marked difference in CD14 internalization at the end of the experiment, suggesting that *E. faecalis* infection-mediated suppression of NF-κB cannot be explained by changes in CD14 levels and internalization **(Fig. 1D)**. Similarly, *E. faecalis* infection had no effect on TLR2 internalization with or without LTA **(Fig. S1D)**. These data show that *E. faecalis* infection suppresses NF-κB activation in macrophages by reducing MyD88 protein levels and preventing the downstream phosphorylation cascade and leading to NF-κB suppression.

### Lactate dehydrogenase-dependent lactic acid production by *E. faecalis* drives suppression of macrophage NF-κB activation

To investigate the bacterial factors involved in suppression of macrophage activation, we screened a ∼15,000 member *E. faecalis* mariner transposon library [36] for mutants that failed to suppress LPS-induced NF-κB activation in RAW 264.7 macrophages. We identified 151 transposon mutants in our primary screen **(Fig. S2A)** which were then subjected to secondary analyses for growth defects, leading to a final list of 47 transposon mutants that failed to suppress NF-κB activation **(Fig. S2B, Table S2)**. Bacterial growth in media containing a pH indicator showed that 18 of 47 mutants failed to acidify the media as a function of growth as compared to wild type (WT) *E. faecalis* **(Fig. S2B).** Among these was *ldh1* (OG1RF_10199), encoding lactate dehydrogenase (LDH) **(Fig. S2B)**, which reduces pyruvate to L-lactate to generate NAD^+^ for ongoing glycolysis in *E. faecalis* [37, 38]. L-lactate is released from *E. faecalis* in the form of lactic acid, which is subsequently deprotonated to L-lactate, releasing a hydrogen ion (H^+^) and in turn lowering the pH in the surrounding environment. Since acidification is a defining feature of *E. faecalis* metabolism due to the release of lactic acid [39], we further investigated the impact of LDH on NF-κB activation.

The *E. faecalis* genome contains two LDH genes, *ldh1* and *ldh2,* where LDH1 plays the major role in lactic acid production under laboratory conditions as compared to LDH2 [37]. As predicted, and confirming the result of the transposon screen, infection of RAW 264.7 macrophages with Δ*ldh1* or the Δ*ldh1*Δ*ldh2* double mutant (hereafter called Ldh), failed to suppress LPS-induced NF-κB activation as compared to WT *E. faecalis*, Δ*ldh2,* or *ldh1* complementation strains **(Fig. 2A).** Immunofluorescence using confocal laser scanning microscopy (CLSM) to investigate p65 nuclear localization confirmed WT suppression and Ldh activation of NF-κB **(Fig. 2B and S3)**. Failure to suppress macrophage activation was not due to a growth defect of the mutants, whether grown in BHI or cell culture medium **(Fig. S4A-B)**. Lactic acid accumulated in culture supernatants of WT *E. faecalis* infected cells in an MOI-dependent manner, resulting in a pH decrease that is delayed for the Ldh mutant **(Fig. S4C-D)**. In contrast to WT *E. faecalis*, macrophage infection with either the Δ*ldh1* or Ldh mutants produced less lactic acid and, consequently, acidified the environment at a slower rate than WT *E. faecalis,* in agreement with lesser suppression of NF-κB **(Fig. 2A and S4E-F)**. Consistent with previous reports, Δ*ldh2* was not attenuated for lactate release [37, 40].

**Figure 2.**
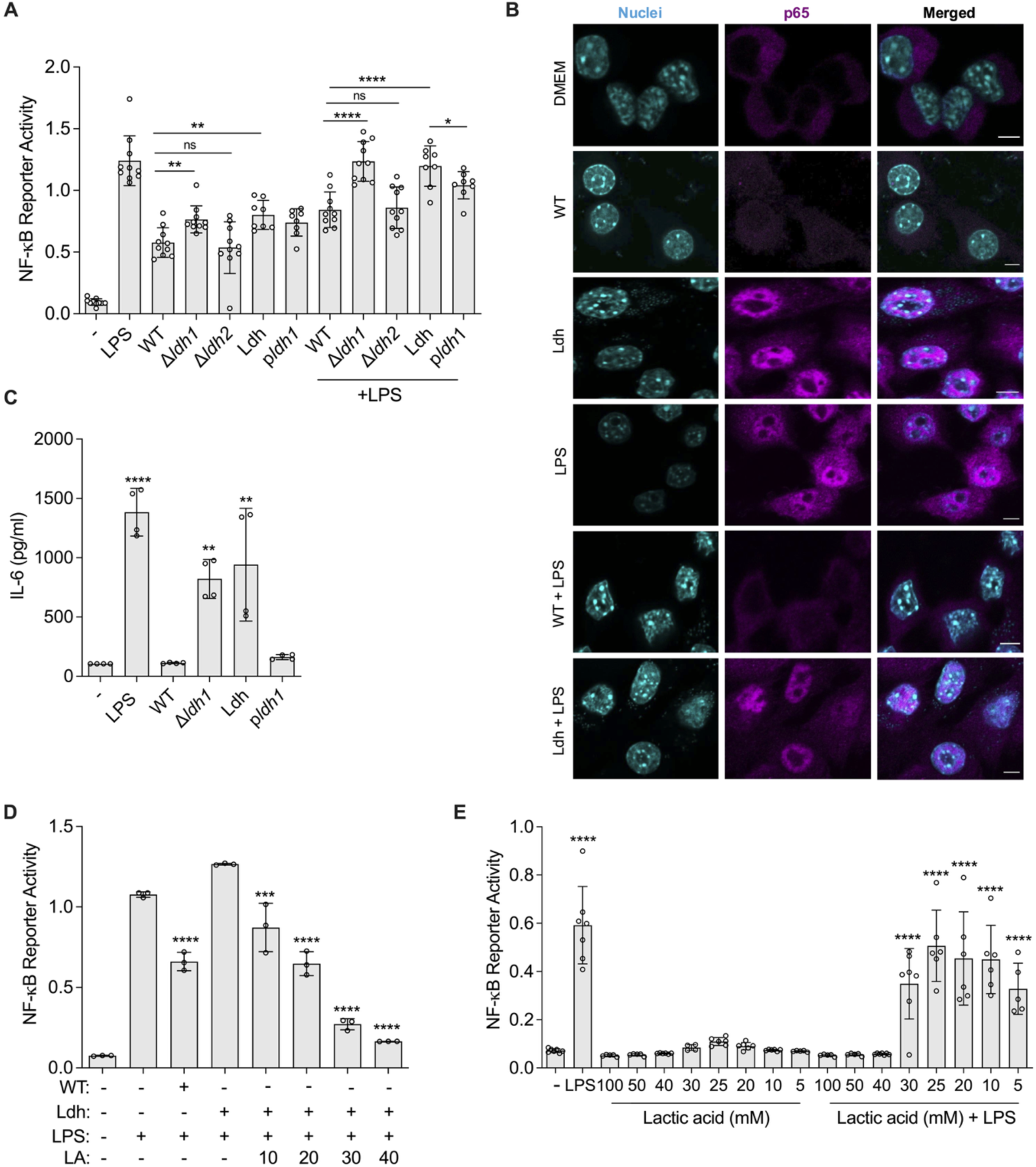
*E. faecalis* LDH is essential for suppression of NF-κB activation in macrophages. **(A)** RAW 264.7 macrophages were infected with WT, Δ*ldh1,* Δ*ldh2,* Ldh, or *ldh1* complementation strains alone or concurrently with LPS at MOI 100 for 6 h prior to measurement of NF-κB-driven SEAP reporter activity. **(B)** Immunofluorescence of p65 in RAW264.7 cells at 6 hpi infection with WT or Ldh in the presence or absence of LPS. Magenta, p65 and cyan, dsDNA stained with Hoechst 33342. Images are representative of 2 independent experiments. Scale bar: 5 μm. **(C)** BMDMs were infected with Δ*ldh1,* Ldh, or *ldh1* complemented strain for 6 h prior to measurement of IL-6 production. **(D)** RAW 264.7 cells were infected with WT *E. faecalis* or Ldh supplemented with lactic acid (LA) at various concentration (mM) for 6 h prior to measurement of NF-κB-driven SEAP reporter activity. 25 mM lactic acid best approximates the lactate concentration measured in MOI 100 supernatants in Fig. S4C. **(E)** RAW 264.7 cells were incubated with cell culture media supplemented with lactic acid at various concentrations for 6 h prior to measurement of NF-κB-driven SEAP reporter activity. Data (mean ± SEM) are a summary of at least three independent experiments. Statistical analysis was performed using one-way ANOVA with Fisher’s LSD in **(B)** or Tukey’s multiple comparison test where *p<0.05, **p<0.01, ***p<0.001 and ****p<0.0001 as indicated or as compared to media alone for **(C)** and **(E)**, and LPS for **(D)**.

To confirm that LDH-mediated macrophage suppression was not specific to RAW 264.7 macrophages, we assessed the release of the NF-κB-regulated cytokine IL-6 from murine bone marrow-derived macrophages (BMDM) infected with Δ*ldh1* and Ldh mutants compared to WT *E. faecalis* **(Fig. 2C)**. Similar to RAW 264.7 macrophages, infection with either WT or the Δ*ldh1* complemented strains had no effect on IL-6 release by BMDM, whereas infection of LPS-treated cells with either Δ*ldh1* or Ldh stimulated IL-6 production **(Fig. 2C)**. qRT-PCR analysis of representative genes showed that *E. faecalis* infection of BMDM, as in RAW264.7 cells, decreased *Myd88* expression **(Fig. S5)**. Infection of BMDM or the human monocytic cell line THP-1 with *E. faecalis* similarly showed an LDH-dependent decrease of MyD88, p-ERK, and p- p65 levels **(Fig. S6A-B)**. Multiplex cytokine analysis of RAW264.7 cells also confirmed an LDH-dependent reduction of pro-inflammatory cytokines (IL-6, IL-12 p70 and IFNγ) and chemokines (CCL3, CCL4, CCL5, CSF2 and CSF3) for *E. faecalis*-infected macrophages **(Fig. S6C)**. Furthermore, supplementation of macrophages infected with the Ldh mutant with a minimum of 20 mM lactic acid restored suppression of LPS-induced NF-κB activation to similar levels as WT *E. faecalis* **(Fig. 2D)**. In the absence of infection, 40 mM lactic acid was sufficient to suppress LPS-induced NF-κB activation **(Fig. 2E)** without compromising macrophage viability **(S6D)**. In addition, WT *E. faecalis* suppressed LPS-induced NF-κB activation even in cells pre-activated with LPS for 1 h prior to infection **(Fig. S6E-F)**. As expected, the Ldh mutant failed to suppress NF-κB activation independent of LPS pre-exposure **(Fig. S6E-F)**. Overall, these results demonstrate that *E. faecalis* suppression of NF-κB activation in macrophages is mediated by LDH-derived lactic acid.

### Extracellular acidification, rather than lactate alone, underlies NF-κB suppression during *E. faecalis* infection

Among the 47 transposon mutants that failed to suppress macrophage activation, 18 did not acidify the medium to the same extent as WT *E. faecalis*, but only 11 produced less lactate **(Fig. S2B)**, suggesting that pH reduction rather than lactic acid production alone is the primary driver of NF-κB suppression. Indeed, supplementation with 40 mM lactic acid suppressed NF-κB activation, whereas 40 mM sodium L-lactate did not **(Fig. 3A)**. To confirm this, we acidified filtered Ldh mutant supernatants to pH 6.4 (pH equivalent to WT MOI 100) or neutralized WT supernatants to pH 7.8 (pH equivalent to MOI 1). Acidified supernatants suppressed LPS-induced NF-κB activation across all MOI in the absence of cytotoxicity, while neutralized supernatants failed to do so **(Fig. 3B-C, S7A-E)**. Consistent with previous reports [41], acidifying cell culture media to pH 6.4 using HCl or phosphoric acid (H3PO4) also suppressed LPS-induced NF-κB activation without affecting macrophage viability **(Fig. 3D and S7F).** p-p65 levels were also reduced in LPS-stimulated macrophages infected with the Ldh mutant when supplemented either with lactic acid or HCl, compared to those without acid supplementation **(Fig. S7G)**.

**Figure 3.**
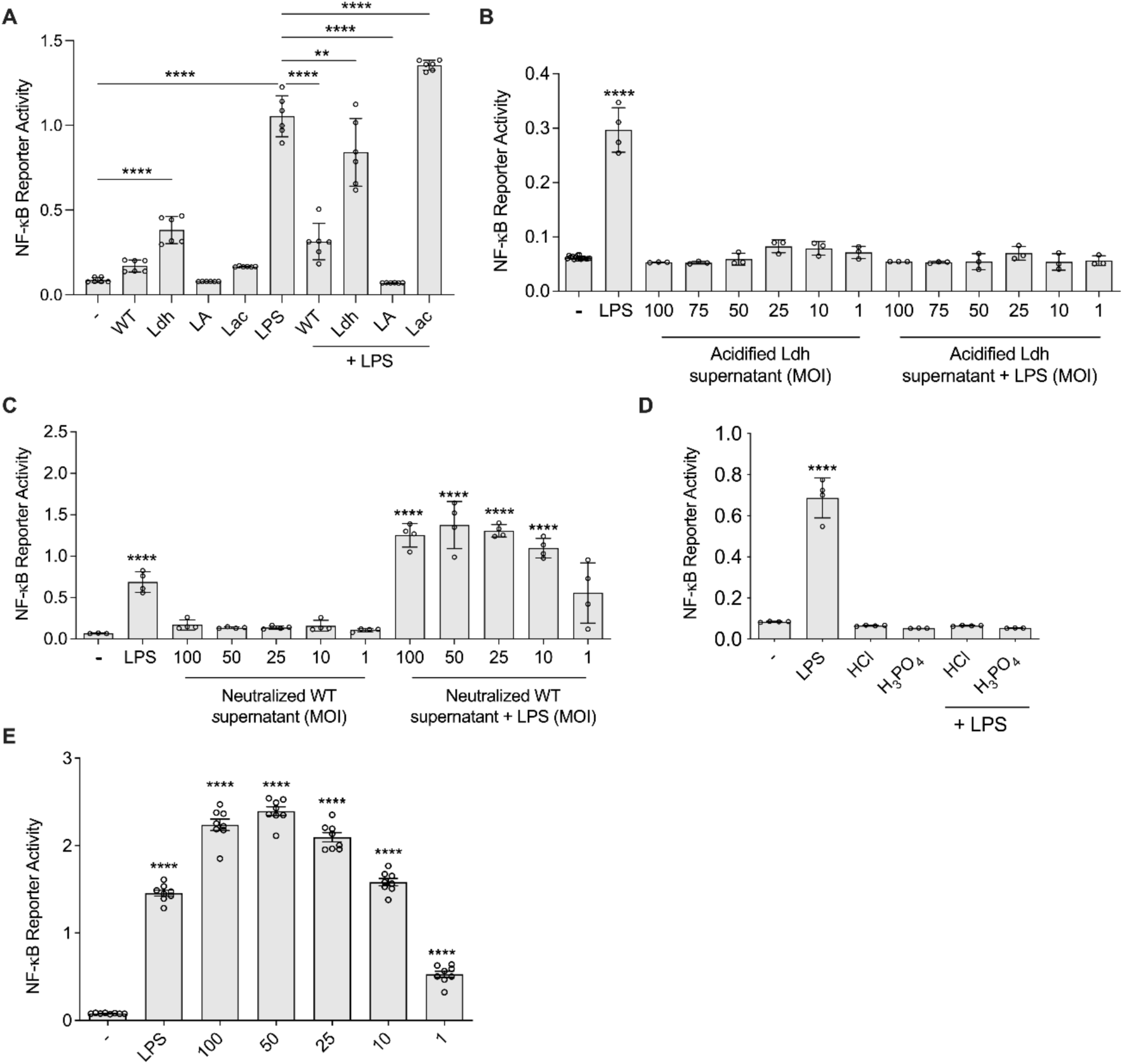
Acidification is necessary and sufficient for *E. faecalis* suppression of macrophage activation. **(A**) RAW 264.7 macrophages were infected with WT or Ldh or incubated with cell culture media supplemented with lactic acid (LA) or sodium lactate (Lac) (40 mM) in the presence or absence of LPS for 6 h prior to measurement of NF-κB-driven SEAP reporter activity. RAW 264.7 macrophages were incubated with **(B)** lactic acid-acidified, **(C)** neutralised bacterial-free supernatant (equivalent to shown MOI starting bacterial culture grown for 6 h in cell culture media in the absence of macrophages) and **(D)** HCl and H3PO4 acidified media for 6 h prior to measurement of NF-κB-driven SEAP reporter activity. **(E)** RAW 264.7 macrophages were incubated with *E. faecalis* WT (MOI 100, 50, 25, 10, 1) 3 h and treated with 500 µg/mL of gentamicin with penicillin G overnight at 37°C prior to measurement of NF-κB-driven SEAP reporter activity. **(A- E)** Data (mean ± SEM) are a summary of at least three independent experiments. Statistical analysis was performed using one-way ANOVA with Tukey’s multiple comparison tests where NS indicates p > 0.05, **p<0.01, ***p<0.001 and ****p<0.0001 as compared to media alone unless otherwise indicated.

Since *E. faecalis* can persist and replicate within macrophages [9, 42], we investigated whether intracellular *E. faecalis* contributed to NF-κB suppression. We therefore infected macrophages with *E. faecalis* for 3 h, followed by antibiotic killing of extracellular bacteria [9]. In the absence of extracellular *E. faecalis*-mediated acidification of the medium, we observed increased NF-κB activation at all MOI during intracellular infection, without significant cell death **(Fig. S8A-B).** In fact, *E. faecalis* infection resulted in higher NF-κB activation than the LPS positive control **(Fig. 3E)**. Heat and UV-killed *E. faecalis* similarly were not capable of suppressing NF-κB at high MOI **(Fig. S8C-D)**. Together, our results suggest that extracellular acidification, rather than lactate levels or intracellular infection, is important for *E. faecalis*-mediated immune suppression in macrophages.

### *E. faecalis* suppresses MyD88 expression by pH-dependent STAT3 inactivation

To understand the mechanism by which *E. faecalis*-mediated acidification impacts NF-κB activation, we focused on the reduction of MyD88 levels, which would give rise to the observed downstream signaling defects. STAT3 is a transcription factor that regulates *Myd88* expression [43]. STAT3 activation depends on its phosphorylation by ERK at serine 727 (S727) [44], preceded by ERK phosphorylation by MEK [45–47]. STAT3 activity can also be regulated by phosphorylation at tyrosine 705 (Y705) (34) and inhibited by acetylation [43]. Consistent with reduced MyD88 level, WT *E. faecalis* infection of macrophages also reduced the level of p-ERK, p-MEK and STAT3 phosphorylation at the S727 residue as compared to uninfected or Ldh-infected macrophages **(Fig. 4A)**. Total STAT3 protein levels were slightly reduced in WT-infected macrophages whereas total MEK protein levels were unchanged.

**Figure 4.**
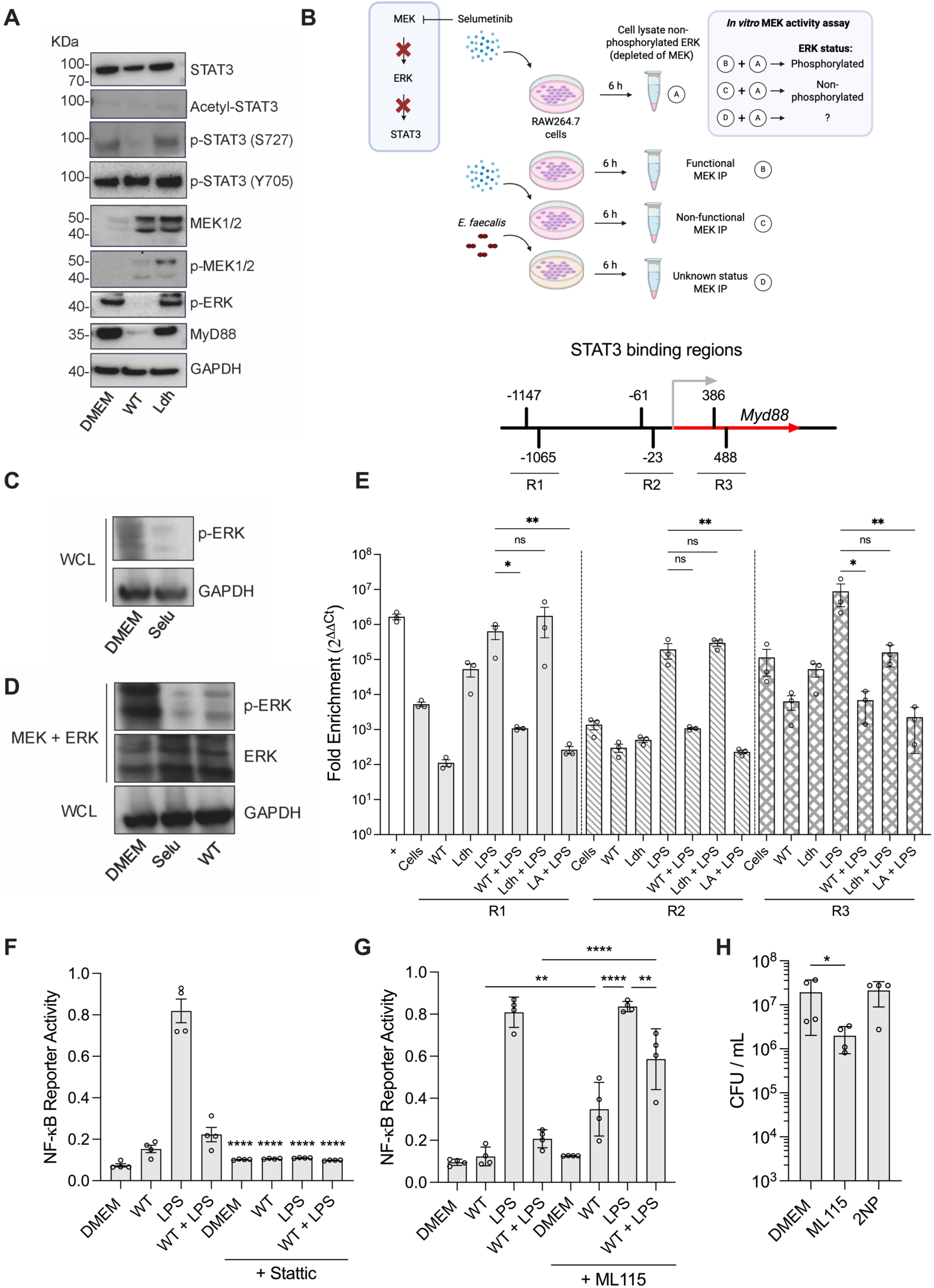
*E. faecalis* infection reduces MyD88 expression by inhibiting STAT3 activity in macrophages. (A) Immunoblot of STAT3, acetyl-STAT3, p-STAT3 (S727), p-STAT3 (Y705), MEK1/2, p-MEK1/2, p-ERK, MyD88 and GAPDH with protein extracted from RAW 264.7 macrophages infected with WT *E. faecalis* or Ldh mutant. **(B)** *In vitro* MEK activity assay. **(C-D)** pERK immunoblot of whole cell lysate (WCL) of cells treated with Selu demonstrating Selu effectively diminishes ERK phosphorylation **(C).** ERK and pERK Immunoblot post *in vitro* MEK activity assay using immunoprecipitated MEK of WT infected WCL. GAPDH immunoblot shows initial reaction input were equal **(D)**. Immunoblot images represent two independent experiments **(C-D)**. **(E)** ChIP–qPCR for STAT3 of the three regions at the MyD88 transcriptional starting site in purified DNA of RAW 264.7 cells that were infected with WT *E. faecalis* or Ldh mutant with or without LPS treatment for 6 h. Treatment with LA in the presence of LPS was also included. A depiction of the three regions relative to the transcriptional starting site (gray arrow) of *Myd88* is shown. *Myd88* coding sequence is represented in red. The positive control was established for an RNA polymerase binding site for the *Gapdh* gene. Replicates from 3 independent experiments are shown. Statistical analysis was performed using Kruskal-Wallis test followed by uncorrected Dunn’s multiple comparison tests where ns, p > 0.05; *p≤0.05 and **p<0.01. **(F-G)** RAW 264.7 macrophages were infected with the WT *E. faecalis* and incubated with cell culture media in the presence or absence of LPS and **(F)** Stattic (10 μM) and **(G)** ML115 (1 μM) for 6 h prior to measurement of NF-κB-driven SEAP reporter activity. Stattic and ML115 were added 30 min before the beginning of the experiment. Data (mean ± SEM) are a summary of at least three independent experiments. Statistical analysis was performed using one-way ANOVA with Tukey’s multiple comparison tests where ns, p > 0.05, **p<0.01, ***p<0.001 and p<0.0001 as compared to LPS unless otherwise indicated. **(H)** RAW264.7 cells were infected with *E. faecalis* for 3 h, followed by 1 h of antibiotic treatment to kill extracellular bacteria. The antibiotic-containing medium was removed, cells were washed, and fresh medium with antibiotics was added together with ML115 (1 μM) and 2NP (45 μM). Viable intracellular CFU were enumerated after 18 h incubation. Data (mean ± SEM) are a summary of at least three independent experiments. Statistical analysis was performed using ordinary one-way ANOVA followed by Tukey’s multiple comparison test ns, p>0.05; *p≤0.05.

Since MEK enzymatic activity is pH sensitive [48], we measured the intracellular pH and MEK activity in macrophages following *E. faecalis* infection. Using the pH-sensitive fluorescent reporter BCECF-AM, we observed that the intracellular pH (pHi) in macrophages infected with WT *E. faecalis* decreased to ∼6.3, whereas it remained between 6.9 and 7.2 in the Ldh-infected and non-infected cells, respectively **(Fig. S9A- B)**. To determine whether reduced pHi could affect MEK-driven signal transduction, we assessed MEK activity by immunoprecipitating MEK from WT-infected macrophages and measuring its *in vitro* phosphorylation capacity of non-phosphorylated ERK from cell lysates. We confirmed that MEK activity from infected cells was reduced **(Fig. 4B-D)**. These findings support the hypothesis that *E. faecalis*- induced changes in pHi ultimately disrupt the MEK–ERK–STAT3 signaling cascade, consistent with report of MEK signaling in fibroblasts [48].

To evaluate whether *E. faecalis* infection reduces STAT3 binding to the *Myd88* promoter, we performed chromatin immunoprecipitation (ChIP) followed by qPCR. Using the Eukaryotic Promoter Database, we identified six potential STAT3 binding sites near the *Myd88* transcription starting site that could be grouped and probed based on their proximity: region 1 (R1; -1147 and -1065), region 2 (R2; -61 and -23) and region 3 (R3; 386 and 488). STAT3 binding was reduced at these regions in WT *E. faecalis*-infected macrophages treated with LPS, whereas Ldh-infected LPS-treated macrophages showed no reduction **(Fig. 4E)**. Further, lactic acid alone was sufficient to reduce STAT3 binding at these sites. Overexpressing *Myd88* in RAW264.7 cells alleviated NF-κB suppression by *E. faecalis* **(Fig. S9G-H)**. Although immunofluorescence of WT *E. faecalis* infected macrophages in the presence or absence of LPS did not reveal major changes in STAT3 localization **(Fig. S9F)**, subcellular fractionation revealed an absence of S727-phosphorylated STAT3 in the nucleus of WT *E. faecalis* infected macrophages **(Fig. S9I)**, while Y705- phosphorylated STAT3 was detected **(Fig. S9J).** These results suggest a mechanism by which reduced STAT3 phosphorylation directly leads to reduced transcription of Myd88 [49].

We also assessed whether STAT1 contributes to NF-κB activity in infected macrophages [50]. As expected, inhibition of either MEK or STAT3 with Selumetinib or Stattic, respectively, impaired NF-κB activation in the presence of LPS, regardless of *E. faecalis* infection **(Fig. 4F, Fig. S10A)**. By contrast, STAT1 inhibition with Fludarabine reduced, but did not completely abolish, NF-κB activation **(Fig. S10B)**. Moreover, LPS-induced NF-κB activation was restored in *E. faecalis*-infected macrophages that were treated with the STAT3 agonist ML115 but not with the STAT1 agonist 2NP **(Fig. 4G and Fig. S10C)**, indicating that STAT3 plays a central role in NF-κB activation. Immunoblot of infected cell lysates treated with ML115 with or without LPS confirmed increased in MyD88 expression **(Fig. S10D)**. Functionally, treatment of macrophages with ML115 to overcome *E. faecalis*-mediated STAT3 suppression enhanced intracellular bacterial clearance by ∼10 fold, whereas treatment with 2NP did not **(Fig. 4H)**. Agonist and inhibitor concentrations did not cause cytotoxicity **(Table S3)**. Collectively, these results show that *E. faecalis* interferes with STAT3 activity to suppress *Myd88* transcription, thereby inhibiting NF-κB activation.

### MCT-1 and GPR81 mediate NF-κB suppression in *E. faecalis*-infected macrophages

Macrophages sense extracellular lactate and pH gradients via monocarboxylate transporters and G-protein coupled receptors [34, 51]. To investigate their contribution to *E. faecalis*-mediated immune suppression, we silenced the genes encoding MCT- 1, GPR81 (also known as hydroxycarboxylic acid receptor 1), and GPR65 in RAW 264.7 macrophages. Knockdown was confirmed by RT-PCR and viability of knockdown cells remained >80% **(Fig. S11F-H and S12)**. Individual knockdowns partially restored NF-κB activation, whereas combined *Mct-1*+*Gpr81*, or a triple knockdown of *Mct-1+Gpr81+Gpr65*, fully restored NF-κB activation in the presence *E. faecalis* and LPS, while non-targeting controls had no effect **(Fig. 5A-B, Fig. S11A-E)**. These results identify MCT-1 and GPR81 as the main mediators of lactate/pH sensing during *E. faecalis* infection.

**Figure 5.**
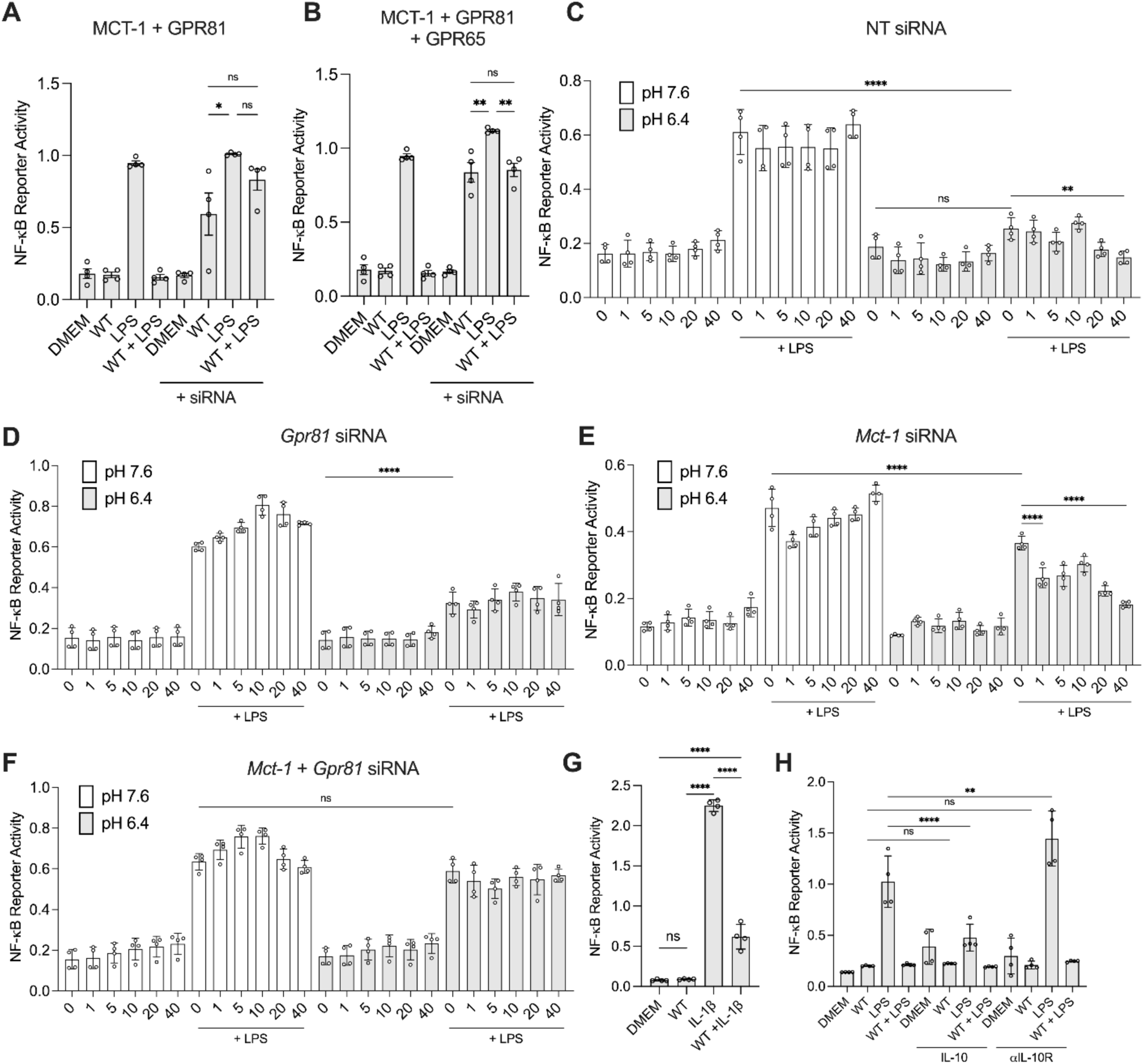
The pH-sensitive lactate transporter MCT-1 and lactate sensor GPR81 mediate suppression of NF-κB by *E. faecalis*. **(A-B)** RAW 264.7 macrophages, with or without siRNA-silenced lactate transporters and sensors, were infected with *E. faecalis* in the presence or absence of LPS for 6 h prior to measurement of NF-κB-driven SEAP reporter activity. **(C-F)** Non-targeting, *Gpr81* (*Hcar1*), *Mct-1* (*Slc16a1*) siRNA alone or combined were used to silence RAW 264.7 cells in pH 7.6 or pH 6.4 in the presence or absence of LPS, in increasing concentrations of lactate, for 6 h prior to measurement of NF-κB-driven SEAP reporter activity. **(G)** RAW 264.7 macrophages were infected with the WT *E. faecalis* and incubated in cell culture media in presence or absence of IL-1β for 6 h prior to measurement of NF-κB-driven SEAP reporter activity. **(H)** NF-κB reporter activity assay with RAW 264.7 cells uninfected or 6 h WT infected that were left untreated or treated with LPS. RAW 264.7 cells were pre-treated with IL-10 or antibody αIL-10R for 30 min prior to the beginning of the assay. Data (mean ± SEM) represent at least two independent experiments. Statistical analysis was performed using ordinary one-way ANOVA followed by Tukey’s multiple comparison test **(A-B and D-G)** or Fisher’s LSD **(C and H),** ns, p>0.05; *p≤0.05; **p<0.01 and ****p<0.0001.

To define the mechanism by which MCT-1 and GPR81 contribute to NF-κB regulation, we focused on GPR81-YAP and MCT-1-pH signaling. We hypothesized that GPR81 activation by extracellular lactate promoted YAP phosphorylation, limiting its nuclear cooperation with p65 [19], and thereby attenuating NF-κB activation. Consistent with this, the YAP agonist PY60 increased NF-κB activity in *E. faecalis-*infected cells, whereas the YAP inhibitor verteporfin blunted NF-κB activation **(Fig. S13A-B)**. Although the lactate-driven YAP response is reported to be adenylyl-cyclase independent [19], and lactate is reported to reduce cAMP and inhibit NF-κB [52], infection did not alter intracellular cAMP **(Fig. S13C)**, supporting a possible noncanonical GPR81-YAP pathway. Combining PY60 with the adenylyl cyclase inhibitor MDL-12330A (MDL-12) further increased LPS-driven NF-κB activation, independent of infection, suggesting that GPR81-mediated YAP phosphorylation dominates during infection **(Fig. S13D)**. Immunofluorescence and subcellular fractionation confirmed altered pYAP localization and increased nuclear YAP **(Fig. S9I and Fig. S13E)**.

MCT-1 imports lactate in a proton-dependent manner, reducing intracellular pH [53]. At pH 6.4, NF-κB activation was suppressed in LPS-treated siRNA control cells, and addition of lactate further enhanced suppression **(Fig. 5C)**. In *Gpr81*-silenced cells, lactate did not increase suppression, indicating that MCT-1 alone can mediate NF-κB inhibition under acidic conditions **(Fig. 5D)**. By contrast, *Mct-1* knockdown cells showed stronger NF-κB suppression with increasing lactate at low pH, demonstrating that GPR81 activity requires both acidity and lactate **(Fig. 5E)**. Notably, DMEM supplemented with 10% FBS contains baseline lactate that may activate GPR81 under low pH, contributing to NF-κB inhibition **(Fig. S14A)**. Finally, dual *Mct-1/Gpr81* knockdown restored NF-κB activation, with any residual suppression likely mediated by other pH sensors such as GPR65 **(Fig. 5F)**.

To further clarify the contribution of MCT-1 to NF-κB suppression, we used the MCT- 1/4 inhibitor α-cyano-4-hydroxycinnamate (CHC), the selective MCT-1 inhibitor AZD3965, and the MCT inhibitor 7ACC2, either alone or in combination with the adenylyl cyclase inhibitor MDL-12, in the presence or absence of *E. faecalis* and LPS **(Fig. S14)**. MDL-12 was included to keep intracellular cAMP levels low, thereby ruling out residual cAMP-driven effects and focusing on pH- and transport-dependent mechanisms. Consistent with its role in NF-κB activation, MDL-12 alone reduced NF- κB reporter activity under neutral pH conditions **(Fig. S14B)**. In infected macrophages, MCT-1 inhibition modestly relieved suppression when using AZD3965 or 7ACC2, but not CHC **(Fig. S14C-D)**. Importantly, combining MCT inhibitors with MDL-12 restored NF-κB activation to levels comparable to LPS stimulation alone **(Fig. S14D-E)**. These results support that MCT-1 activity contributes to NF-κB suppression and that the effect is independent of canonical cAMP signaling, as blocking both lactate transport and adenylyl cyclase eliminates the suppression phenotype.

### NF-κB suppression by *E. faecalis* occurs independently of HIF-1α, IL-10, and HDAC pathways

We also considered whether the pH sensor HIF-1α [54] was involved in *E. faecalis* immunosuppression of macrophages. HIF-1α was stabilized in WT *E. faecalis*-infected macrophages, enhancing IL-1β production **(Fig. S15A-D)**. Hypoxia response element (HRE) reporter assays confirmed enhanced HIF-1α activity during infection and an IL- 1R reporter assay similarly confirmed bioactive IL-1β, which was reduced with HIF-1α inhibition by LW6 and enhanced by the HIF-1α activator O-phenanthroline **(Fig. S15E- G)**. However, NF-κB remained suppressed despite elevated IL-1β during infection **(Fig. 5G)**, in line with the ability of *E. faecalis* to disrupt MyD88 expression and thereby impair IL-1β signaling, which depends on MyD88 **(Fig. S15H)**.

IL-10 activates IL-10R, triggering JAK1/TYK2-dependent STAT3 Y705 phosphorylation **(Fig. S16A)** [55]. Since IL-10 is produced during *E. faecalis* wound infection regardless of strain **(Fig. S16B-D)**, we asked whether infection-derived IL-10 could contribute to NF-κB suppression during infection. We first exposed NF-κB reporter cells ± LPS to 20 μL bacteria-free supernatants from WT *E. faecalis*-infected macrophages. These supernatants activated NF-κB **(Fig. S16E-F)**, consistent with our prior findings on IL-1β signaling, suggesting that infection-derived IL-1β overrides, or that endogenous IL-10 is insufficient to drive macrophage suppression. By contrast, supernatants from IL-10-supplemented infections suppressed LPS-driven NF-κB activity, consistent with carry-over of exogenous IL-10, while blockade with anti-IL-10R yielded a response similar to untreated supernatant **(Fig. S16F)**. Finally, we directly infected NF-κB reporter cells with WT *E. faecalis* under all combinations of LPS, IL-10, and αIL-10R. NF-κB suppression remained robust regardless of IL-10 modulation **(Fig. 5H)**. These results indicate that although IL-10 is produced during infection, it plays little to no role in *E. faecalis*-mediated NF-κB suppression.

Finally, we measured HDAC contribution to lactic acid-mediated NF-κB suppression in macrophages during *E. faecalis* infection. WT *E. faecalis* infection did not alter global H3 acetylation, whereas the Ldh mutant reduced it **(Fig. S17A).** Pan HDAC inhibition with trichostatin A (trichoA) partially restored NF-κB activation in LPS-treated, *E. faecalis*-infected macrophages **(Fig S17B)** and HDAC6 inhibition with tubastatin A (tubA) modestly relieved suppression, but inhibition of other isoforms had no effect **(Fig S17C-G)**. Together these results show that *E. faecalis*-derived lactic acid suppresses NF-κB primarily through MCT-1-mediated proton import and GPR81- driven YAP phosphorylation, whereas HIF-1α/ IL-1β responses occur but do not rescue NF-κB activation and HDAC activity contributes only marginally.

### *E. faecalis* LDH contributes to *E. faecalis* persistence in wound infections

*E. faecalis* infection suppresses the immune response and impairs healing in a murine model of wound infection [56]. We predicted that LDH-mediated immune modulation is important for *E. faecalis* survival in wounds. To test this, wounds were inoculated with either WT *E. faecalis* or the Ldh mutant. Both strains showed similar survival at 24 hpi **(Fig. 6A)**, despite LDH-dependent reduction of lactate in wound tissues at the same time point **(Fig. S18A),** but in agreement with *in vitro* data that show no difference between WT and Ldh survival in RAW264.7 cells and BMDMs 24 hpi **(Fig S18B)**. By 7 dpi, however, Ldh CFU were ∼1-log lower than WT **(Fig. 6B)**, indicating that LDH primarily supports *E. faecalis* persistence rather than acute infection. Supporting this, a modified killing assay maintaining extracellular acidosis revealed reduced macrophage bactericidal capacity at low pH, independent of strain **(Fig. S18C)** or the source of bacterial-free supernatant **(Fig. S18D)**. Importantly, combined inhibition of MCT-1 with AZD3965 and YAP activation with PY-60 enhanced bacterial killing in both RAW264.7 cells and BMDMs **(Fig. S18E)**.

**Figure 6.**
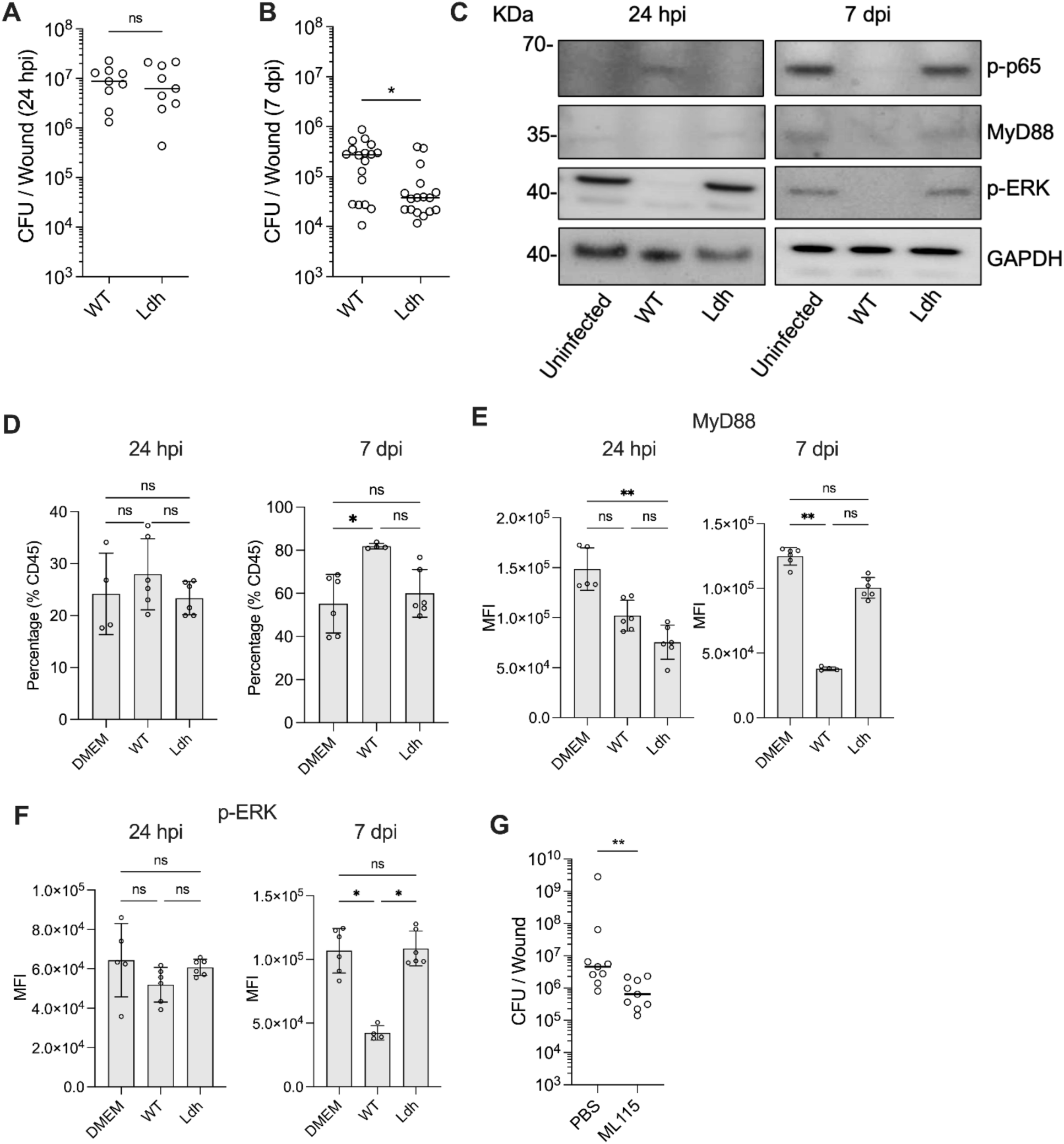
***E. faecalis* LDH activity contributes to persistence in the wound. (A-B)** Using a wound model, mice were infected with 10^6^ CFU each of WT or Ldh mutant and CFU were enumerated at 24 hpi and 7 dpi. Each symbol represents one mouse with the median indicated by the horizontal line. Data are from at least 2 independent experiments with three to five mice per experiment. Statistical analysis was performed using the nonparametric Mann-Whitney test to compare ranks; ns, p>0.05, *p≤0.05. **(C)** Immunoblot performed on protein extracted from harvested wound lysates of WT or Ldh infections 24 hpi or 7 dpi and probed with antibodies against p-p65, MyD88, p- ERK and GAPDH. **(D)** Percentage of CD11b^+^ F4/80^+^ macrophages recovered from uninfected or infected wounds with WT or Ldh after 24 hpi or 7 dpi. Comparison of mean fluorescence intensity (MFI) of MyD88 **(E)** and p-ERK **(F)** staining gating on CD11b^+^ F4/80^+^ macrophages recovered from uninfected or infected wounds with WT or Ldh after 24 hpi or 7 dpi. **(D-F)** Each dot represents one mouse. Data (mean ± SEM) are a summary of at least two independent experiments with two to four mice per experiment. Statistical analysis was performed using Kruskal-Wallis test with Dunn’s posttest; ns, p>0.05; \**p*≤ 0.05; and **p≤ 0.01. **(G)** Mouse wounds were infected with 10^6^ CFU of WT *E. faecalis* and CFU were enumerated at 24 hpi after treatment with either PBS or 10 μL of ML115 (100 μM). Each symbol represents one mouse with the median is indicated by the horizontal line. Data are from at least 2 independent experiments with three to five mice per experiment. Statistical analysis was performed using the nonparametric Mann-Whitney test to compare ranks; **p<0.01.

To define how LDH influences immune signaling *in vivo*, we examined p-p65, p-ERK, and MyD88 in wound tissues. WT *E. faecalis* infection, but not Ldh mutant, reduced p-ERK at 24 hpi and all three proteins at 7 dpi **(Fig. 6C)**. Flow cytometry revealed that macrophages expanded to ∼80% of immune cells at 7 dpi in WT *E. faecalis* -infected wounds versus ∼25% at 1 dpi **(Fig. 6D, S18F)**, coinciding with the stronger reduction in signaling proteins **(Fig. 6C)**. Direct analysis of macrophages confirmed marked reductions in MyD88 and p-ERK at 7 dpi in WT *E. faecalis*-infected wounds compared to uninfected or Ldh-infected wounds **(Fig. 6E-F)**. By contrast, topical application of the STAT3 agonist ML115 on WT-infected wounds reduced *E. faecalis* CFU by 1.1- log compared to PBS treatment **(Fig. 6G)**, further supporting the model that *E. faecalis* suppresses macrophages killing via MyD88 downregulation. Altogether, these results demonstrate that LDH contributes significantly to persistent *E. faecalis* wound infection by promoting an acidic environment that impairs macrophage killing and dampens immune signaling pathways.

### *E. faecalis* LDH-mediated immune suppression promotes *E. coli* survival during mixed species infection

We previously showed that co-infection with *E. faecalis* enhances *E. coli* virulence and growth in a mouse wound infection model [57]. To test whether immune suppression contributed to this phenotype, we co-infected RAW 264.7 macrophages with *E. coli* and either WT or Ldh *E. faecalis* and observed higher NF-κB activation for the Ldh co-infection compared to WT **(Fig. 7A)**. Similarly, WT *E. faecalis* supernatants suppressed NF-κB activation by other species, including *E. faecium*, *P. aeruginosa* (PA), *S. aureus* (USA300), *L. monocytogenes* (LM), and *S. Typhimurium* (ST), highlighting the potential of *E. faecalis* to promote a variety of polymicrobial infections **(Fig. 7B)**. With the exception of the *E. faecalis* Ldh mutant, *P. aeruginosa* PAO1, and *L. monocytogenes* ATCC19115, each of these species also acidified the medium to below pH 7 **(Fig. S19A)**. However, only enterococci and *S. aureus* produced detectable amounts of lactic acid **(Fig. S19B)** and suppressed LPS-induced NF-κB activation at high MOI **(Fig. S19C)** in the absence of marked cytotoxicity **(Fig. S19D)**. In a murine wound model, co-inoculation with *E. coli* and either WT *E. faecalis* or the Ldh mutant yielded similar *E. faecalis* CFU at 24 hpi **(Fig. 7C)**, whereas *E. coli* growth was significantly enhanced during co-infection with WT, not the mutant **(Fig. 7D)**. To exclude *E. faecalis*-derived lactate as a nutrient source for *E. coli*, we tested *E. coli* growth *in vitro* with increasing lactate concentrations and observed no enhancement **(Fig. S19E)**. Altogether, these findings demonstrate that LDH-dependent lactic acid production by *E. faecalis* suppresses host NF-κB signaling, thereby facilitating *E. coli* proliferation during wound infection. Lactic acid-driven immune suppression may thus represent a conserved strategy among certain lactic-acid producing species to modulate host responses and promote polymicrobial persistence.

**Figure 7.**
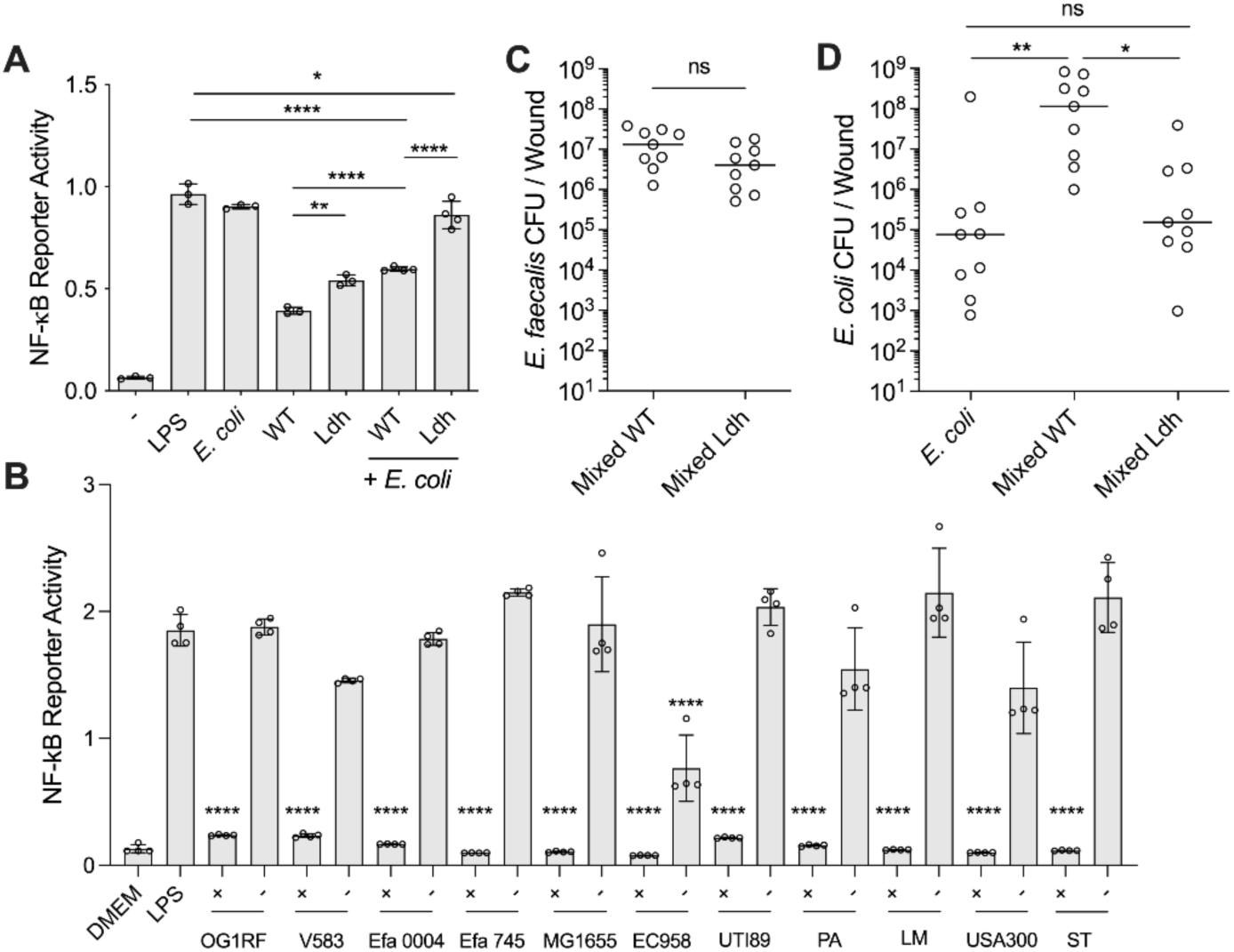
*E. faecalis* LDH-mediated immune suppression promotes *E. coli* growth during polymicrobial infection. **(A)** RAW 264.7 macrophages were infected with *E. coli* strain UTI89 at MOI 0.125, WT *E. faecalis* or Ldh mutant at MOI 100 in single species (control) or mixed species infection for 6 h prior to measurement of NF-κB-driven SEAP reporter activity. **(B)** RAW 264.7 macrophages were infected with different bacterial species as indicated at MOI 1 for 6 h in the presence or absence of WT supernatant prior to measurement of NF-κB-driven SEAP reporter activity. **(A-B)** Data (mean ± SEM) are summary from at least 3 independent experiments. Statistical analysis was performed using ordinary one-way ANOVA, followed by Tukey’s multiple comparison test; ns, p>0.05; \*\*\**p*≤ 0.001; and ****p≤ 0.0001. Mouse wounds were infected with 10^2^ CFU/wound *E. coli* UTI89 or 10^6^ CFU each of WT *E. faecalis* or Ldh mutant in single-species control for UTI89 or mixed species infections. 24 hpi CFU enumeration was performed for **(B)** *E. faecalis* and **(C)** *E. coli*. Each symbol represents one mouse with the median indicated by the horizontal line. Data are from at least 2 independent experiments with four to five mice per experiment. Statistical analysis was performed using the nonparametric Mann-Whitney test to compare ranks **(B)** and using Kruskal-Wallis test with Dunn’s posttest **(C)**; ns, p>0.05; \**p*≤ 0.05; and **p≤ 0.01.

## Discussion

*E. faecalis* is among the top pathogens associated with hospital acquired infections including CAUTI, central line-associated bloodstream infection, and surgical site infection [4, 58]. While previous studies focused on host inflammatory responses at lower MOI [29, 59], we and others have shown that *E. faecalis* exhibits immunomodulatory properties at higher MOI via mechanisms and bacterial factors that were previously unclear [30, 31, 60]. Here we demonstrate that *E. faecalis* infection at MOI 100 suppresses macrophage activation by interfering with STAT3 and YAP activity, resulting in reduced MyD88 levels and inhibition of downstream signaling events required for NF-κB activation. Additionally, we identify *E. faecalis* lactate dehydrogenase (LDH) and its lactic acid product as critical factors for macrophage suppression and persistence in wounds, as well as for promoting *E. coli* growth during co-infection.

Lactic acid production is a metabolic hallmark of *E. faecalis* fermentation [61]. *E. faecalis* LDH catalyzes the reduction of pyruvate to L-lactate, which is exported as lactic acid and deprotonates into L-lactate at physiological pH, reducing the local pH. Here we show that *ldh1* is necessary, and its byproduct (lactic acid) is sufficient, to suppress LPS-induced NF-κB activation. This may be why LDH-deficient *E. faecalis* exhibit reduced virulence in systemic murine infections [38]. Similarly, LDH is important for virulence of *S. aureus* in a murine sepsis model [62]. Lactic acid-induced immunosuppression appears widespread; other lactic acid bacteria, such as *Lactobacillus*, suppress NF-κB activation in macrophages and myometrial cells [63, 64]. While we focused on the role of LDH in this study, we identified additional mutants that failed to suppress LPS-induced NF-κB activation despite acidifying the environment, suggesting additional immune suppressive mechanisms. Beyond infection, lactic acid contributes to immune suppression in tumor microenvironments, inducing tissue acidosis that suppresses NF-κB activity, through mechanisms involving pH changes and histone acetylation [17, 65–75]. Here we show that *E. faecalis* lactic acid-mediated suppression of MyD88 levels and downstream signaling events can be mimicked by acidic conditions, driven by pH effects rather than phosphorylation changes. These mechanisms may be context dependent and warrant further study.

The MEK-ERK-STAT3 axis is disrupted by *E. faecalis*-derived lactic acid, reducing STAT3 phosphorylation and binding to the *Myd88* promoter, consistent with previous reports [43]. While other bacterial infections activate the MEK/ERK cascade for oxidative and nitrosative bursts, endosomal trafficking, and proinflammatory macrophage polarization [76–78], *E. faecalis* subverts this defense [79, 80]. *Salmonella typhimurium* and *Helicobacter pylori*, for example, induce STAT3 phosphorylation via secreted factors to alter immune responses [79]. We show that pH sensitive kinases, including MEK and ERK, are disrupted at lower intracellular pH, inhibiting STAT3 activity in alignment with previous studies [81]. Indeed, pH 6.2-6.4 also reduces MEK and ERK phosphorylation in fibroblasts and prevent NF-kB activation in macrophages [48, 82].

How then does extracellular *E. faecalis*-derived lactic acid exert these pH-driven intracellular effects? We identified the host lactate transporter MCT-1 and the lactate sensor GPR81 as mediators of NF-κB suppression during *E. faecalis* infection. MCT- 1 import of lactate, other metabolites and protons was previously shown to mediate the inhibition of HDAC11 activity that leads to the activation of HDAC6 to promote IL- 10 release [83]. Here we show that MCT-1 transport of protons is driving intracellular pH down which disrupts the MEK-ERK-STAT3 phosphorylation signaling cascade leading to lower MyD88 expression. At the same time, GPR81 sensing of extracellular lactate prevents the activation of the transcription factor YAP and, ultimately, leads to NF-κB suppression [19, 83]. Classically, lactate activates GPR81 to lower cAMP, whereas lower cAMP via PKA activates LATS1/2 to phosphorylate YAP and retain it in the cytoplasm. Therefore, logically, a simple drop in cAMP would be expected to decrease, not increase, YAP phosphorylation leading to greater NF-κB activation [84]. However, lactate-GPR81 signaling increases YAP phosphorylation in macrophages likely in an adenylyl cyclase-dependent manner, consistent with the hypothesis that GPR81 can drive AMPK-LATS signaling pathway to rather inactivate YAP and dampen NF-κB [19]. In line with this model, *E. faecalis* infection did not measurably reduce intracellular cAMP in our system, and the adenylyl-cyclase inhibitor MDL-12 reduced rather than increased NF-κB activation in uninfected and infected macrophages. Interestingly, we also detected nuclear pYAP signal, although Ser127 phosphorylation typically promotes cytoplasmic retention, suggesting non-canonical nuclear YAP pools during infection.

Expression of the genes encoding MCT-1 and GPR81 is pH-dependent [25–27]. Moreover, both MCT-1 and GPR81 may be activated at lower pH [34, 85]. Lactate supplementation alone did not replicate *E. faecalis* LDH-mediated immune suppression, suggesting that MCT-1/GPR81 activation is indeed pH dependent. Initially, our study seems to contrast a study showing that lactic acid can induce phosphorylation of ERK in rat hepatocyte RLN-10 cells via GPR81 sensing [86]. However, that study was conducted under adjusted neutral pH, excluding the potential effect of pH on ERK phosphorylation. Similarly, studies showing increased MEK and ERK phosphorylation via GPR81 used sodium lactate, which does not lower pH in their experiments [87, 88]. Conversely, others have observed that acidosis (pH 4-5) can activate ERK2, but not ERK1, in human epidermoid squamous carcinoma A431 cells [89], whereas pH 6.6 can increase the phosphorylation of ERK after 2 h of incubation but gradually inhibit phosphorylation after 4 h in human prostate carcinoma epithelial 22Rv1 cells [90]. Similarly, pH 6.3 was observed to inhibit ERK phosphorylation in Jurkat cells after 24 h [91]. These studies are consistent with our model that pH affects the phosphorylation status of these kinases. *E. faecalis* wound infections are immune suppressive and delay wound healing [42].

We show that *E. faecalis* LDH activity drives immune suppression, *E. faecalis* persistence in wounds, as well as the growth of *E. coli* during co-infection. Given the prevalence of infections caused by lactic acid-producing bacteria [92, 93], as well as their beneficial contributions to healthy microbiomes, further studies are needed to delineate how extracellular pH and lactate mechanistically shape polymicrobial interactions. We found that several bacterial species can acidify their environment, often through metabolites other than lactate, although many exhibited cytotoxicity at higher MOI. At lower MOI, when these species activated NF-κB, *E. faecalis* supernatant alone was sufficient to suppress this activation. Together, these findings reveal *E. faecalis* as a paradigm for metabolic immune modulation, demonstrating how bacterial acidification can reprogram host signaling to foster interspecies coexistence and persistence within infected tissues. Importantly, compounds that override this suppression and activate macrophage NF-κB signaling during *E. faecalis* infection enhance macrophage intracellular killing of bacteria [94, 95]. Here, we observed that the STAT3 agonist ML115 was also effective both *in vitro* and *in vivo* to reduce the bacterial burden of *E. faecalis* infection, further highlighting the therapeutic potential of immunomodulation for immunosuppressive bacterial infections. Overall, a comprehensive understanding of the immunological pathways affected by *E. faecalis* infection could inform new strategies to mitigate persistent infections.

## Materials and Methods

### Bacterial strains and growth conditions

*E. coli* strains used in this study were grown overnight in Luria-Bertani (LB) (Difco) broth or agar at 37°C under static conditions. *E. faecalis* strain OG1RF or *ldh* mutants and other bacterial strains **(Table S4)** were grown statically in brain heart infusion (BHI) (Becton, Dickinson, USA) broth or agar at 37°C overnight. Overnight cultures of bacteria were centrifuged at 6,000 *g* for 5 minutes and resuspended in PBS at OD600 0.7 (2 x 10^8^ CFU/mL) for *E. faecalis* and at OD600 0.4 (2 x 10^8^ CFU/mL) for *E. coli*.

### Genetic manipulations

To create deletion mutants, we used a thermosensitive shuttle plasmid pCGP213 which was isolated using the Monarch Plasmid Miniprep Kit (New England BioLabs Inc., USA) following the manufacturer’s instruction [96]. OG1RF genomic DNA is isolated using Wizard Genomic DNA Purification Kit (Promega, USA) and Phusion DNA polymerase (Thermo Fisher Scientific, Singapore) was used to amplify approximately 1 kb upstream and downstream of the gene of interested according to the manufacturer’s protocol. Primers used in the study are listed in **Table S5**. Subsequently, the amplified products were sewn together using the primers dLDH1_For, dLDH1_Rev, dLDH2_For and dLDH2_Rev. *ldh1* and *ldh2* constructs were then subcloned individually into pGCP213 using In-Fusion HD Cloning Kit using (Clontech, Takara, Japan) following manufacturer’s instruction. Deletion constructs were then transformed into *E. coli* and correct transformants were selected at 37°C under erythromycin (Sigma-Aldrich, St Louis, MO) pressure 500 µg/mL. The amplified plasmids were then purified from *E. coli* using the Monarch Plasmid Miniprep Kit. The plasmids were then transformed into OG1RF using electroporation, and the correct transformants were selected at 30°C under erythromycin pressure 25 µg/mL. Chromosomal integrations were selected by passaging 37°C in the absence of erythromycin. The loss of *ldh1* or *ldh2* is then verified by PCR using primer pairs LDH1_F, LDH1_R, LDH2_F and LDH2_R. Δ*ldh1*Δ*ldh2* was created by subcloning the pGCP213-*ldh1* deletion construct into Δ*ldh2*. All deletion constructs were subsequently confirmed using whole genomic sequence to ensure that only the gene of interest is deleted in the absence of any spontaneous mutation.

### Growth kinetics and pH curve

Bacterial cultures were grown as described above and overnight cultures were normalized to obtain a final concentration of 10^3^ cells/mL before inoculating at 1:25 ratio into 200 µL of either BHI or DMEM supplemented with 10% FBS. Subsequently, growth rates were measured by recording OD 600 over 20 h at 37°C using a microplate reader (Tecan infinite M200 Pro, Switzerland).

For pH measurements, bacterial cultures were grown and normalized to 3.4 x 10^7^ cells/mL (assuming MOI 100 for 10^6^ macrophages) in DMEM supplemented with 10% FBS. The pH was then measured at 1-h interval for up to 6 h, and subsequently measured at 8 h and 24 h using a digital pH meter (Sartorius, Germany).

### Cell culture

RAW 264.7 cells derived from RAW 264.7 macrophages (Invivogen, USA), containing a plasmid encoding a secreted embryonic alkaline phosphatase (SEAP) reporter under transcriptional control of an NF-κB-inducible promoter, were cultivated in Dulbecco Modified Eagle medium (DMEM) containing 4500 mg/L high glucose (1X) with 4.0 nM L-glutamine, without sodium pyruvate (Gibco, Thermo Fisher Scientific, Singapore) and supplemented with 10% fetal bovine serum (FBS; PAA) supplemented with 200 µg/mL Zeocin (Gibco, Thermo Fisher Scientific, Singapore), at 37^°^C in 5% CO2. For MyD88 rescue experiments and HIF-1α reporter assays cells were transduced with pCMV3-mMYD88 (Sino Biological) and pGL4.42[luc2P/HRE/Hygro] (Promega), respectively, using the Lipofectamine LTX with Plus Reagent (Thermo Fisher Scientific) following the manufacturer’s instructions at least 1 day before the assays were performed. HEK-Blue IL1R cells (Invivogen), which is a HEK 293 reporter cells for human and murine IL-1α & IL-1β cytokines, were cultured in DMEM (Gibco) supplemented with 10% (v/v) heat-inactivated fetal bovine serum, 100 U/mL penicillin, 100 µg/mL streptomycin, and 1x HEK-Blue selection (Invivogen) at 37°C in a humidified incubator with 5% CO₂. Cells were passaged when they reached 80% confluency. For passaging or harvesting cells for assays, cells were washed and incubated in PBS for 2 min at 37°C, followed by vigorous pipetting of PBS over the cell monolayer to detach the cells.

### Bone marrow-derived macrophages (BMDM) isolation and cultivation

BMDMs were harvested according to previously published protocol [97]. Briefly, femurs were harvested from 6 to 10 weeks old C57BL/6J mice (InVivos Pte. Ltd., Singapore). Following euthanasia, the legs were first sprayed with 70% ethanol before dissection and removing the skin and muscles. The bone was then placed in 70% ethanol and transferred into a sterile flow hood. Subsequently, the bone was cut using scissor and bone marrow was flushed out with cell culture media (containing DMEM with 10% FBS and supplemented with 100 U/mL penicillin and 100 µg/mL streptomycin (Gibco, Thermo Fisher Scientific, Singapore) using a 26G needle into a sterile falcon tube (Becton, Dickinson and Company, Franklin Lakes, NJ). Subsequently, cell suspension was centrifuged at 500 G for 5 min and the supernatant was removed. The cell pellet was then resuspended in 1X RBC Lysis Buffer (eBioscience, Thermo Fisher Scientific, Singapore) for lysis of erythrocytes. After 5 mins, PBS was added to inactivate the lysis buffer before centrifuging the cell suspension. Supernatant was discarded, and cell pellet was resuspended in cell culture media supplemented with 100 ng/mL macrophage colony-stimulating factor (M-CSF) (Biolegend, San Diego, USA), 100 U/mL penicillin and 100 µg/mL streptomycin. Cells were adjusted to 2 x 10^6^ cells/mL and seeded into T175 flask (Nunc, Thermo Fisher Scientific, China) and incubated at 37°C with 5% CO2. After 2-3 days, cells were washed, and fresh cell culture media supplemented with 100 ng/mL M-CSF was added. Macrophages are fully differentiated after 6 to 7 days, ready for experimental use.

### RAW 264.7 macrophage or BMDM infection

RAW 264.7 cells and BMDM were seeded into 96-well and 6-well microtiter plates (Nunc, Thermo Fisher Scientific, Singapore) with cell density of 1 x 10^5^ or 1 x 10^6^ cells/well respectively in antibiotic free media (DMEM with 10% heat inactivated FBS) and incubated overnight at 37°C with 5% CO2 when necessary. Following overnight incubation, the cells were washed once with PBS and fresh media was added. The SEAP reporter assay was established by empirically defining the minimal agonist LPS concentration that induced the maximum SEAP activity in the absence of cell death. Cells were stimulated using LPS purified from *E. coli* O111:B4 (Sigma-Aldrich, St Louis, MO) (100 ng/mL) as positive control, or media alone as a negative control. RAW 264.7 cells were infected with *E. faecalis* (at MOI of 100:1, 10:1 and 1:1) for 6 h with or without TLR agonists. Variations of this assay included lowering the pH to 6.4 with lactic acid, H3PO4 or HCl, and also neutralizing acidic supernatants to pH 7.8 with Tris. For co-infection experiments, RAW 264.7 cells were simultaneously infected with *E. coli* UTI89 (MOI of 0.125:1) and *E. faecalis* (MOI of 100:1). For killing assays, cells were infected at a multiplicity of infection (MOI) of 10 or 100 for up to 6 hours. Following infection, the medium was aspirated, and the cells were washed three times in PBS and incubated with gentamicin (150 μg/mL) (Sigma-Aldrich) and vancomycin (20 μg/mL) (Sigma-Aldrich) to kill extracellular bacteria in complete DMEM for up to 24 h to selectively kill extracellular bacteria. The antibiotic-containing medium was then removed, and the cells were washed three times in PBS before addition of 2% Triton X-100 (Sigma-Aldrich) PBS solution to lyse the cells for enumeration of the intracellular bacteria. Variations of this assay included lowering the pH to 6.4 with wither LA or HCl, pretreatment of mammalian cells for 30 min, before bacterial infection, with AZD3965 (10μM), PY60 (10 μM) or both followed by antibiotic treatment also containing these compounds.

### Fluorescent staining and microscopy

Cells were seeded at 2 × 10^5^ cells/well in a 24-well plate with 10 mm coverslips and allowed to attach overnight at 37 °C and 5% CO2. Infection and LPS treatment were performed for 6 h prior to fixation with 4% PFA at 4 °C for 15 min. Cells were then blocked for 1 h with PBS supplemented with 0.1% saponin and 2% bovine serum albumin (BSA). For antibody labelling, antibody solutions were diluted in PBS with 0.1% saponin at a 1:100 dilution for anti-p65 (Cell Signaling Technology), anti-STAT3 (Cell Signaling Technology) and anti-p-Yap (Cell Signaling Technology) and incubated overnight at 4°C. The following day, coverslips were washed 3 times in 1× PBS with 0.1% saponin and incubated with a 1:100 dilution of goat anti-Rabbit IgG (H+L) Alexa Fluor 568 secondary antibody (Thermo Fisher Scientific) for 1 h at room temperature.

For actin labeling, the phalloidin-Alexa Fluor 647 conjugate (Thermo Fisher Scientific) was diluted 1:40 in blocking solution and incubated for 1 h. Coverslips were then washed three times in PBS with 0.1% saponin. For nuclei staining, Hoechst 33342 was added in a dilution of 1:1000 for 5 min. Coverslips were then subjected to a final wash with PBS thrice. Finally, the coverslips were mounted with SlowFade Diamond Antifade (Thermo Fisher Scientific) and sealed. Confocal images were then acquired on a 63×/NA1.4, Plan Apochromat oil objective fitted onto an Elyra PS.1 with an LSM 780 confocal unit (Carl Zeiss), using the Zeiss Zen Black 2012 FP2 software suite. Laser power and gain were kept constant between the experiments. Z-stacked images were processed by using Zen 2.1 (Carl Zeiss). Acquired images were visually analyzed using ImageJ.

### Intracellular pH measurement

BCECF AM (Thermo Fisher Scientific) was used to measure intracellular pH as per manufacturer’s instructions. Cells were loaded with 1 mM BCECF AM for 30 min prior to infection. After 6 h of infection, fluorescence at excitation 440 nm and emission 535 nm was measured using a plate reader. To calibrate intracellular fluorescence ratio to pH, the CCCP technique was used [98]. Cells were first loaded with BCECF AM, then treated with 2 mM CCCP for 30 min following addition of pH-defined medium and fluorescence measurement.

### ChIP-qPCR and histone H3 acetylation assay

For each sample, chromatin immunoprecipitated DNA was captured using a STAT3 Rabbit mAb (D3Z2G; CST) and a high-sensitivity ChIP kit (Cat. #ab185913; Abcam) according to the manufacturer’s instructions and as inspired by [83] .The captured and purified DNA was prepared for ChIP-qPCR, and specific primers (**Table S5**) were designed for the sites that we identified as potential STAT3 binding sites near the *Myd88* transcriptional starting site (TSS). The prediction of STAT3 binding sites around the *Myd88* TSS was made using the Eukaryotic Promoter Database (https://epd.expasy.org/epd/). We identified six potential STAT3 binding sites near the *Myd88* TSS (-1147, -1065, -61, -23, 386 and 488). We named them region 1 (R1, - 1147 and -1065), region 2 (-61 and -23), and region 3 (386 and 488). For R1, we designed primers that covered from region -100 to 0; for R2 from -1030 to -1182 and R3 from 362 to 464 **(Table S5)**. qPCR was performed for each primer set. Amplification was performed with KAPA SYBR Fast (Kapa Biosystems) with specific primers **(Table S5)** and detected by StepOnePlus Real-Time PCR machine (Applied Biosystems). Relative quantification was performed using the delta-delta method [99] with calibrator primers that interrogate the *Gapdh* gene. Positive control was established for an RNA polymerase binding site for the *Gapdh* gene. To measure the levels of histone H3 acetylation of samples infected with *E. faecalis*, the histone H3 acetylation assay Kit (#ab115102, Abcam) was used as per the manufacturer’s instructions.

### Flow cytometry

To assess CD14 and TLR2 surface levels, flow cytometry was performed as described in [35, 95]. RAW 264.7 cells were either infected or treated with LPS (1 μg/mL) or LTA (10 μg/mL) alone or in combination for 30 min to 120 min before CD14 (LPS) and TLR2 (LTA) endocytoses were examined by flow cytometry. Cells (10^7^ per sample) were then incubated with 1 μL of Fc-blocker (anti-CD16/CD32 antibody; BioLegend) for 30 min, followed by incubation with a CD14 and TLR2 marker-conjugated with BV605 and PE, respectively, antibodies (BioLegend) (1:100 dilution) for 30 min at room temperature. Cells were then centrifuged at 500 *g* for 5 min at 4°C and washed in FACS buffer (2% FBS and 0.2 mM ethylenediaminetetraacetic acid (EDTA) in PBS (Gibco, Thermo Fisher Scientific). Following which, cells were analyzed using the BD LSRFortessa X-20 Cell Analyzer (Becton Dickinson). The samples were gated on side scatter area (SSC-A) versus forward scatter area (FSC-A) to select the RAW264.7 population. The RAW264.7 population was subsequently gated on side scatter width (SSC-W) versus SSC-A to exclude doublets. The resulting single-cell population was then assessed for CD14 and TLR2 fluorescent marker. Data was calculated relative to uninfected untreated cells which were set as 100% levels for the examined receptors.

For wound samples, flow cytometry was performed as described in [9]. Excised skin samples were placed in 1.5 mL Eppendorf tubes containing 2.5 U/ml liberase prepared in DMEM with 500 μg/mL of gentamicin and penicillin G (Sigma-Aldrich). The mixture was then transferred into 6-well plates and incubated for 1 h at 37 °C in a 5% CO2- humidified atmosphere with constant agitation. Dissociated cells were then passed through a 70 μm cell strainer to remove undigested tissues and spun down at 1350 rpm for 5 min at 4 °C. The enzymatic solution was then aspirated, and cells were blocked in 500 μL of FACS buffer (2% FBS and 0.2 mM ethylenediaminetetraacetic acid (EDTA) in PBS (Gibco, Thermo Fisher Scientific). 10^7^ cells per sample were then incubated with 1 μL of Fc-blocker (anti-CD16/CD32 antibody, Biolegend) for 30 min, followed by incubation with an antimouse CD45, CD11b, and F4/80 (macrophages) plus MyD88 (ProteinTech) and p-ERK (Biolegend) markers conjugated with PE-Cy5, PE-Cy7, BV605, FITC and APC, respectively, antibodies (1:100 dilution) for 30 min at room temperature. Cells were then centrifuged at 500*g* for 5 min at 4 °C and washed in FACS buffer. Cells were fixed in 4% PFA for 15 min at 4 °C, before a final wash in FACS buffer and final resuspension in this buffer. Following which, cells were analyzed using a BD LSRFortessa X-20 Cell Analyzer (Becton Dickinson). Samples were first gated on side scatter area (SSC-A) versus forward scatter area (FSC-A) to identify the wound cell population. This population was then gated on forward scatter height (FSC-H) versus FSC-A to exclude doublets. The resulting single-cell population was analyzed for CD45 expression to identify immune cells, followed by assessment of CD11b and F4/80 fluorescence to define the macrophage population. Compensation was done using AbC Total Antibody Compensation Bead Kit (Thermo Fisher Scientific) as per manufacturer’s instructions.

### siRNA transfection

Silencer select pre-designed siRNA (Thermo Fisher Scientific) against *Slc16a1 (Mct- 1)*, *Hcar1 (Gpr81), Gpr65, Gapdh* and NT siRNA were used. RAW264.7 cells were seeded at 40% density the day before transfection in 6-well plates, and siRNA transfections were performed with Silencer™ siRNA Transfection II Kit (Invitrogen) according to the manufacturer’s instructions. Cells were used for experiments after 48 h and also harvested at the same time point after transfection to verify successful gene silencing by RT-PCR.

### Collection of bacteria cell-free culture supernatants

Bacteria were grown in cell culture media for 6 h, and bacteria-free culture supernatants were collected after centrifugation (6,000 *g*) followed by filtration (using a 0.2 µm syringe filter). Alternatively, supernatants were collected after infecting macrophages with bacteria at various MOI and then filtered by using a 0.2 µm syringe filter. Sterility of bacteria-free culture supernatants were verified by the absence of viable bacteria when plated on BHI agar.

### NF-κB reporter assay

Post-infection, 20 µL of supernatant was added to 180 μL of QUANTI-Blue reagent (Invivogen) and incubated overnight at 37 ^°^C. SEAP levels were determined at 640 nm using a TECAN M200 microplate reader. All experiments were performed in biological triplicate.

### IL-1 reporter assay

After RAW264.7 infection, bacteria were pelleted and supernatants were filter-sterilized. 20 µl of bacterial supernatant filtrate were added to 50,000 HEK-Blue IL-1R cells (Invivogen) in 180 µl DMEM supplemented with 10% FBS and 100 U/mL Penicillin-Streptomycin in a 96-well plate. As negative control, DMEM was used instead of bacterial supernatants. DMEM supplemented with human mature IL-1β (100 pg/mL) served as a positive control. Some RAW264.7 cells were pre-treated with HIF- 1α inhibitor LW6 (10 μM) or HIF-1α activator O-phenanthroline (5 μM) for 30 min prior to the beginning of the assay. Samples were incubated for 6 h at 37°C. Next, 20 µl of HEK-Blue IL-1R cell supernatants were transferred to 180 µl of QUANTI-Blue Solution (invivogen) in a 96-well plate and incubated at 37°C for 1 h, followed by absorbance at 640 nm in a plate reader (TECAN).

### Cell viability assay

Simultaneously with supernatant collection for SEAP determination, culture supernatants were collected from each well to measure lactate dehydrogenase (LDH) release, using an LDH cytotoxicity assay (Clontech) according to manufacturer’s instructions. Background LDH activity was determined using mock (PBS) treated RAW 264.7 cells. Maximal LDH activity was determined by lysing cells with 1% Triton-X. Each condition was assayed in triplicate. Percentage cytotoxicity was calculated as follows: (sample absorbance-background absorbance)/(maximal absorbance- background absorbance) x 100.

### Transposon library screen

The transposon library was kindly provided by Professor Dunny Gary [100]. To screen the library, the mutants were first inoculated into DMEM for 24 h and 20 μL of the bacteria suspension will be transferred to previously seeded RAW 264.7 cells with 180 μL fresh DMEM in the presence of 100 ng/mL of LPS in 96 well microtitre plate. Macrophages were stimulated for 6 h before the supernatant was collected and tested with SEAP assay to determine the NF-κB activity. LPS-only group was used as positive control for reporter activity and signals higher than the means of LPS-only group minus 1 standard deviation was considered positive hits and non-suppressing mutants. Growth kinetics of the selected mutants were performed to exclude the mutants that grow poorly in DMEM. In addition, pictures of the overnight cultures were taken to record the acidifying ability of each mutant by phenol red, a color indicator in DMEM.

Mutants that grew similarly to *E. faecalis* WT further underwent a secondary screening, where they were normalized to MOI 100 and infected the RAW 264.7 cells in the presence of LPS. For the secondary screen, WT was used as control and signals higher than means of WT group plus 2 standard deviations were considered weak-suppressing mutants and validated hits. To measure the acidification ability of the validated hits, 10^8^ CFU of bacteria were inoculated into 5 mL of DMEM for indicated time, and the pH value was determined by pH meter.

### Lactate and cAMP levels measurement

Bacteria were grown in DMEM supplemented with 10% FBS for 6 h in the absence of macrophages before collecting the supernatant and storing it at -80°C until assessment by the L-Lactate Assay Kit (Abcam), according to manufacturer’s recommendations. All samples were assessed using the same kit lot and at the same time to avoid inter-assay variability. Alternatively, blood gas analysis using abl90 flex series analyser (Radiometer Medical) or or QuantiQuik L-Lactic acid quick test strips (BioAssay Systems) were used to quantify lactate concentration. Intracellular levels of cAMP were assessed using cAMP Direct Immunoassay Kit (Abcam) according to the manufacturer’s instructions.

### IL-6, IL-1β and IL-10 quantification

Wound lysates from 24 hpi and supernatants collected from RAW 264.7 cells 6 hpi were stored at -80 °C until assessment by the ELISA MAX Deluxe Set Mouse for IL- 6, IL-1β and IL-10 assay kit (Biolegend), according to manufacturer’s recommendations. All samples were assessed using the same kit lot and at the same time to avoid inter-assay variability.

### Cytokine Luminex xMAP analysis

Supernatants were collected from RAW 264.7 cells 6 hpi and stored at -80°C until assessment by the Bio-Plex ProTM mouse cytokine 23-plex assay kit (Bio-Rad Laboratories), according to manufacturer’s recommendations. All samples were assessed using the same kit lot and at the same time to avoid inter-assay variability.

### Mouse model of wound infection

Bacterial cultures, prepared as described above, were used to infect wounds in 7 to 8 weeks old, 20 to 23 g female C57BL/6J mice (InVivos). Wounds were prepared as previously described [57]. Briefly, mice were anesthetized with isoflurane (3%) and the dorsal hair was trimmed. After trimming, remaining fine hairs were removed by first applying Nair Cream (Church and Dwight) and shaving using a scalpel. Next, the skin was first disinfected using 70% ethanol and a 6 mm biopsy punch (Integra Miltex) was used to create a fill-thickness wound and inoculated with bacteria subsequently. Inoculum volumes of 10 µL, containing a bacterial suspension of either single or polymicrobial species prepared in PBS: (i) 10^2^ CFU of *E. coli* strain UTI89 with 10^6^ CFU *E. faecalis* WT, and (ii) 10^2^ CFU of *E. coli* strain UTI89 with 10^6^ CFU *E. faecalis* Ldh. Single species controls (10^2^ CFU of *E. coli* strain UTI89, 10^6^ CFU *E. faecalis* OG1RF, or 10^6^ CFU *E. faecalis* Ldh) were performed alongside polymicrobial infections. The wound site was then sealed with a sterile transparent dressing (Tegaderm 3M, St Paul Minnesota, USA). After 24 h, animals were euthanized by carbon dioxide inhalation and cervical dislocation and a 1 cm by 1 cm squared piece of skin surrounding the wound was excised and homogenized in 1 mL PBS for CFU enumeration by serial dilution on MacConkey agar (Becton, Dickinson and Company) or BHI agar supplemented with 10 µg/mL colistin (Sigma-Aldrich) and 10 µg/mL nalidixic acid (BHI-CNA) (Sigma-Aldrich) to isolate *E. coli* or *E. faecalis*, respectively.

### RNA extraction

After 6 hpi with *E. faecalis* at various MOI in the presence or absence of LPS (100 ng/mL), cells were washed once with the tissue culture grade 1X PBS (Gibco, Thermo Fisher Scientific) prior to RNA extraction. 1 mL of Trizol (Invitrogen, Thermo Fisher) per well and was stored at -80°C until RNA extraction using Direct-zol RNA Miniprep Plus kit (Zymo Research) according to manufacturer’s protocol. Briefly, 200 μL of chloroform (Sigma Aldrich) was first added to 1 mL of Trizol extracted samples, mixed vigorously for 2 min and allowed to stand at room temperature for minimum 10 min. The samples were centrifuged at 12,000 *g* for 15 min at 4°C and the top layer (aqueous phase) was transferred to 500 μL of ethanol (Fisher Scientific, Canada) before loading into the Zymo-Spin IIICG Column. Subsequently, RNA extraction was performed according to manufacturer’s protocol for Direct-zol RNA Miniprep Plus kit. Next, extracted RNA samples were further cleaned and concentrated using RNA Clean & Concentrator-5 (Zymo Research) following the manufacturer’s protocol.

RNA samples were first screened using TapeStation (Aligent Technologies) to obtain RIN value and concentration. Qubit analysis was also performed to measure the DNA and RNA concentration using Qubit dsDNA HS Assay Kit (Invitrogen, Thermo Fisher Scientific) and Qubit RNA BR Assay Kit (Invitrogen, Thermo Fisher Scientific), before storage at -80°C until sequencing using an Illumina Hiseq®2500 v.2 (Illumina), 150 bp paired-end.

### qRT-PCR

To quantify the levels of RNA transcripts, total RNA was extracted from noninfected or WT-infected RAW264.7 cells in presence or absence of LPS (100 ng/mL) with the RNeasy MinElute Kit (QIAGEN) and reverse-transcribed using SuperScript III First-Strand Synthesis SuperMix (Thermo Fisher Scientific), followed by amplification with KAPA SYBR Fast (Kapa Biosystems) and detected by StepOnePlus Real-Time PCR machine (Applied Biosystems). Relative quantification of gene expression was performed using the comparative CT method [101], the CT values were normalized with the housekeeping gene glyceraldehyde phosphate dehydrogenase (GAPDH) for comparison.

### Protein extraction and immunoblot

Following infection with *E. faecalis* as described previously, proteins were harvested using 2x Laemmli buffer (Bio-rad, Hercules) containing 50 mM dithiothreitol (Sigma Aldrich) and denatured at 99°C for 10 min. 30 μL of protein samples were loaded and subjected to Sodium Dodecyl Sulfate-Polyacrylamide Gel Electrophoresis (SDS-PAGE) using NuPAGE 4-12% Bis-Tris Gel (Invitrogen, Thermo Fisher Scientific) in 1 x NuPAGE MOPS SDS Running Buffer (Invitrogen, Thermo Fisher Scientific). The protein samples were transferred to polyvinylidene fluoride (PVDF) membrane using iBlot® Gel Transfer Device (Invitrogen, Thermo Fisher Scientific, USA) as per manufacturer’s instructions. The PVDF membrane was then blocked using 5% BSA in Tris-buffered saline (20 mM Tris-HCl pH 7.4, 150 mM NaCl) containing 0.1% Tween- 20 (Sigma Aldrich) at room temperature for 1 h. Subsequently, primary antibodies such as MyD88, TAK1, phospho-TAK1, IKKα, IKKβ, phospho-IKKα/β, IκBα, phospho- IκBα, p65, phospho-p65, SAPK/JNK, p38 MAPK, phospho-p38, p44/p42 MAPK (ERK1/2), phospho-p44/p42 MAPK, c-Jun, phospho-c-Jun, c-Fos, phospho-cFos and GADPH (Cell Signalling Technology) were used for probing overnight at 4 °C. The membranes were washed with Tris-buffered saline and incubated with 1:3000 diluted anti-rabbit IgG antibodies conjugated with horseradish peroxidase (Invitrogen, Thermo Fisher Scientific, USA) at room temperature for 1 h. Following which, membranes were washed, and proteins were detected using SuperSignal West Femto Maximum Sensitivity Substrate (Invitrogen, Thermo Fisher Scientific) and autoradiography films (Kodak).

### Subcellular Protein Fractionation

The subcellular protein fractionation kit for cultured cells (Thermo Fisher Scientific) was used as per manufacturer’s instructions. Briefly, after infection cells were washed and harvested, then gently plasma membrane permeabilized to collect the cytosolic proteins. Cellular membranes were solubilized to remove membrane-associated proteins while keeping nuclei intact. Nuclear proteins were then extracted using nuclear buffer. Collected samples were then used for immunoblot.

### *In vitro* MEK activity assay

*In vitro* MEK activity assay was performed using the MEK Activity Assay Kit (Sigma, code CS0490) instructions with some modifications. MEK was immunoprecipitated from samples with anti-MEK antibody (Thermo Fisher Scientific) and Protein A/G magnetic beads (Abcam). The immunoprecipitated MEK was used to phosphorylate the non-phosphorylated ERK substrate from cells that were treated with Selumetinib (1 μM) for 6h and further depleted of MEK via IP. The level of ERK phosphorylation was detected by immunoblotting using anti-p-ERK.

### Transcriptomics analysis

Sequencing data are available at NCBI’s BioProject (accession no. GSE188916). The sequencing was performed at the Singapore Centre for Environmental Life Sciences Engineering High Throughput Sequencing Unit, NTU, Singapore. Following quality control (FastQC v0.11.9), trimming (cutadapt v4.2) and pre-processing using TrimGalore (v0.6.6), the reads were mapped to the Gencode Mouse Release M32 (GRCm39) using Salmon (v.1.10.0). Differentially expressed gene analysis was conducted in DESeq2 (v1.38.3) from raw counts. For over-representation and gene set enrichment analysis (GSEA), we utilised clusterProfiler (v 4.6.0), enrichplot (v 1.18.3) and DOSE (v 3.24.2) packages for Gene Ontology (GO) analyses. The workflow can be found at https://github.com/cenk-celik/daSilva_et_al.

### Statistical Analysis

Statistical analysis was performed utilizing GraphPad Prism V6 and V10.4.1 software (GraphPad Software). One-way ANOVA (Analysis of Variance) were used to compare between median and the different conditions. Statistical significance for multiplex immunoassay was determined using Kruskal-Wallis statistical analysis. CFU titers were compared using the non-parametric Mann-Whitney U test. *P*-values less than 0.05 were deemed significant. Unless otherwise stated, values represented means ± SEM derived from at least 3 independent experiments.

### Ethics Statement

All animal experiments were performed with approval from the Institutional Animal Care and Use Committee (IACUC) in Nanyang Technological University (NTU), School of Biological Sciences under protocols ARF-SBS/NIE-A18061, A19061 and A24066.

## Author contributions

R.A.G.d.S., B.Y.Q.T., and K.A.K. designed the experiments. R.A.G.d.S., B.Y.Q.T., P.H.N.K., H.A., A.Z.C.T., and K.K.L.C. performed experiments. R.A.G.d.S., B.Y.Q.T., P.H.N.K., C.C., M.H.I., and G.H. analyzed data. R.A.G.d.S., B.Y.Q.T. and K.A.K. interpreted the results. B.Y.Q.T. and K.A.K. conceived the project. G.T., J.C., and K.A.K. supervised, acquired funding, and provided resources. R.A.G.d.S., B.Y.Q.T. and K.A.K. wrote the manuscript. All authors reviewed and approved the final manuscript.

## Supplementary Figures

**Figure S1.**
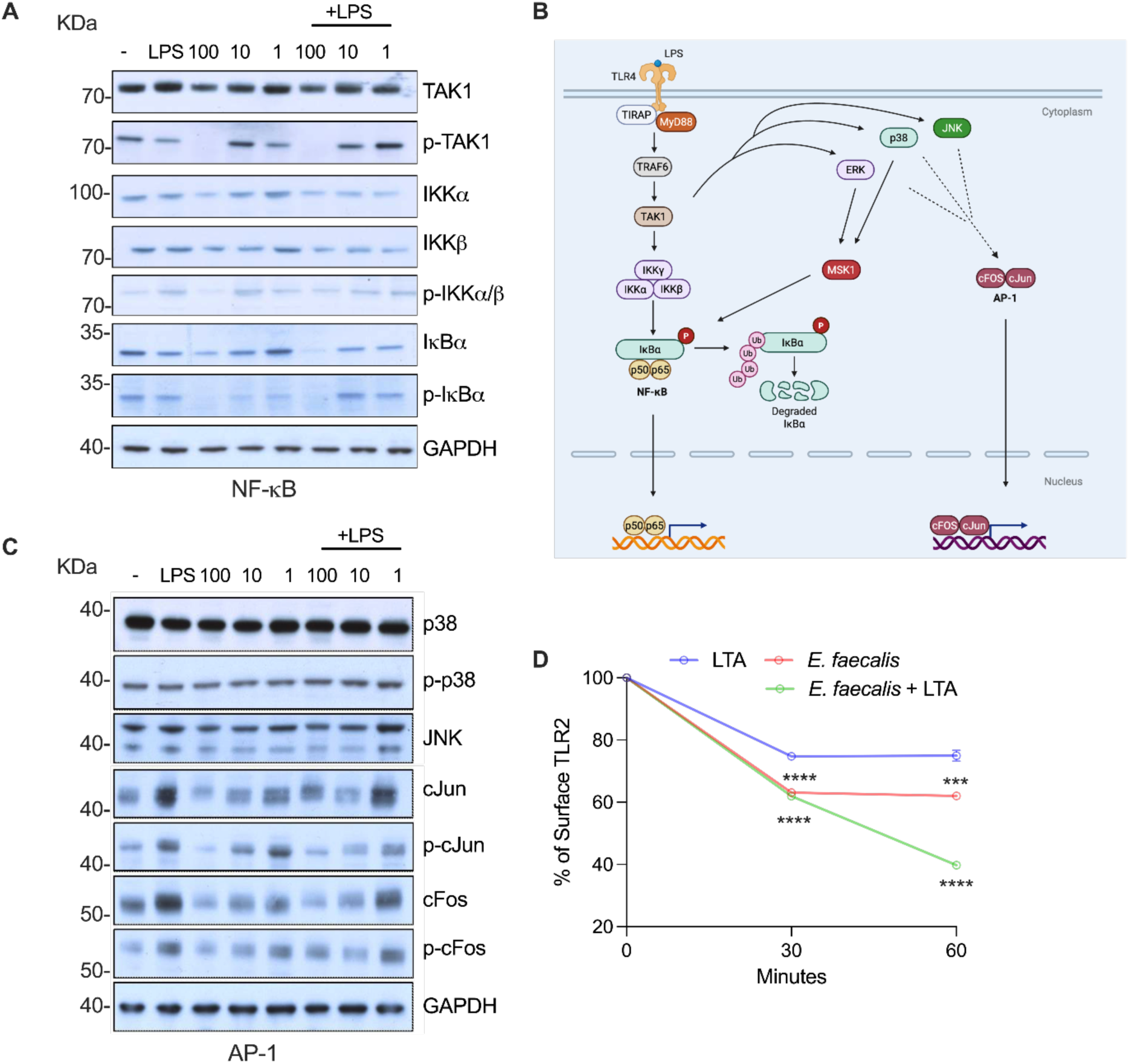
*E. faecalis* infection modulates macrophage response to bacteria by affecting the production and phosphorylation status of different proteins of the NF-κB pathway. **(A)** Immunoblot performed on protein extracted from RAW 264.7 macrophages infected with WT *E. faecalis* at the indicated MOI (100, 10, 1) with or without LPS (100 ng/mL) for 6 h and probed with antibodies specific for TAK1, phospho-TAK1, IKKα, IKKβ, phospho-IKKα/β, IκBα, phospho-IκBα, p65, phospho-p65 and GAPDH. M represents uninfected macrophages. GAPDH was used as an internal control for loading control. **(B)** TLR-NF-κB signaling pathway and cross-talk between NF-κB and AP-1 are shown. **(C)** Immunoblot performed on protein extracted from RAW 264.7 macrophages infected with WT *E. faecalis* at the indicated MOIs (100, 10, 1) with or without LPS for 6 h and probed with antibodies specific for p38, phospho-p38, JNK, ERK, c-Jun, phospho c-Jun, c-Fos, phospho c-Fos and GAPDH. **(D)** RAW 264.7 cells were either infected or treated with LTA (10 μg/mL) for the times indicated before TLR2 endocytosis was examined by flow cytometry. Data (mean ± SEM) are a summary of three independent experiments and calculated relative to uninfected untreated cells which were set as 100% levels for the examined receptor. Statistical analysis was performed using ordinary one-way ANOVA followed by Dunnett’s multiple comparison test ***p<0.001 and ****p<0.0001.

**Figure S2.**
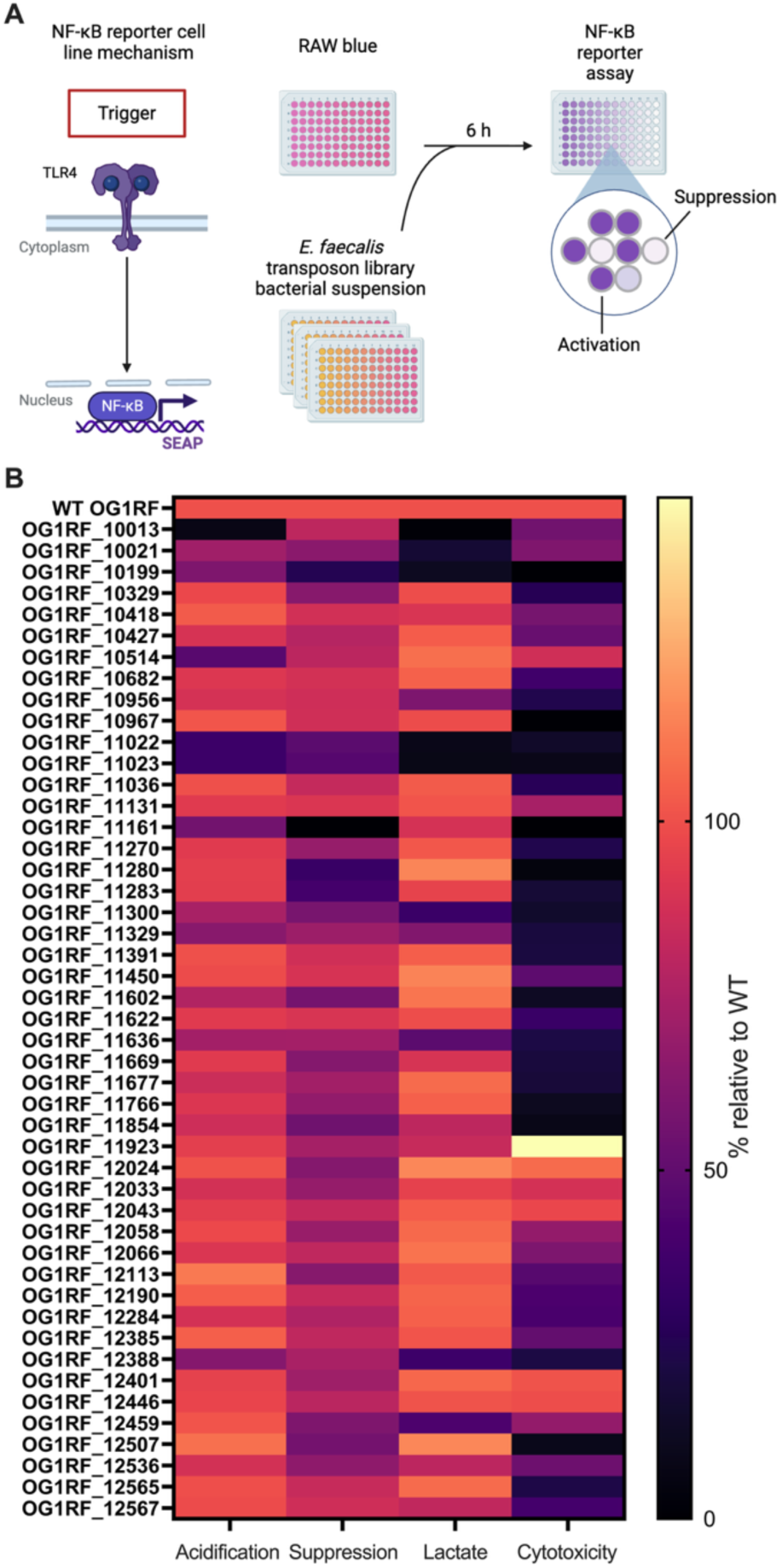
Methodology for transposon library screen to identify bacterial factor(s) important in macrophage suppression. **(A)** Screening workflow: all mutants were inoculated in DMEM + 10% FBS for 24 h before 20 μL of the bacteria suspension was transferred to RAW 264.7 cell culture. The macrophages were stimulated in the presence of LPS for 6 h and supernatant was collected for SEAP assay. Mutants showed significant higher NF-κB activity were identified as weak-suppressing mutants. **(B)** Bacterial genes identified in a transposon library screen for genes important in macrophage suppression at MOI 100. The heat map shows the degree of NF-κB suppression, medium acidification, lactate production, and cytotoxicity in RAW 264.7 macrophages infected at MOI 100 at 6 hpi relative to WT levels set to 100%.

**Figure S3.**
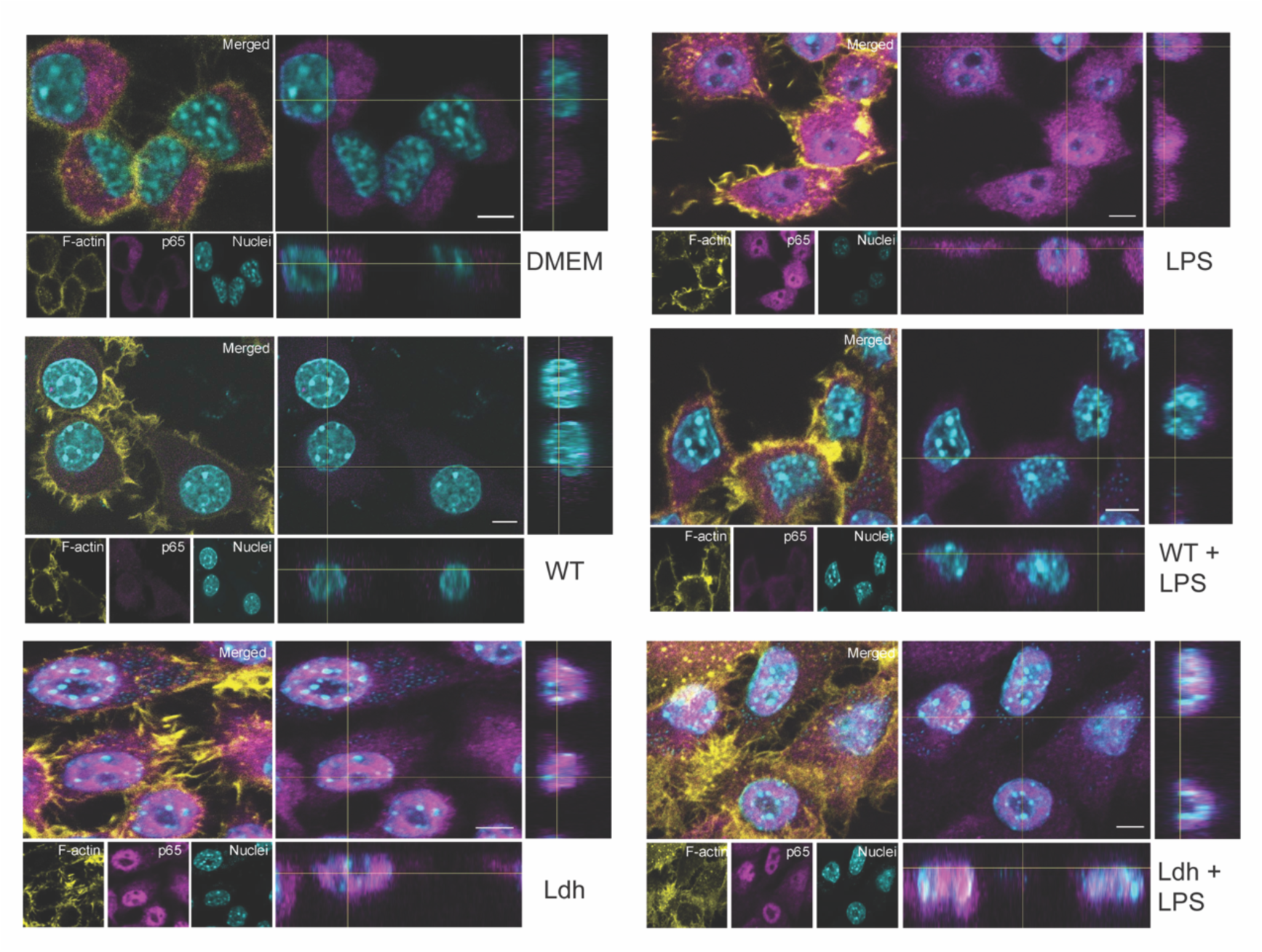
***E. faecalis* infection prevents p65 nuclear translocation.** CLSM orthogonal view of RAW 264.7 macrophages at 6 hpi with WT or Ldh strains in presence or absence of LPS. Magenta, p65; cyan, dsDNA stained with Hoechst 33342; yellow, F-actin stained with phalloidin. Images are representative of 2 independent experiments. Scale bar: 5 μm.

**Figure S4.**
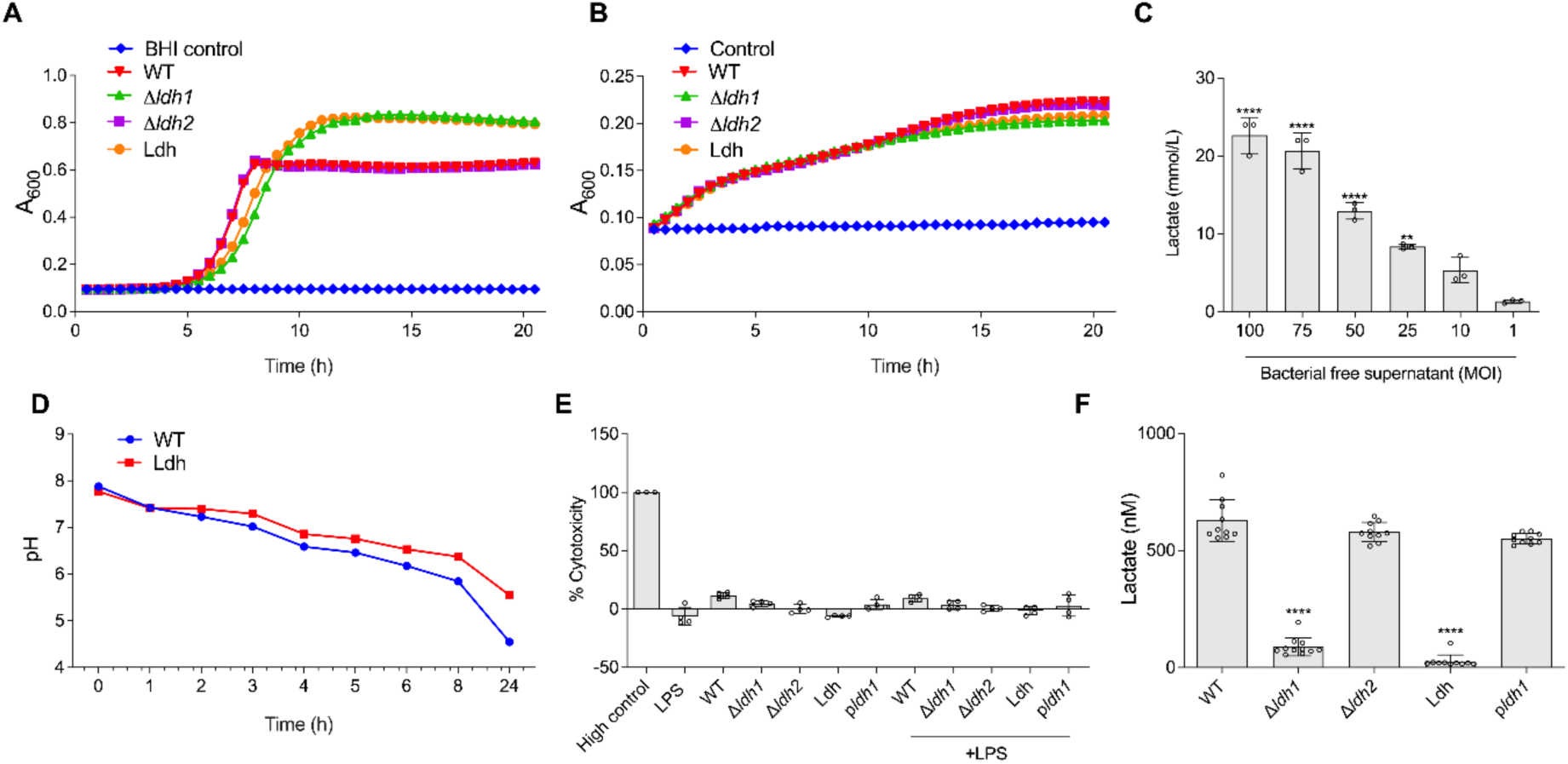
LDH mutant does not have growth defects and acidifies the environment weakly. Growth curve for *E. faecalis* strains grown overnight in either **(A)** BHI or **(B)** cell culture media supplemented with 10% FBS. **(C)** Lactate present in bacterial-free supernatant was quantified via blood gas analysis. **(D)** pH for *E. faecalis* strains grown in cell culture media measured over a time course. **(E)** RAW 264.7 macrophages infected with WT OG1RF or Ldh mutant supplemented with lactic acid at various concentration for 6 h prior to measurement of cytotoxicity. **(F)** Lactate concentration was measured using 6 hpi bacterial free supernatant. Statistical analysis was performed using one-way ANOVA with Tukey’s multiple comparison test where *p*<0.0001 among all the different *E. faecalis* strains as compared to WT.

**Figure S5.**
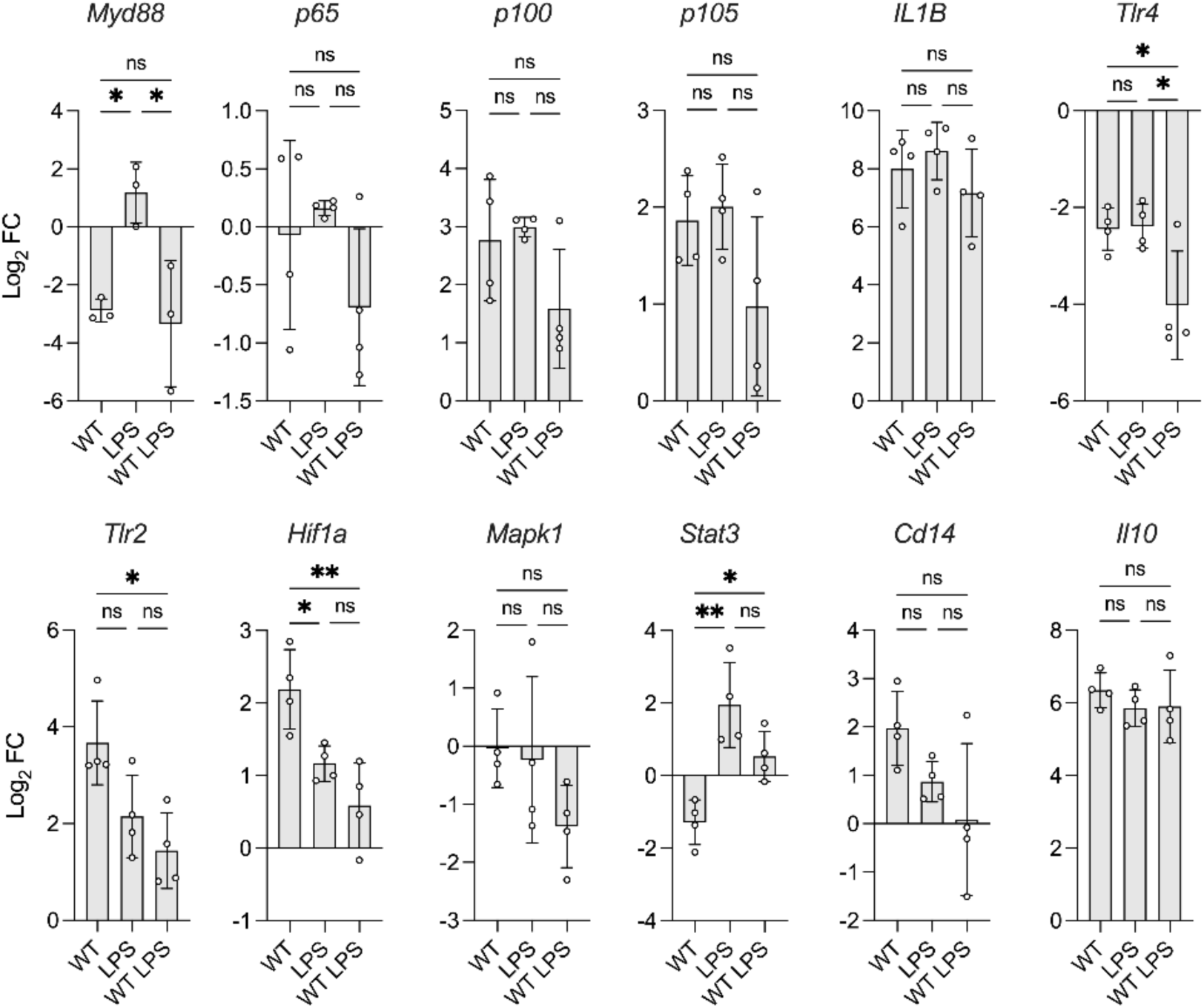
***E. faecalis* infection affect transcription of immune response genes in BMDM.** qRT-PCR analysis of NF-κB-related genes transcript level in RAW 264.7 cells that were infected with WT *E. faecalis* for 6 h in the presence or the absence of LPS. Each dot represents one biological replicate. Statistical analysis was performed using ordinary one-way ANOVA, followed by Tukey’s multiple comparison test, NS p > 0.05; *p≤0.05; and **p<0.01.

**Figure S6.**
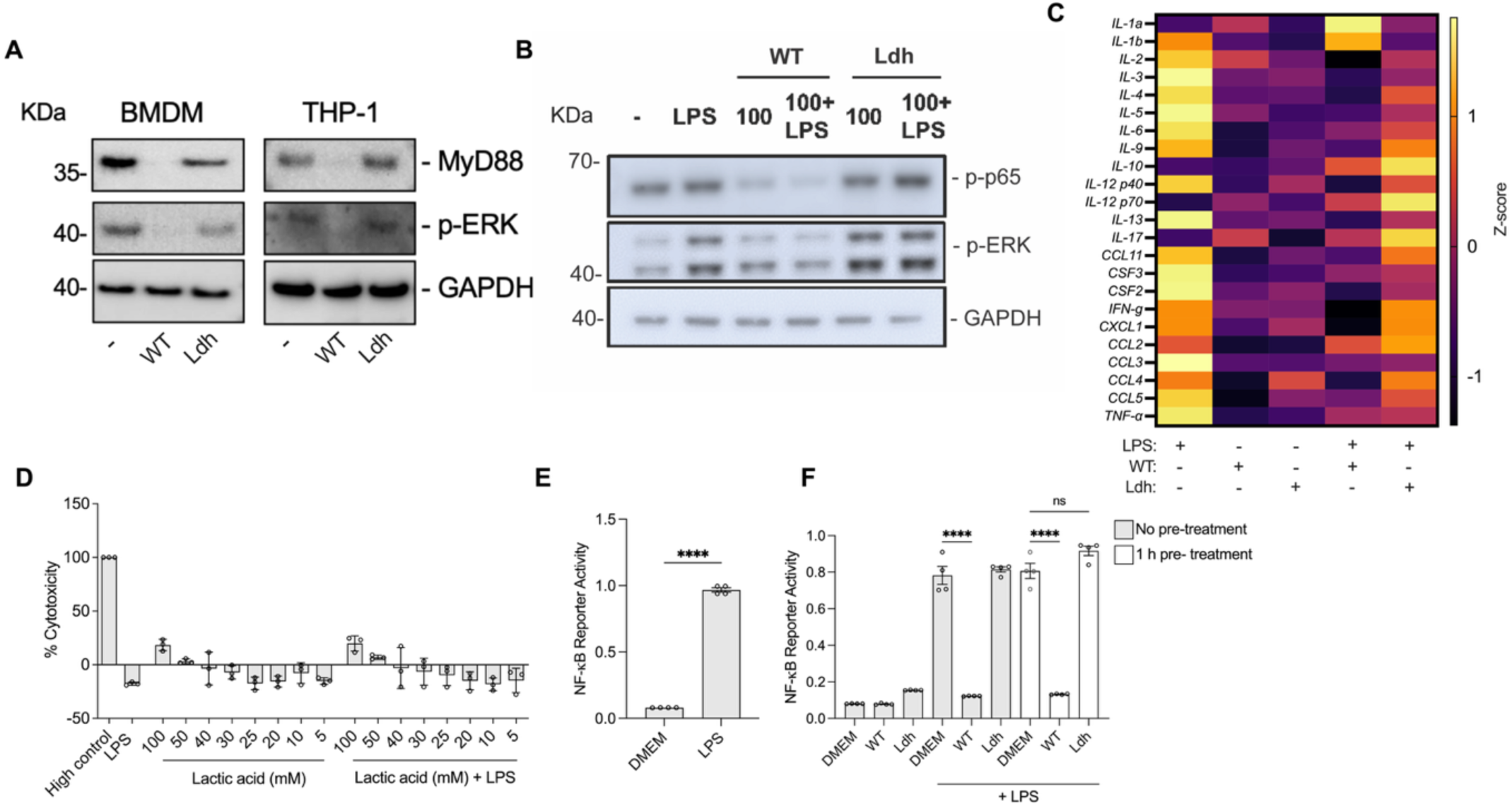
*E. faecalis*-derived lactic acid suppresses NF-κB activation in LPS pre-treated cells. Immunoblot performed on proteins extracted from BMDMs, THP-1s and RAW 264.7 macrophages infected with *E. faecalis* MOI 100 for 6 h and probed with antibodies specific for **(A)** MyD88, p-ERK and GAPDH, and **(B)** p-p65, p-ERK and GAPDH. Samples in **(B)** are with or without LPS. Data (mean ± SEM) are a summary of at least two independent experiments. **(C)** Heatmap showing log2-transformed, row-scaled fold changes in cytokine, chemokine, and growth factor expression relative to uninfected controls, detected in filtered supernatants at 6 hpi from WT *E. faecalis* and the Ldh mutant. The scale bar represents z-scores (mean = 0, SD = 1). **(D)** RAW 264.7 macrophages were incubated with cell culture media supplemented with lactate at various concentrations for 6 h prior to measurement of cytotoxicity. **(E)** RAW 264.7 cells were pre-treated for 1 h with LPS prior to measurement of NF-κB-driven SEAP reporter activity. **(F)** RAW 264.7 cells infected with WT *E. faecalis* or Ldh mutant for 6 h prior to measurement of NF-κB-driven SEAP reporter activity were either pre-treated or not with LPS for 1 h prior to infection. Data (mean ± SEM) are a summary of at least three independent experiments. Statistical analysis was performed using one-way ANOVA with Tukey’s multiple comparison test where NS, p>0.05 and ****p<0.0001.

**Figure S7.**
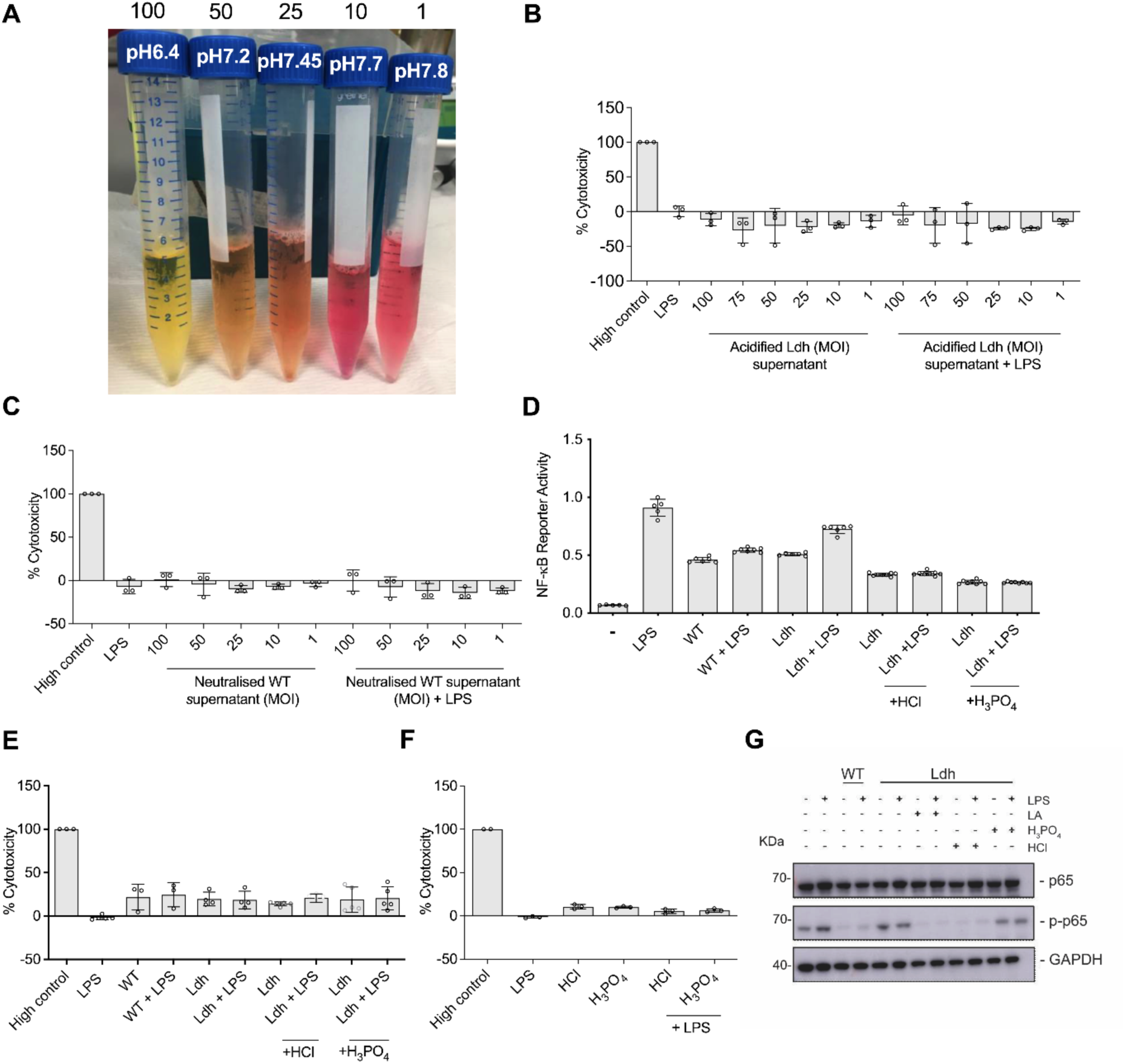
pH adjusted medium does not affect macrophage viability. **(A)** Picture of bacterial-free supernatant at indicated MOI. The color indicator in the cell culture medium turns yellow upon acidification. RAW 264.7 macrophages were incubated with **(B)** acidified bacterial-free Ldh supernatant, **(C)** neutralized bacterial-free WT supernatant, **(D-E)** WT *E. faecalis* or Ldh deletion mutant treated concurrently with LPS at specific MOI for 6 h, and **(F)** acidified cell media in the absence of bacteria, prior to measurement of NF-κB activity or cytotoxicity. **(B-C, E-F)** Triton-X treatment served as a positive control (high control) for cell death. **(D-E)** RAW 264.7 macrophages incubated with WT or Ldh mutant were also supplemented with different acids to acidify the pH for the duration of the experiment prior to measurement of NF-κB-driven SEAP reporter activity and LDH activity. **(B-F)** Data (mean ± SEM) are a summary of at least three independent experiments. **(G)** Immunoblots were performed on protein extracted from RAW 264.7 macrophages infected with WT or Ldh that were incubated in acidified cell culture media in the presence or absence of LPS for 6 h and probed with antibodies specific for p65, p-p65, and GAPDH. Cell culture media was acidified to pH 6.4 with lactic acid and with the inorganic acids H3PO4 and HCl. Data shown are representative of at least 3 independent experiments.

**Figure S8.**
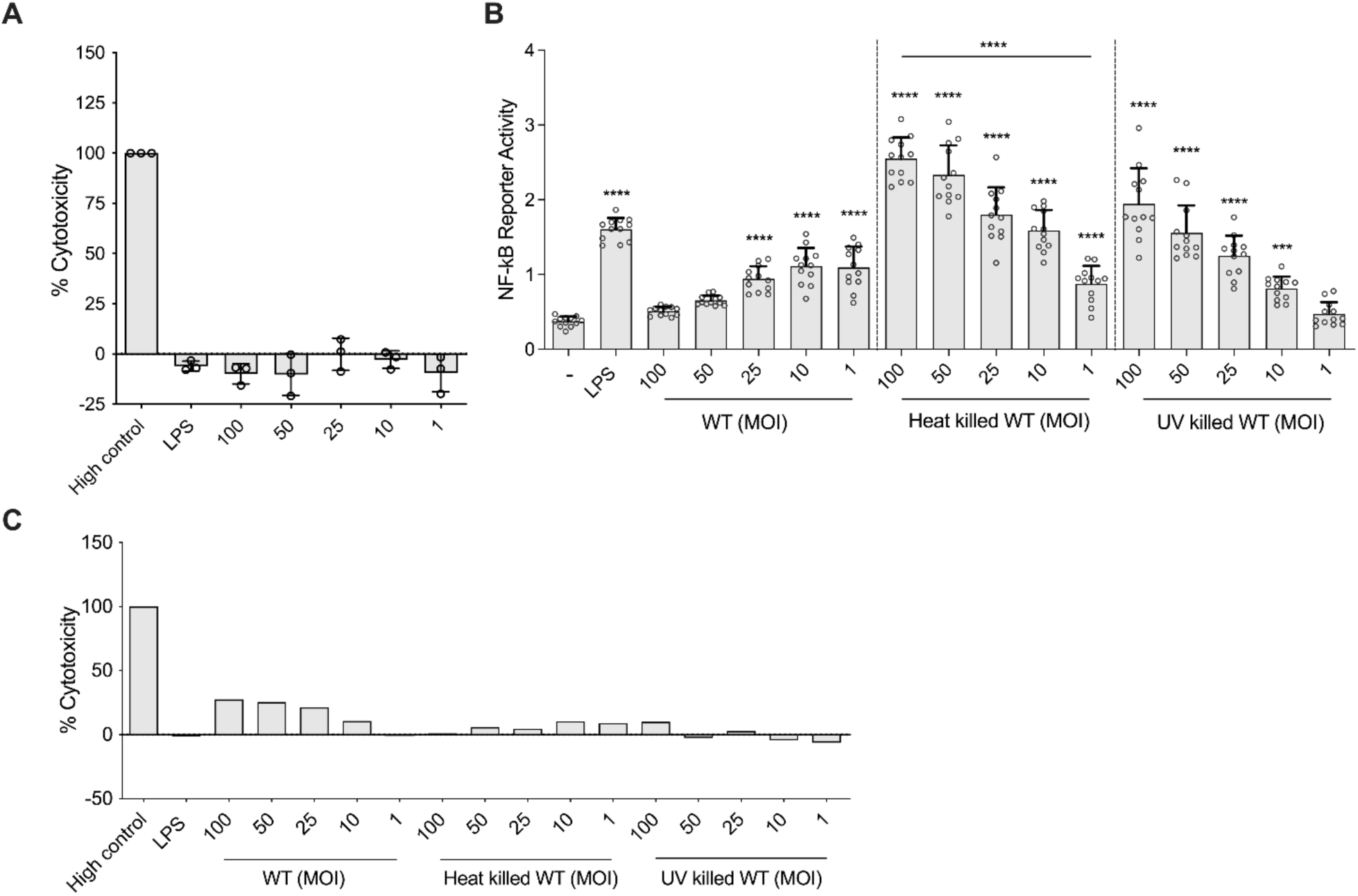
Intracellular, heat or UV killed *E. faecalis* do not contribute to NF-κB suppression in macrophages. RAW 264.7 macrophages were incubated with *E. faecalis* WT (MOI 100, 50, 25, 10, 1) for 3 h and treated with 500 µg/mL of gentamicin with penicillin G overnight at 37°C **(A)** and were incubated with live, heat killed or UV killed WT (MOI 100, 50, 25, 10, 1) for 6 h **(B)** prior to measurement of NF-κB-driven SEAP reporter activity and **(A and C)** measurement of cytotoxicity. **(A-C)** Data (mean ± SEM) are a summary of at least three independent experiments. Statistical analysis was performed using one-way ANOVA with Tukey’s multiple with comparison where ***p<0.001 and ****p<0.0001 as compared to media control.

**Figure S9.**
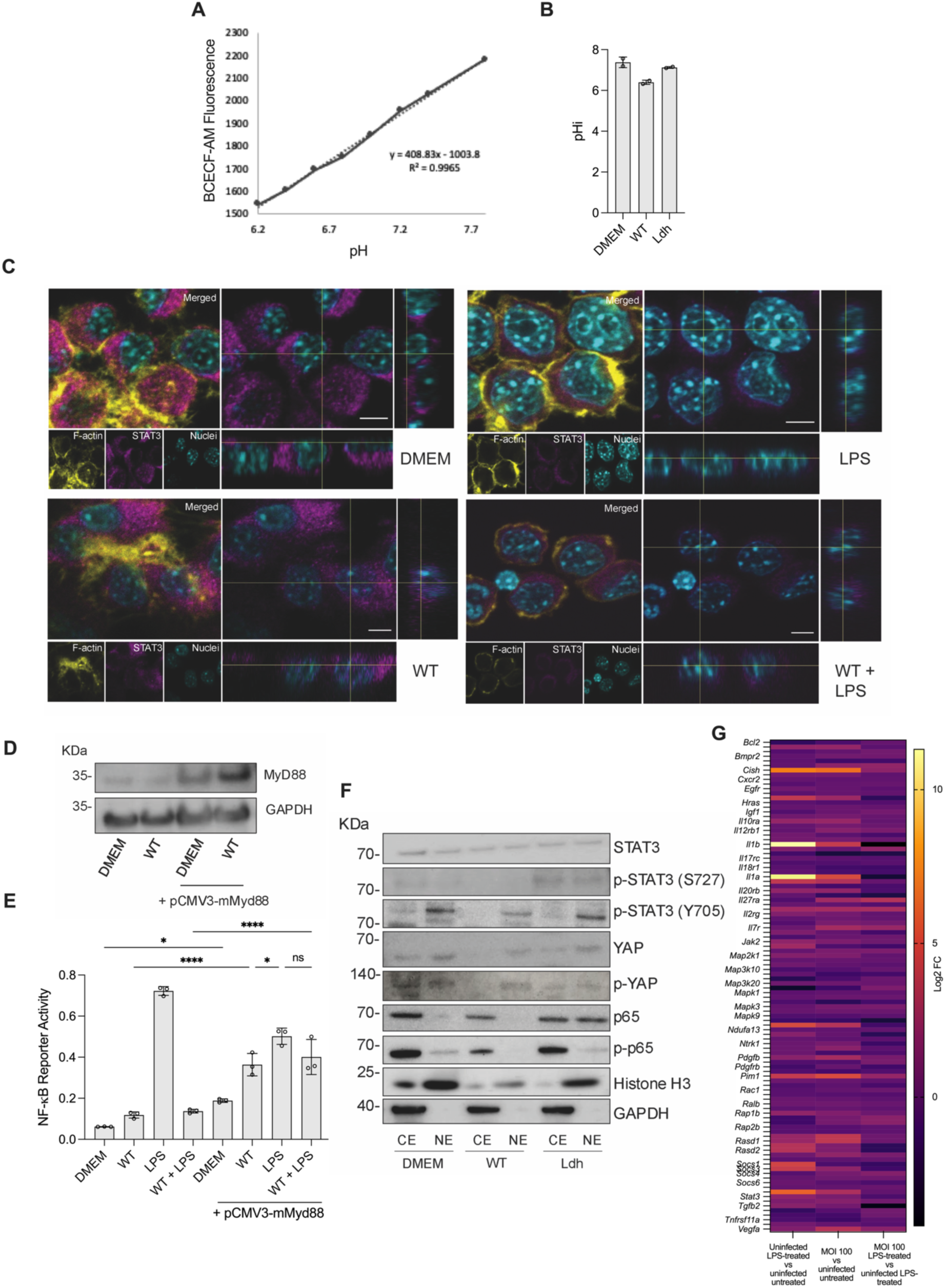
Low pH during *E. faecalis* infection prevents ERK phosphorylation. **(A)** BCECF-AM calibration curve used in the BCECF-AM experiments. **(B)** pHi in non-infected, WT or Ldh infected RAW264.7 6 hpi using BCECF-AM. **(C)** CLSM orthogonal view of RAW264.7 macrophages at 6 hpi with WT in the presence or absence of LPS. Magenta, STAT3; cyan, dsDNA stained with Hoechst 33342; yellow, F-actin stained with phalloidin. Images are representative of 2 independent experiments. Scale bar: 5 μm. **(D-E)** MyD88 rescue experiment using the mammalian expression vector pCMV3- mMyd88. A representative MyD88 immunoblot of successfully transduced RAW264.7 cells that were uninfected or infected with *E. faecalis* is shown **(D)**. **(E)** Cells were transduced 24 h prior to performing NF-κB reporter assay. Data (mean ± SEM) represent three independent experiments. Statistical analysis was performed using ordinary one-way ANOVA followed by Tukey’s multiple comparison test, ns, p>0.05; *p≤0.05; and ****p<0.0001. **(F)** Immunoblot was performed on proteins from RAW 264.7 macrophages subcellular fractions after WT or Ldh mutant infection for 6 h and probed with antibodies raised against STAT3, p-STAT3 (S727), p-STAT3 (Y705), YAP, p-YAP, p65, p-p65, histone H3 and GAPDH. CE and NE stand for cellular and nuclear extract, respectively. Immunoblot images represent two independent experiments **(D and F)**. **(G)** Heatmap of differentially expressed STAT3-related genes.

**Figure S10.**
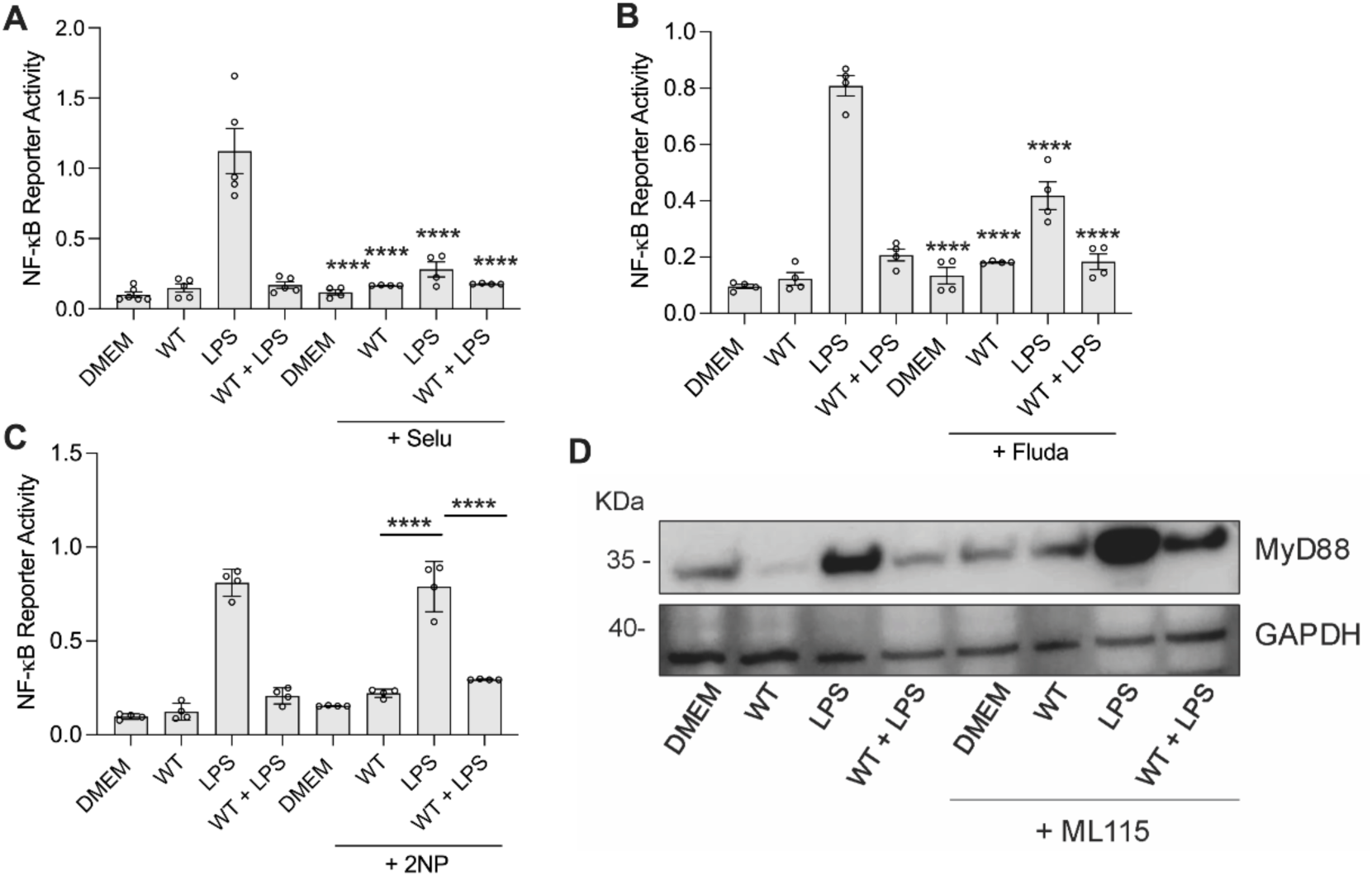
STAT1 does not play a role in suppressing NF-κB in *E. faecalis*- infected macrophages. RAW 264.7 macrophages were infected with the WT *E. faecalis* and incubated with cell culture media in presence or absence of LPS and **(A)** Selu (1 μM) and **(B)** Fluda (10 μM) and **(C)** 2NP (45 μM) for 6 h prior to measurement of NF-κB-driven SEAP reporter activity. The compounds were added 30 min before the beginning of the experiment. Data (mean ± SEM) are a summary of at least three independent experiments. Statistical analysis was performed using one-way ANOVA with Tukey’s multiple comparison tests where ****p<0.0001 as compared to LPS unless otherwise indicated. (D) Immunoblot was performed on proteins from RAW 264.7 macrophages uninfected or infected with WT in presence or absence of ML115 (± LPS) for 6 h and probed with antibodies raised against MyD88 and GAPDH.

**Figure S11.**
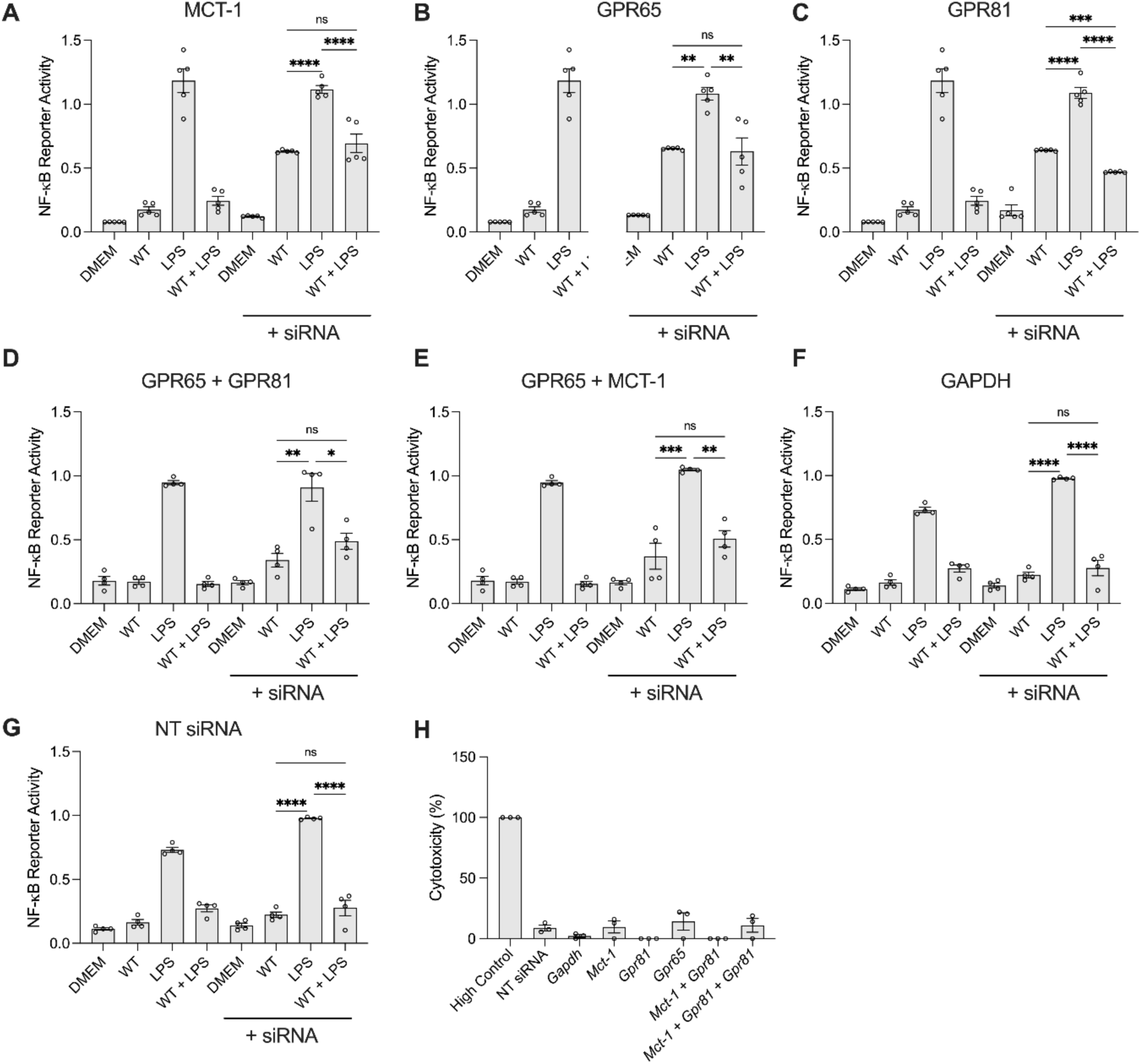
The lactate transporters MCT-1 and GPR81 are redundant in their role in the suppression of NF-κB activation of *E. faecalis* infection. **(A-G) RAW** 264.7 macrophages, with successfully silenced lactate transporters as indicated, were infected with the WT *E. faecalis* and incubated with cell culture media in the presence or absence of LPS for 6 h prior to measurement of NF-κB-driven SEAP reporter activity. Silencing the expression of *Mct-1*, *Gpr81* and *Gpr65* alone or in combination was achieved by siRNA. Data (mean ± SEM) are a summary of at least two independent experiments. NT siRNA was used so was *Gapdh* siRNAs controls for adverse effects of the technique. Statistical analysis was performed using ordinary one-way ANOVA followed by Tukey’s multiple comparison test ns, p>0.05; *p≤0.05; and **p<0.01; ***p<0.001 and ****p<0.0001. (H) Ldh assay was performed to evaluate potential cytotoxicity of the technique under the different siRNAs used.

**Figure S12.**
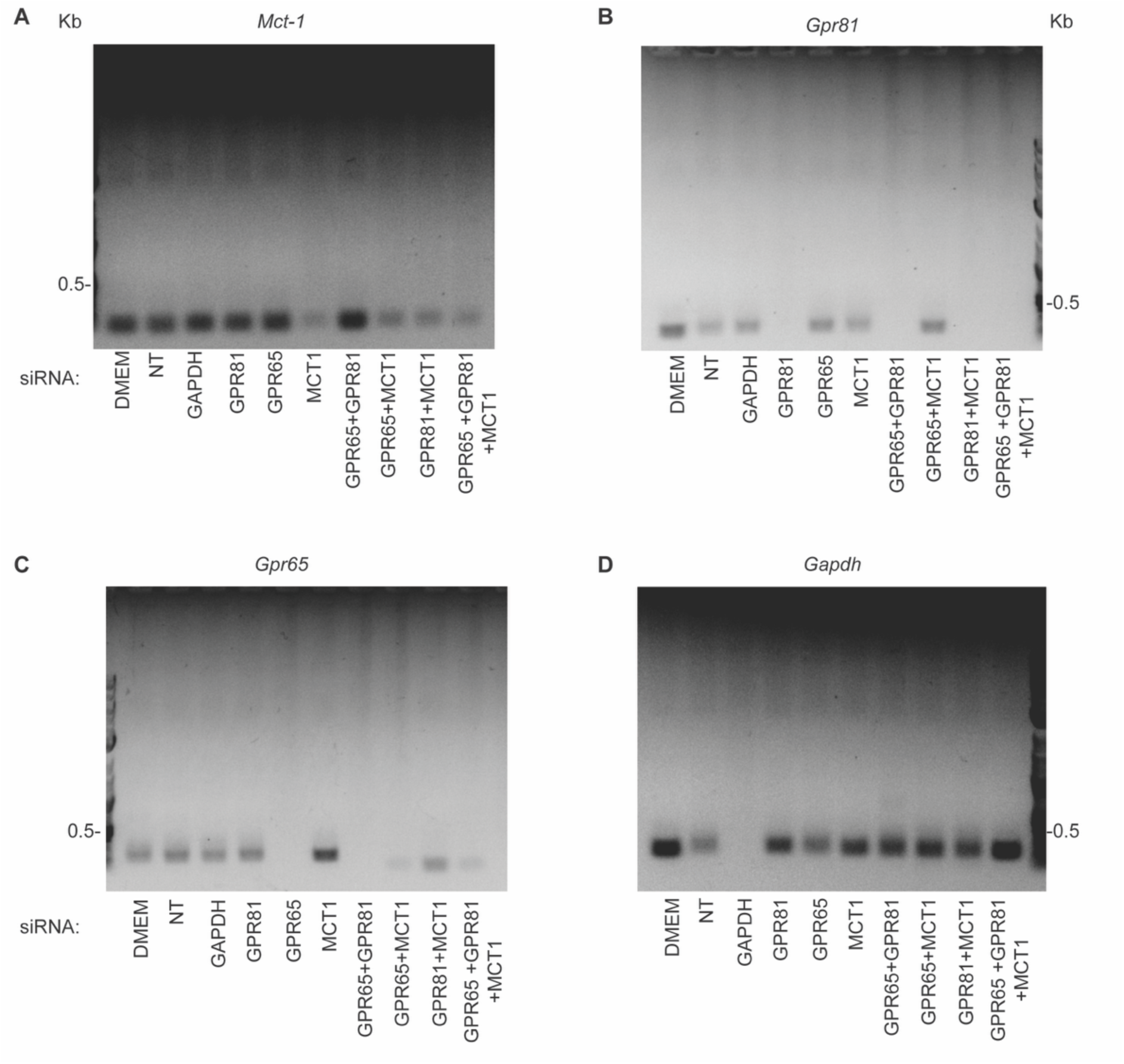
Successful knockdown of lactate transporters of interest was verified with RT-PCR. (A-D) Transcript levels of the silenced genes of interest were assessed with specific primers for each target. Negative control (NT) used was the Silencer Select Negative Control siRNAs that contain siRNAs with sequences that do not target any gene product.

**Figure S13.**
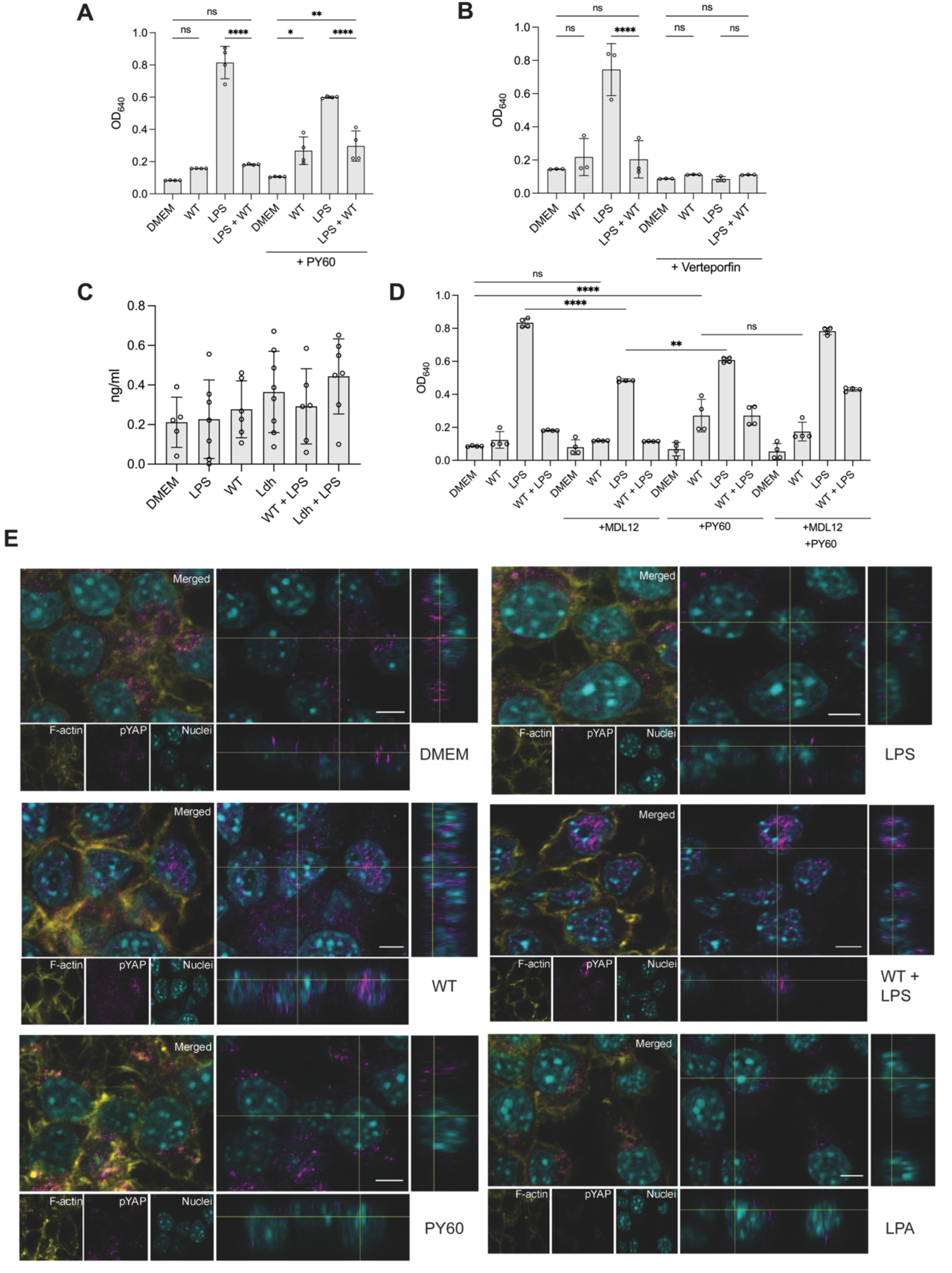
*E. faecalis* infection disturbs pYAP function and localization. RAW 264.7 macrophages were infected with the WT *E. faecalis* and incubated in cell culture media in the presence or absence of LPS and **(A)** PY-60 (10 μM), **(B)** verteporfin (10 μM), **(D)** MDL-12 alone or in combination with either PY-60 or verteporfin for 6 h prior to measurement of NF-κB-driven SEAP reporter activity. The compounds were added 30 min before the beginning of the experiment. Data (mean ± SEM) are a summary of at least three independent experiments. Statistical analysis was performed using one-way ANOVA with Tukey’s multiple comparison tests where NS, p>0.05, *p<0.05, **p<0.01, and ****p<0.0001. **(C)** Intracellular cAMP quantification of RAW264.7 cells uninfected or infected with WT or Ldh in presence or LPS. **(E)** CLSM orthogonal view of RAW264.7 macrophages at 6 hpi with WT in the presence or absence of LPS. Magenta, p-YAP; cyan, dsDNA stained with Hoechst 33342; yellow, F-actin stained with phalloidin. Images are representative of 2 independent experiments. Scale bar: 5 μm.

**Figure S14.**
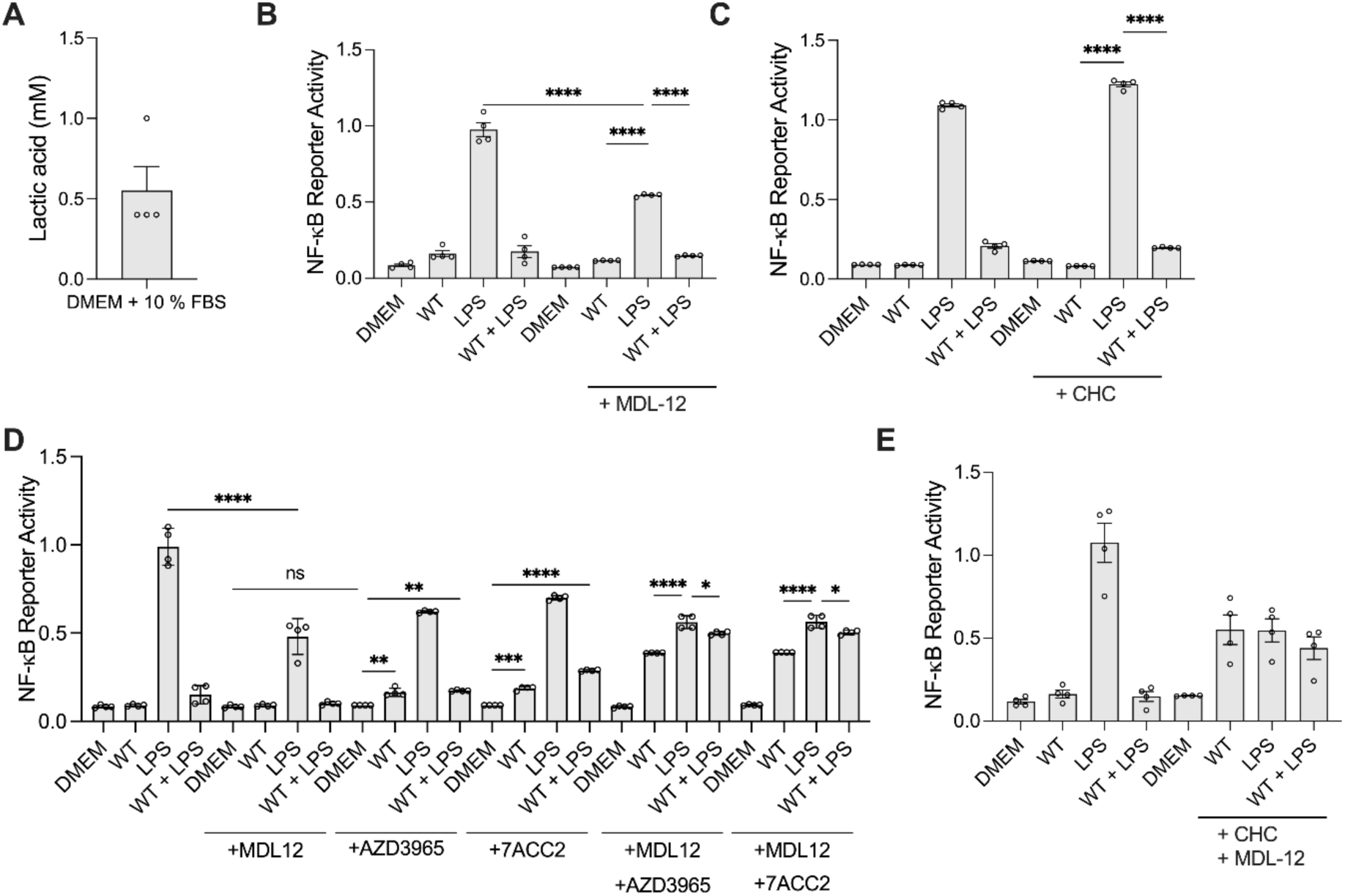
MCT1 mediate *E. faecalis* suppression of NF-κB activation in macrophages. **(A)** Lactic acid concentration of DMEM + 10 % FBS media. RAW 264.7 macrophages were infected with the WT *E. faecalis* and incubated in cell culture media in presence or absence of LPS and **(B)** CHC (5 mM) and **(C)** MDL-12 (20 μM),**(D-E)** AZD3965 (1μM), 7ACC2 (1μM) and CHC alone or in combination with MDL-12 for 6 h prior to measurement of NF-κB-driven SEAP reporter activity. The cell cultures were treated with compounds for 30 min before the beginning of the experiment. Data (mean ± SEM) are a summary of at least three independent experiments. Statistical analysis was performed using one-way ANOVA with Tukey’s multiple comparison tests where NS, p>0.05 and ****p<0.0001.

**Figure S15.**
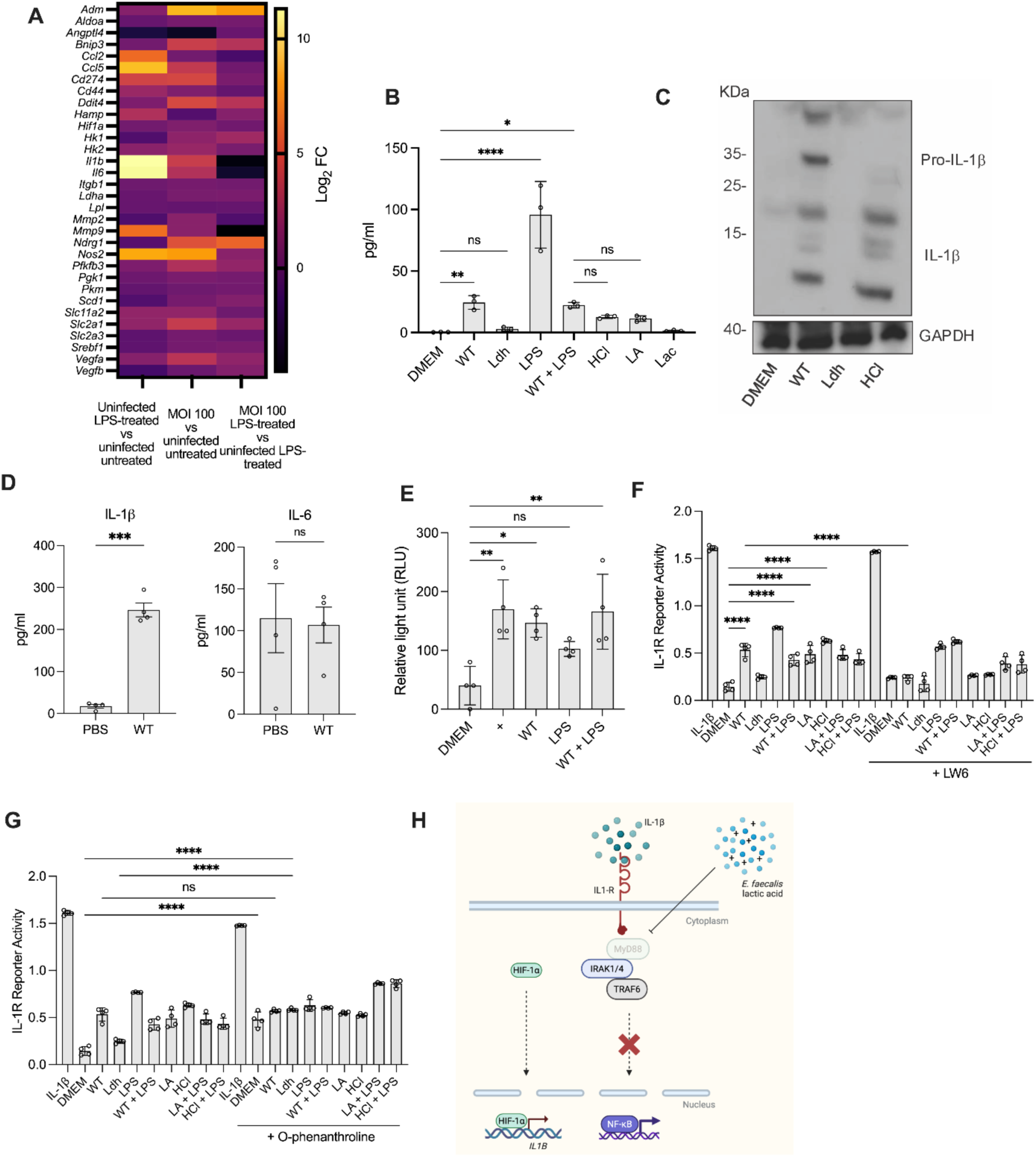
*E. faecalis* prevents IL-1β activation of NF-κB in RAW264.7 macrophages **(A**) Heatmap of differentially expressed HIF-1α-related genes. **(B)** Levels of IL-1β from the supernatants of uninfected or infected with WT or Ldh RAW264.7 macrophages treated with LPS, HCl, LA or sodium lactate (Lac). **(C)** Immunoblot performed on protein extracted from RAW 264.7 macrophages infected with WT or Ldh for 6 h and probed with antibodies specific for IL-1β and GAPDH. **(D)** Levels of IL-1β and IL-6 from uninfected or WT-infected wound lysates at 24 hpi. Each dot represents one mouse. **(E)** HIF-1α reporter activity assay in RAW264.7 macrophages that were uninfected or infected with WT treated with O-phenanthroline (+) or LPS. **(F-G)** IL-1R reporter activity assay with HEK-Blue IL-1R cells treated with 20 μl bacteria-free supernatants from uninfected or infected with WT or Ldh RAW264.7 macrophages that were left untreated or treated with LPS, HCl, or LA for 6 h. Some RAW264.7 cells pre- treated with LW6 or O-phenanthroline for 30 min prior to the beginning of the assay. **(H)** IL-1β activation of NF-kB signaling pathway. Data (mean ± SEM) are summary of at least 2 independent experiments **(B-G)**. Statistical analysis was performed using unpaired t test with Welch’s correction **(D)**, ordinary one-way ANOVA followed by Tukey’s multiple comparison test **(B, E)** or Fisher’s LSD **(F-G),** ns, p>0.05; *p≤0.05; **p<0.01, ***p<0.001, and ****p<0.0001.

**Figure S16.**
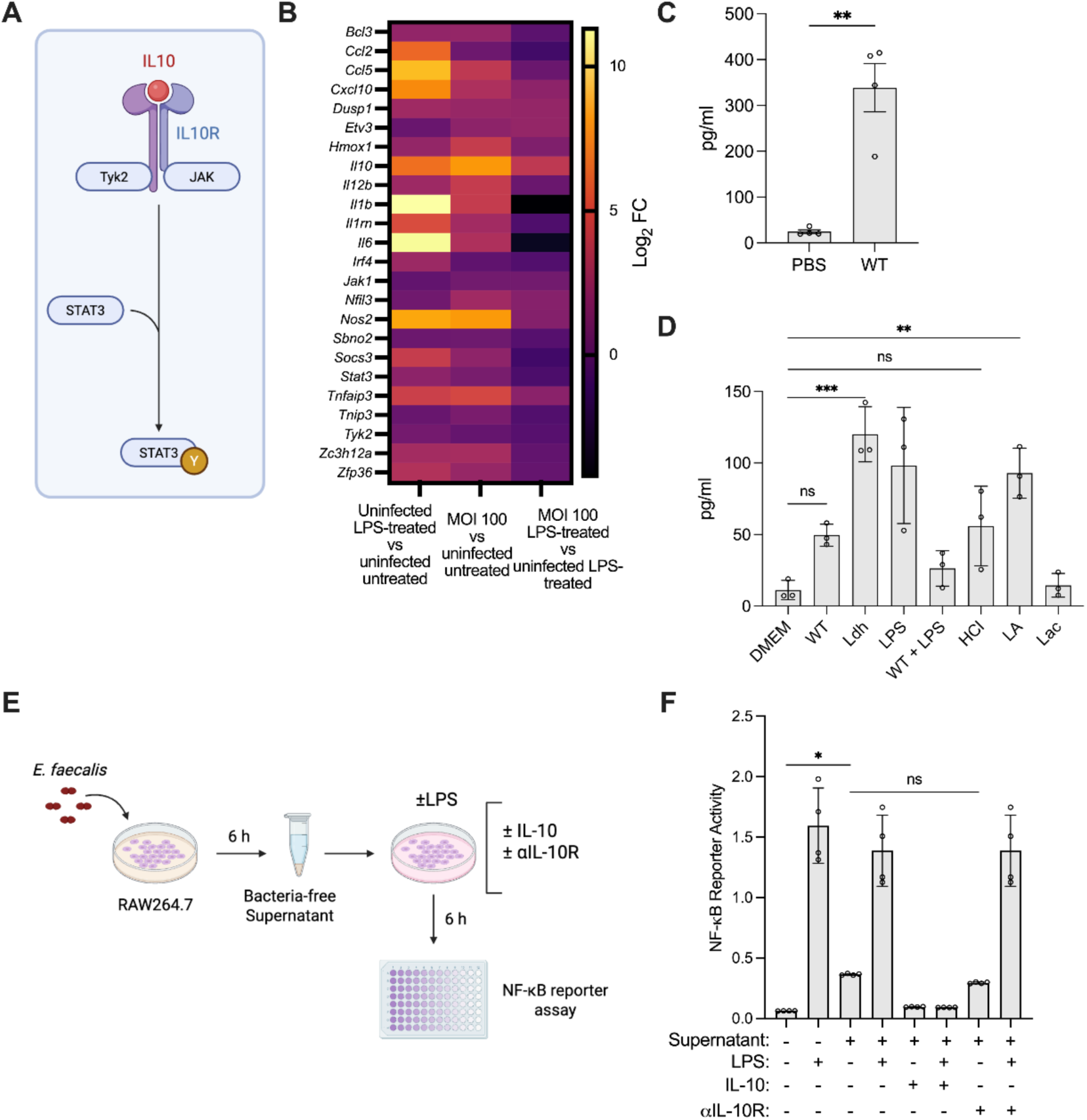
IL-10 from *E. faecalis* infected macrophages does not suppress NF- κB during enterococcal infection. **(A**) IL-10 signaling pathway. **(B)** Heatmap of differentially expressed IL-10-related genes. **(C)** IL-10 levels from uninfected or WT-infected wound lysates at 24 hpi. Each dot represents one mouse. **(D)** Levels of IL-10 from the supernatants of uninfected or infected with WT or Ldh RAW264.7 macrophages treated with LPS, HCl, lactic acid (LA) or sodium lactate (Lac). **(E-F)** NF-κB reporter activity assay with RAW264.7 cells treated with 20 μl bacteria-free supernatants from uninfected or 6 h WT infected RAW264.7 macrophages that were left untreated or treated with LPS. RAW264.7 cells were pre-treated with IL-10 or antibody αIL-10R for 30 min prior to the beginning of the assay. Data (mean ± SEM) are summary of at least 3 independent experiments **(C-D and F)**. Statistical analysis was performed using unpaired t test with Welch’s correction **(C)**, ordinary one-way ANOVA followed by Tukey’s multiple comparison test **(D)** or Fisher’s LSD **(F),** ns, p>0.05; *p≤0.05; **p<0.01, ***p<0.001, and ****p<0.0001.

**Figure S17.**
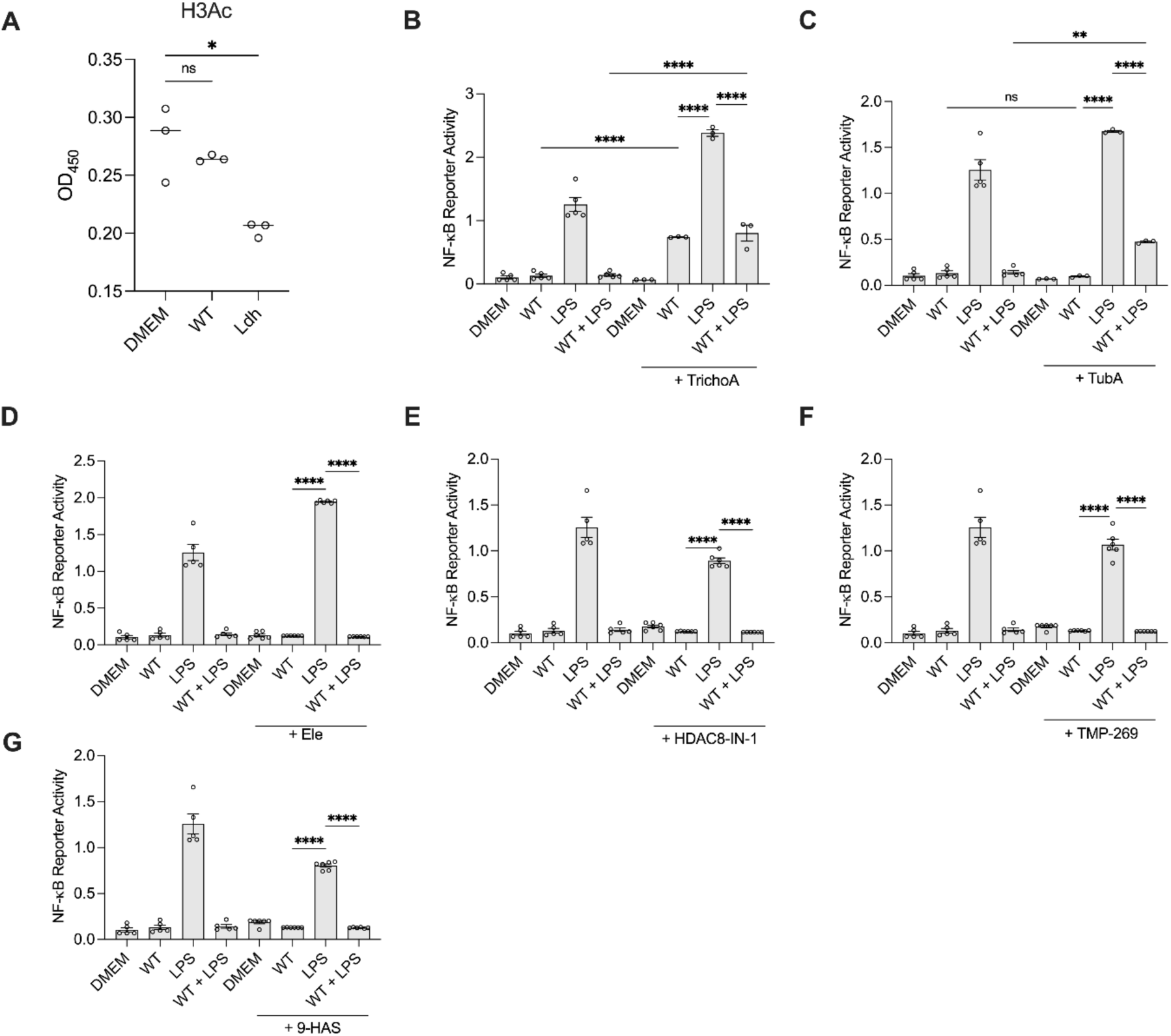
HDACs likely play a partial role in mediating *E. faecalis* suppression of NF-kB activation in macrophages. **(A)** Relative abundance of H3Ac by ELISA of uninfected or WT and Ldh infected macrophages. RAW 264.7 macrophages were infected with the WT strain and incubated with cell culture media in the presence or absence of LPS and **(B)** TrichoA (10 nM) and **(C)** TubA (15 nM), **(D)** Ele (1 μM), **(E)** HDAC8-IN-1 (100 nM), **(F)** TMP-269 (10 μM) and **(G)** 9-HAS (100 μM) for 6 h prior to measurement of NF-κB-driven SEAP reporter activity. The compounds were added 30 min before the beginning of the experiment. Data (mean ± SEM) are a summary of at least three independent experiments. Statistical analysis was performed using one- way ANOVA with Tukey’s multiple comparison tests where ns, p>0.05; *p≤0.05; and ****p<0.0001.

**Figure S18.**
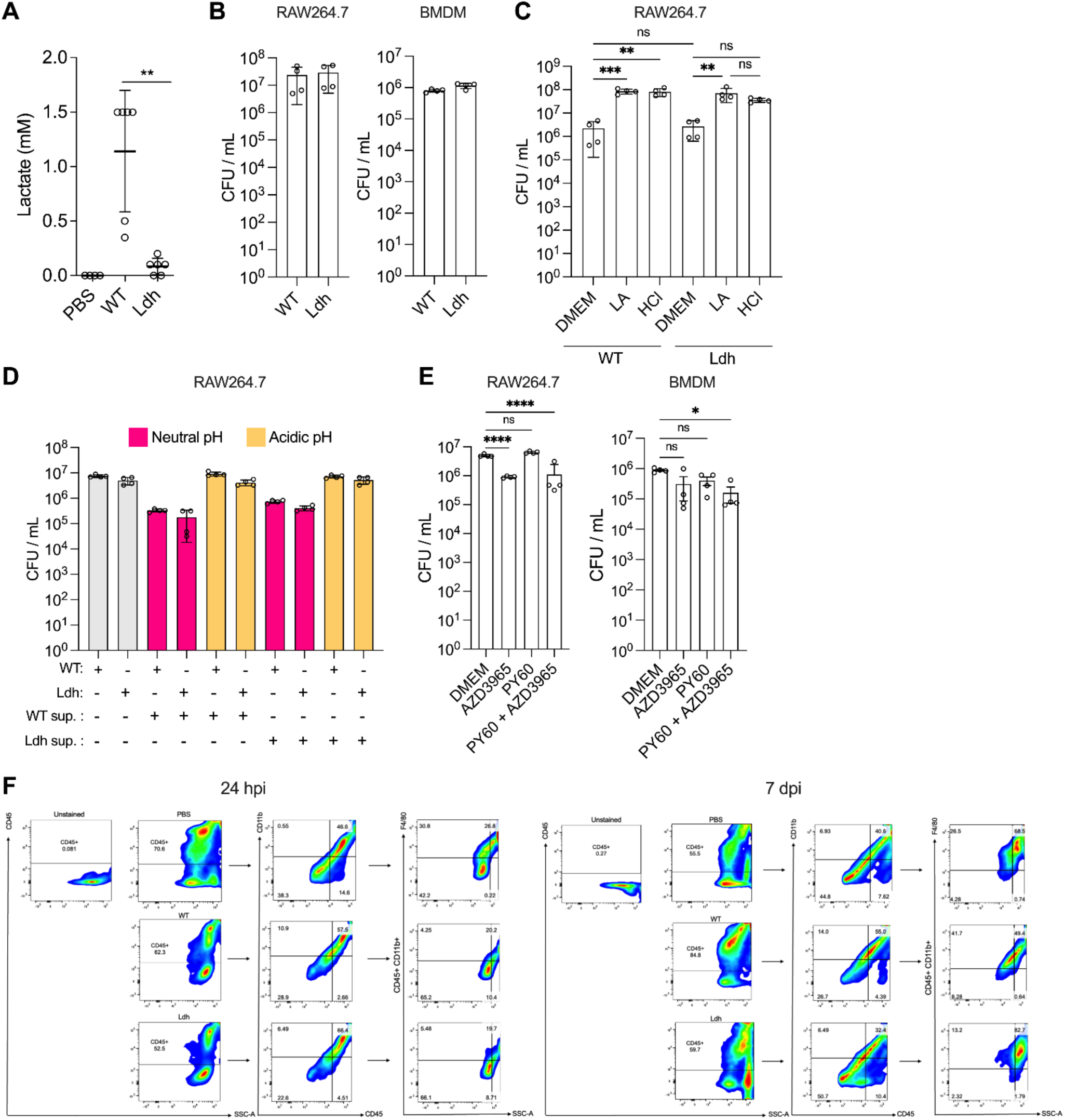
Lactic acid reduces macrophage killing of bacteria. **(A)** Lactate concentration of wound measured at 24 hpi for the indicated condition. **(B-E)** RAW264.7 cells or BMDMs were infected with either WT or Ldh without any other treatment **(B)**, in the presence of LA (40 mM) or HCl to lower the extracellular pH **(C)**, bacterial free supernatant from WT or Ldh cultures that were neutralized with NaOH or acidified with HCl **(D)**, or in the presence of AZD3965 (10 μM), PY60 (10 μM), or both **(E).** Intracellular bacterial CFU was quantified. Statistical analysis was performed using one-way ANOVA with Tukey’s multiple comparison tests where p>0.05; *p≤0.05; **p<0.01, ***p<0.001, and ****p<0.0001**. (F)** Representative gating strategy for macrophages (CD45^+^ CD11b^+^ F4/80^+^) from wounds that were treated with PBS or infected with WT or Ldh strains at 24 hpi or 7 dpi. The number indicate percentages of cells within the gated areas. Recorded MFI levels of MyD88 and p-ERK derived from this group.

**Figure S19.**
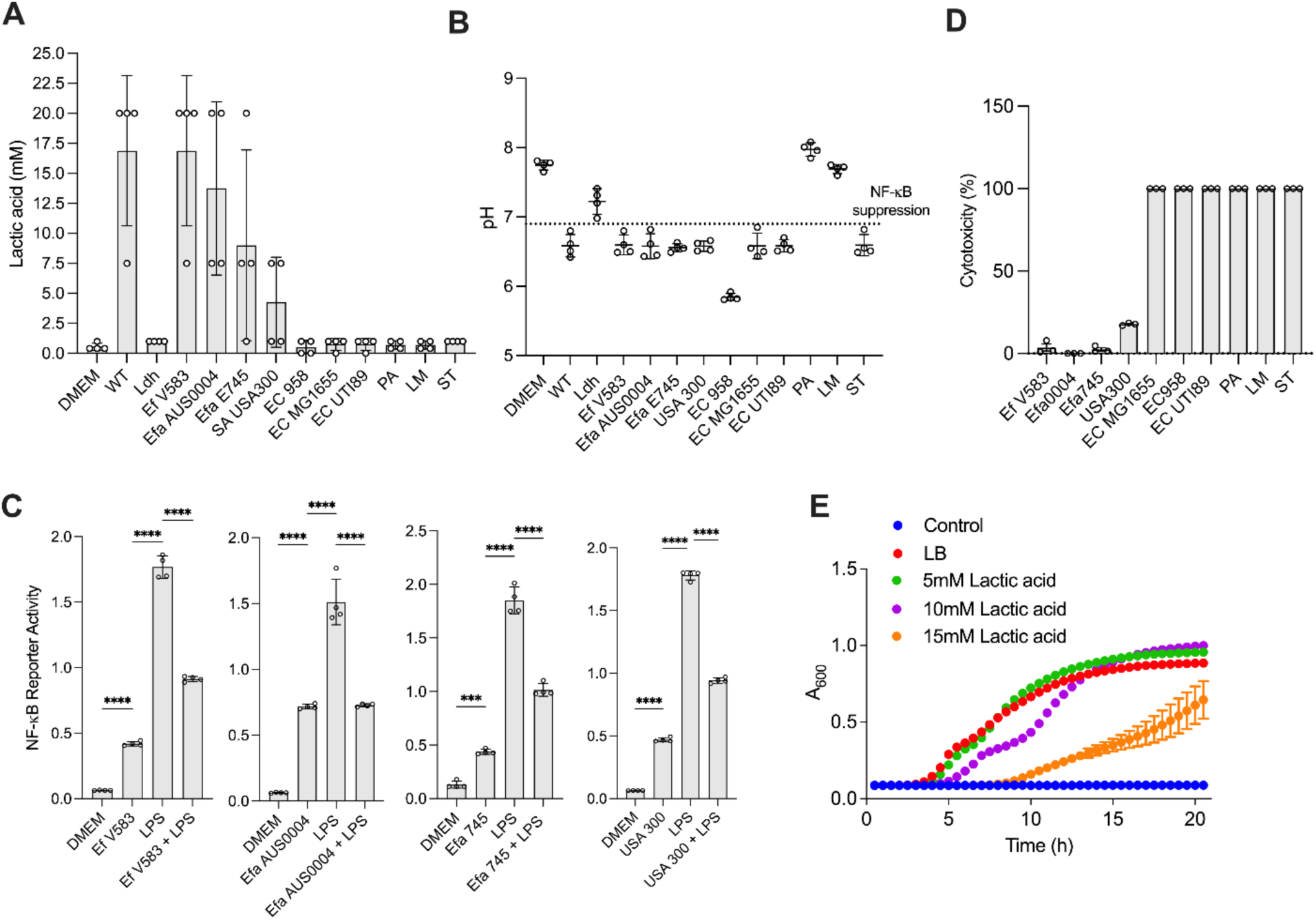
Lactic acid-producing species acidify the medium and suppress macrophage NF-κB activation at high MOI. **(A)** Lactic acid concentration of several bacterial species grown in cell culture media at initial MOI 100 measured after 6 hpi. **(B)** Extracellular pH following bacterial growth in cell culture media with starting inoculum equivalent to MOI 100 measured after 6 hpi. **(C)** RAW 264.7 macrophages were infected with the non-cytotoxic bacterial species of *E. faecalis* strain V583, *E. faecium* strains AUS0004 and E745 and *S. aureus* strain USA300 at initial MOI 100 for 6 h prior to NF-kB reporter assay. **(D)** RAW 264.7 macrophages were infected with different bacterial species at initial MOI 100 for 6 h prior to LDH assay to measure cytotoxicity. Data (mean ± SEM) are summary of at least 2 independent experiments **(A-D)**. Statistical analysis was performed using ordinary one-way ANOVA followed by Tukey’s multiple comparison test **(C)** ***p<0.001, and ****p<0.0001. **(E)** Growth curve for *E. coli* strain UTI89 grown overnight in LB media supplemented with different concentration of lactate.

## Supplementary Tables

**Table S1.** Differentially expressed NF-κB/ AP-1-related genes

|  |  | Condition |  |  |  |  |  |
| --- | --- | --- | --- | --- | --- | --- | --- |
|  |  | Uninfected LPS-treated vs uninfected untreated |  | MOI 100 untreated vs uninfected untreated |  | MOI 100 LPS-treated vs uninfected LPS-treated |  |
| Gene | Gene product | Log2FC | Significant? | Log2FC | Significant? | Log2FC | Significant? |
| NF-κB |  |  |  |  |  |  |  |
| *Tlr4* | TLR4 | -0.84 | No | -0.66 | No | -1.33 | No |
| *Cd14* | CD14 | 2.15 | Yes | 0.31 | No | -2.13 | Yes |
| *Myd88* | MyD88 | 1.11 | No | -1.06 | No | -1.86 | Yes |
| *Tirap* | TIRAP | 0.38 | No | -1.03 | No | -1.16 | No |
| *Ly96* | MD-2 | 0.89 | No | 0.29 | No | -0.14 | No |
| *Ticam1* | TRIF | 1.84 | No | 1.59 | Yes | 0.28 | No |
| *Ticam2* | TRAM | -1.35 | No | 1.28 | No | 2.48 | Yes |
| *Traf6* | TRAF6 | 0.46 | No | 0.57 | No | -0.09 | No |
| *Map3k7* | TAK1 | -0.06 | No | 0.07 | No | 0.17 | No |
| *Chuk* | IKKα | 0.32 | No | 0.35 | No | 0.004 | No |
| *Ikbkb* | IKKβ | 0.37 | No | -0.02 | No | -0.13 | No |
| *Nfkbia* | IKBα | 2.95 | Yes | 3.01 | Yes | 0.95 | No |
| *Rel* | c-Rel | 1.60 | Yes | 1.70 | Yes | 0.29 | No |
| *Rela* | p65 | -0.07 | No | 0.41 | No | 0.59 | No |
| *Relb* | RelB | 1.26 | No | 2.33 | Yes | 1.01 | No |
| *Nfkb1* | p105 | 3.20 | Yes | 2.33 | Yes | 0.11 | No |
| *Nfkb2* | p100 | 1.98 | Yes | 1.84 | Yes | 0.89 | No |
| AP-1 |  |  |  |  |  |  |  |
| *Mapk14* | p38 | -0.62 | No | -1.87 | Yes | -0.94 | No |
| *Mapk8* | JNK | 0.28 | No | 0.54 | No | 0.40 | No |
| *Jun* | cJun | 0.47 | No | 0.82 | No | -0.34 | No |
| *Fos* | cFos | 0.39 | No | -0.65 | No | -1.79 | Yes |
| **Common** |  |  |  |  |  |  |  |
| *Rps6ka5* | MSK1 | -2.58 | Yes | -1.87 | Yes | -0.99 | No |
| *Rps6ka4* | MSK2 | 0.29 | No | -0.30 | No | 0.22 | No |
| *Mapk3* | ERK1 | 0.22 | No | 0.87 | No | 1.00 | No |
| *Mapk1* | ERK2 | -0.09 | No | 0.11 | No | 0.05 | No |

**Table S2.**
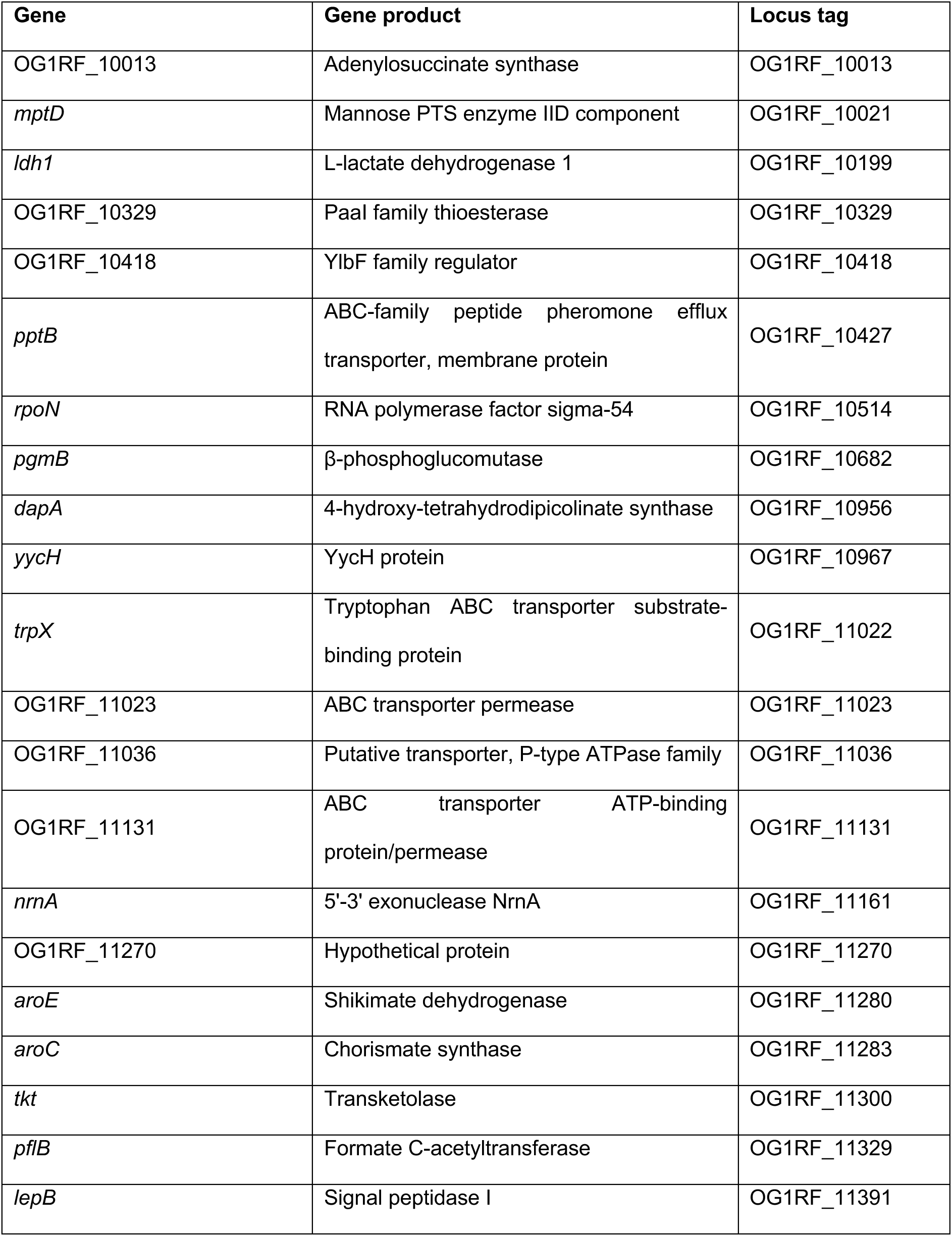
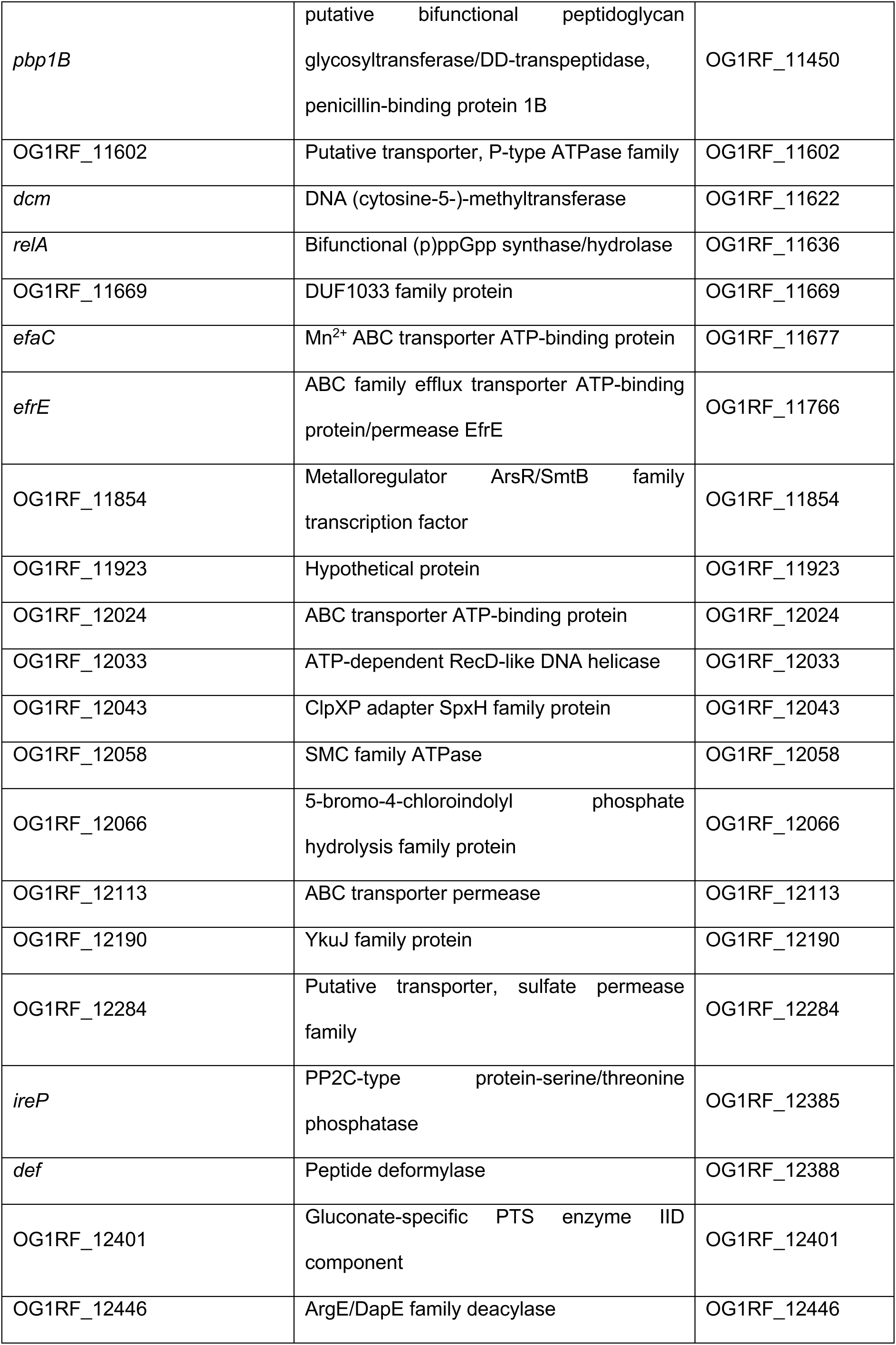
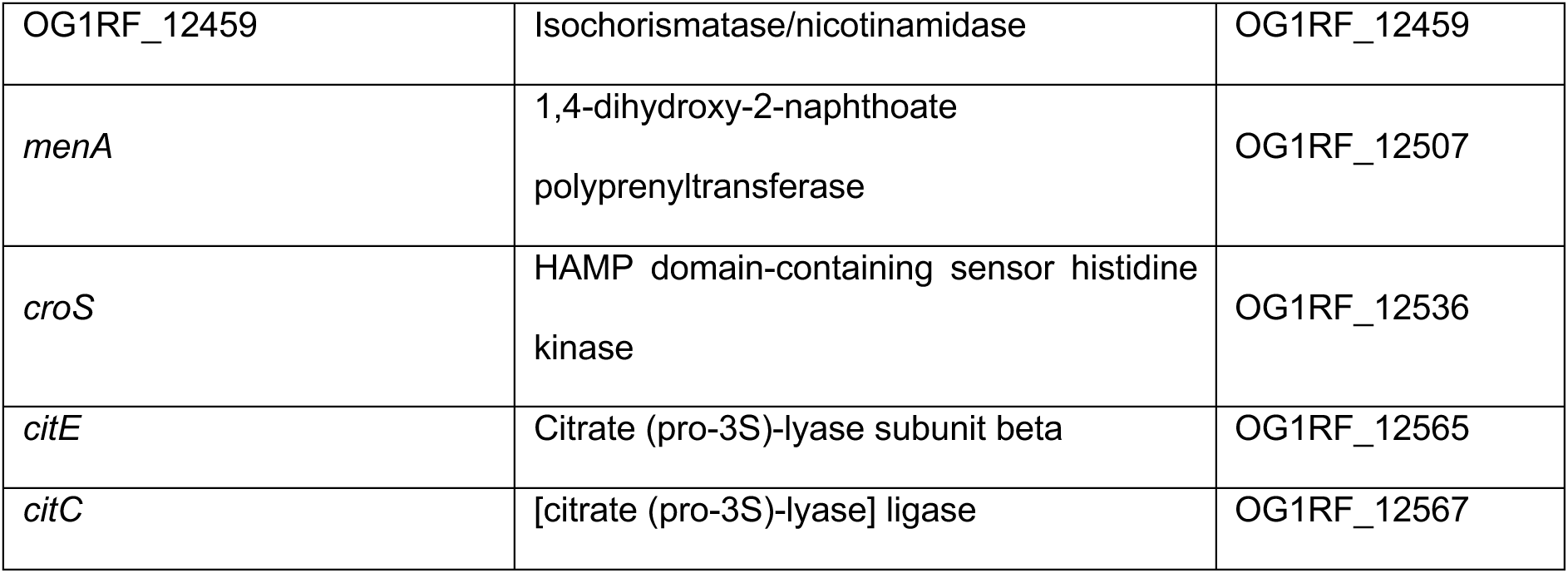
List of hits after transposon screen for genes that contribute to suppress NF-κB activation in macrophages

**Table S3.**
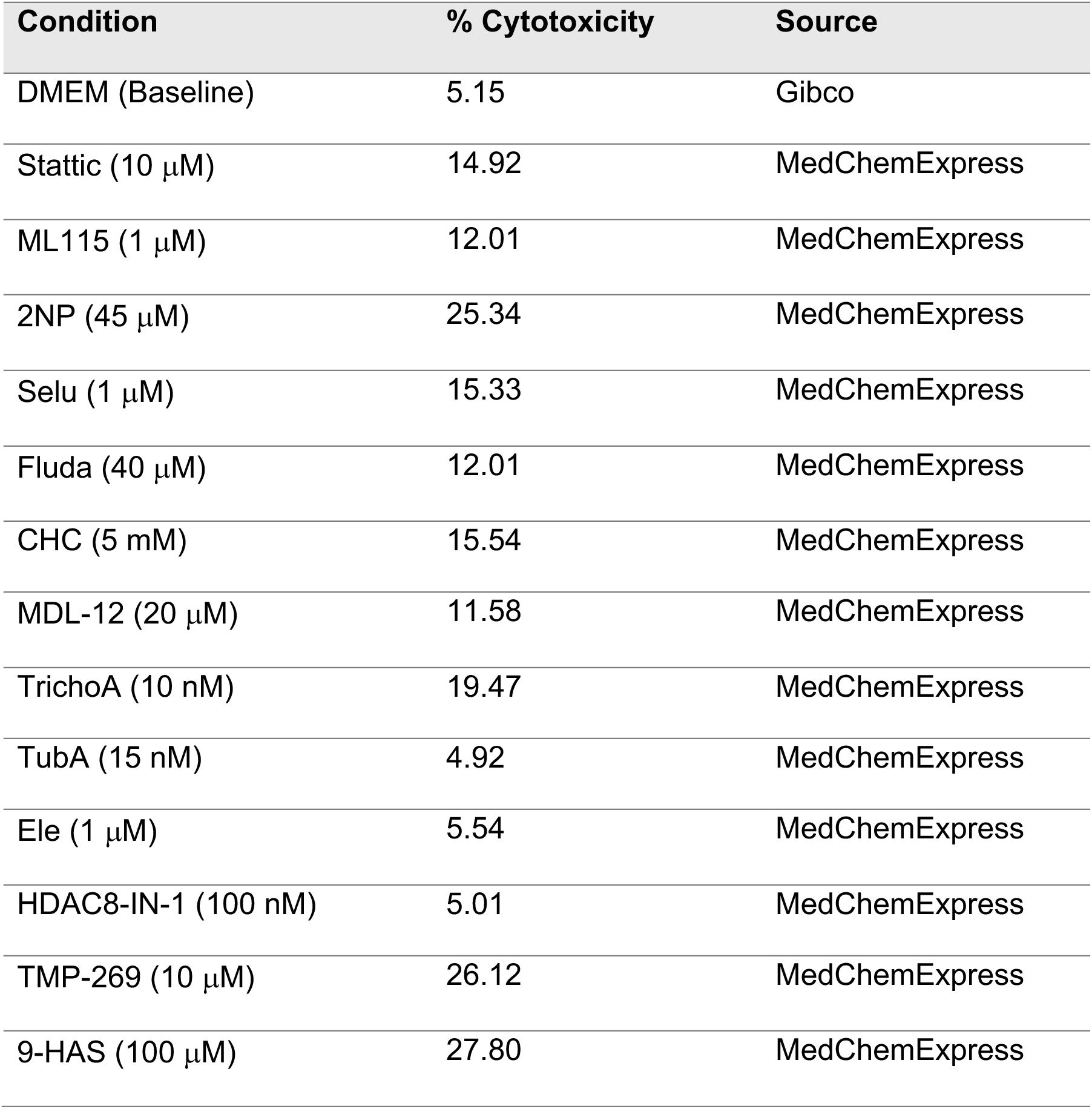
Cytotoxicity as measured by LDH assay of compounds used in this study.

| Condition | % Cytotoxicity | Source |
| --- | --- | --- |
| DMEM (Baseline) | 5.15 | Gibco |
| Stattic (10 μM) | 14.92 | MedChemExpress |
| ML115 (1 μM) | 12.01 | MedChemExpress |
| 2NP (45 μM) | 25.34 | MedChemExpress |
| Selu (1 μM) | 15.33 | MedChemExpress |
| Fluda (40 μM) | 12.01 | MedChemExpress |
| CHC (5 mM) | 15.54 | MedChemExpress |
| MDL-12 (20 μM) | 11.58 | MedChemExpress |
| TrichoA (10 nM) | 19.47 | MedChemExpress |
| TubA (15 nM) | 4.92 | MedChemExpress |
| Ele (1 μM) | 5.54 | MedChemExpress |
| HDAC8-IN-1 (100 nM) | 5.01 | MedChemExpress |
| TMP-269 (10 μM) | 26.12 | MedChemExpress |
| 9-HAS (100 μM) | 27.80 | MedChemExpress |

**Table S4.** Strains used in this study

| Strain | Reference |
| --- | --- |
| *E. faecalis* OG1RF | [102] |
| *E. faecalis* OG1RF *Ldh* ($\Delta ldh1 \Delta ldh2$) | This study |
| *E. faecalis* OG1RF $\Delta ldh1$ | This study |
| *E. faecalis* OG1RF $\Delta ldh2$ | This study |
| *E. faecalis* OG1RF *pldh1* (+ pGCP123::*ldh1*) | This study |
| *E. faecalis* V583 | [103] |
| *E. coli* MG1655 | [104] |
| *E. coli* EC958 | [105] |
| *E. coli* UTI89 | [106] |
| *S. aureus* USA300 | [107] |
| *S. typhimurium* 14028S | [108] |
| *P. aeruginosa* PAO1 | [109] |
| *E. faecium* E745 | [110] |
| *E. faecium* AUS0004 | [111] |
| *Listeria monocytogenes* ATCC19115 | [112] |

**Table S5.**
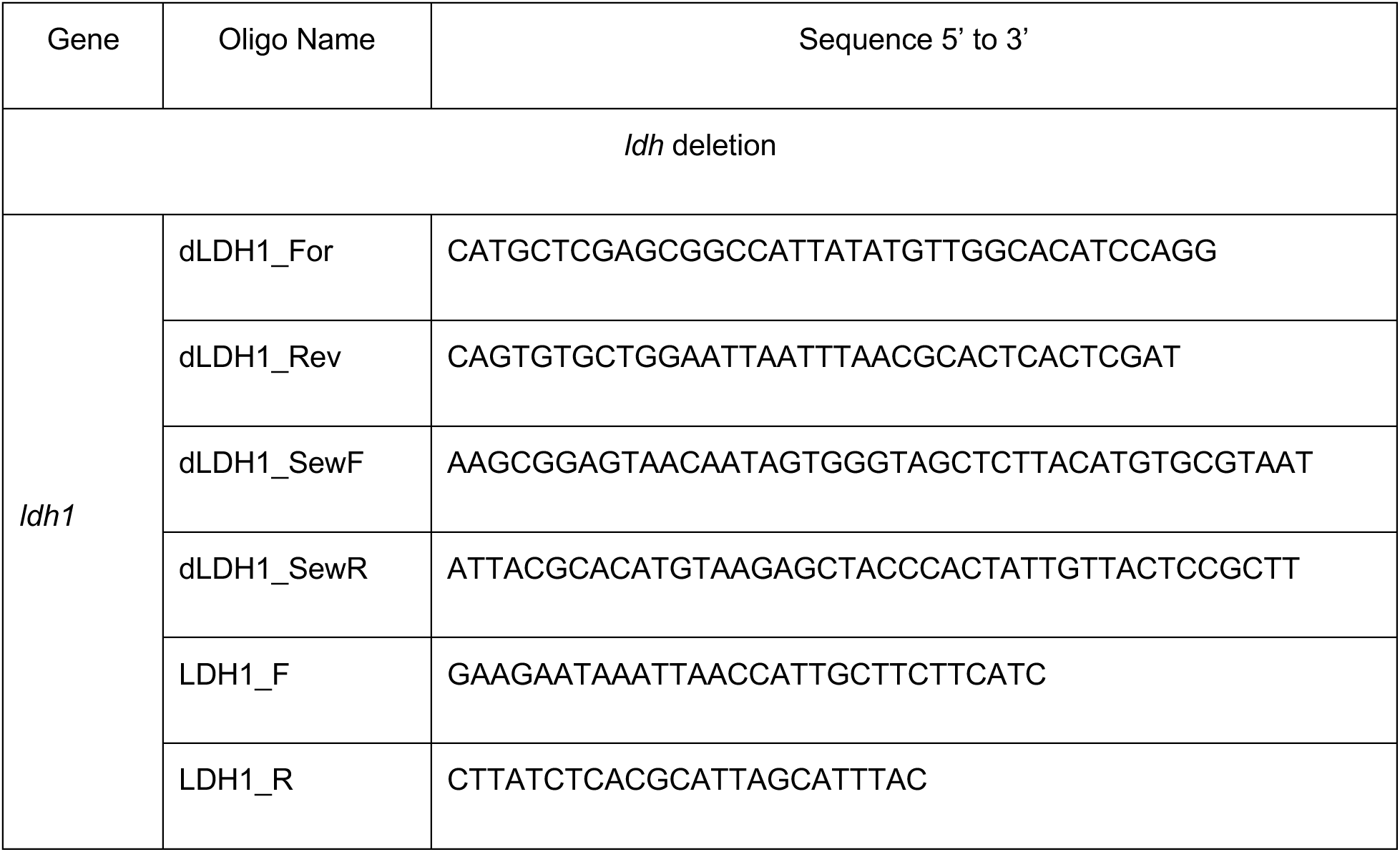
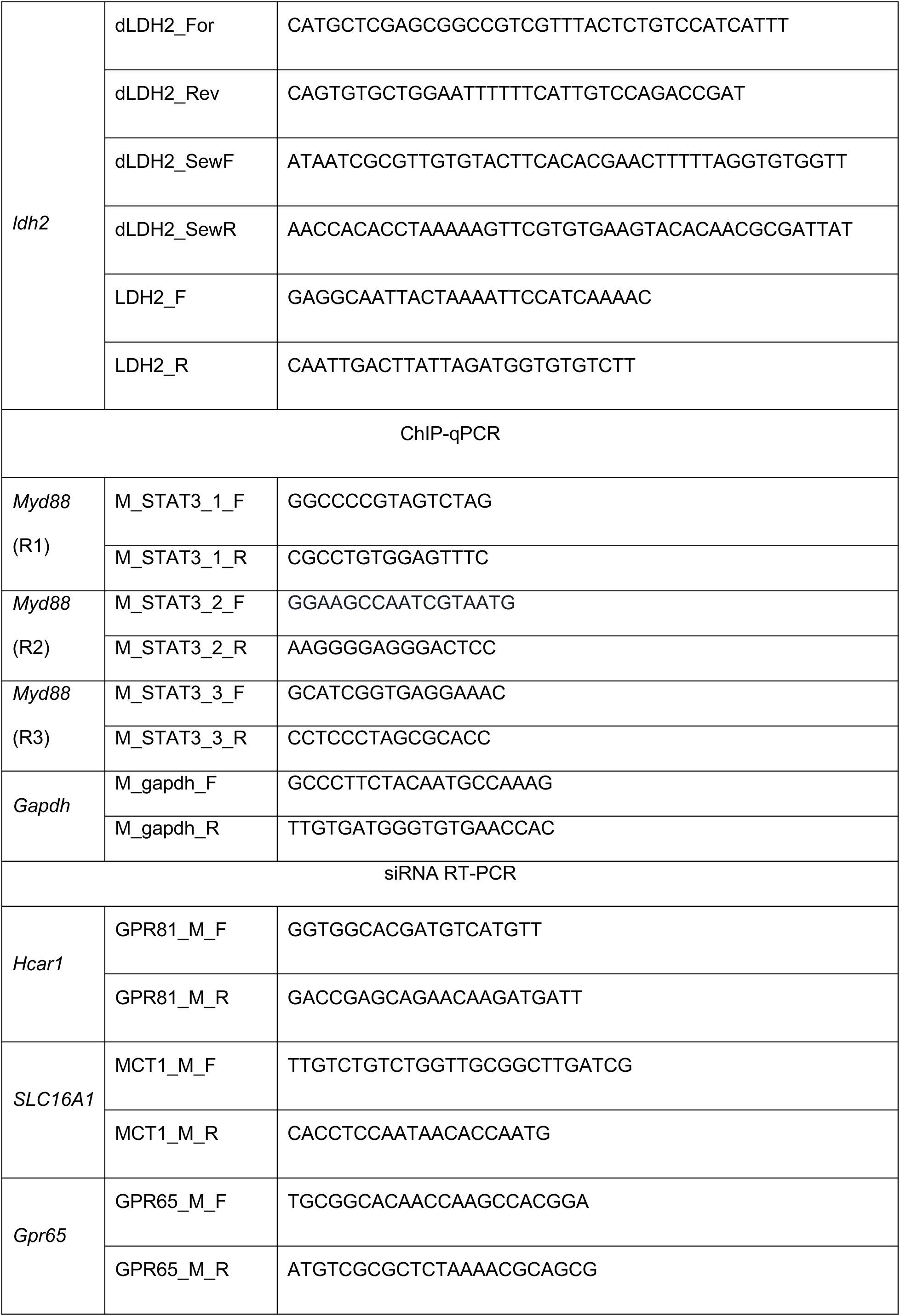
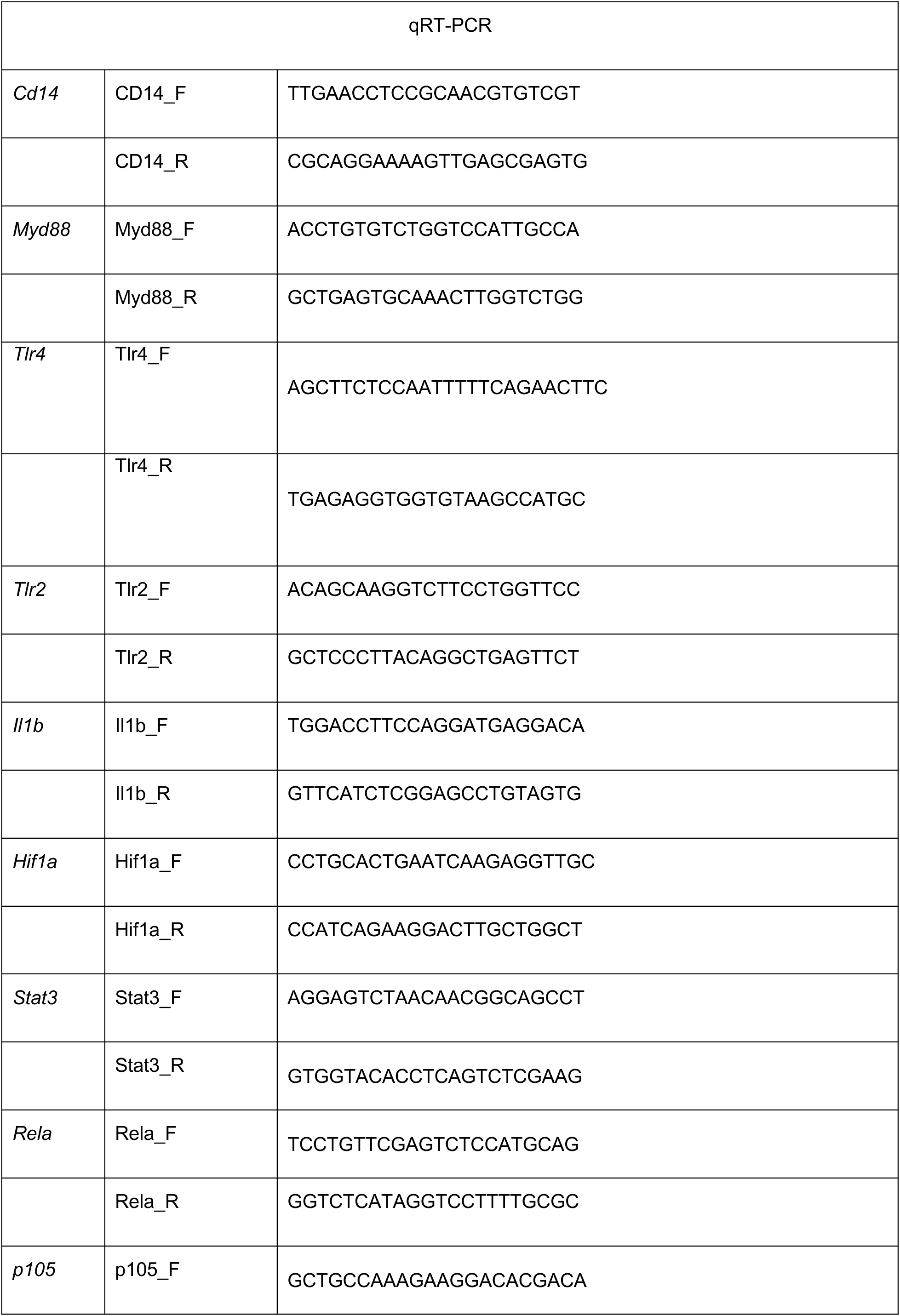
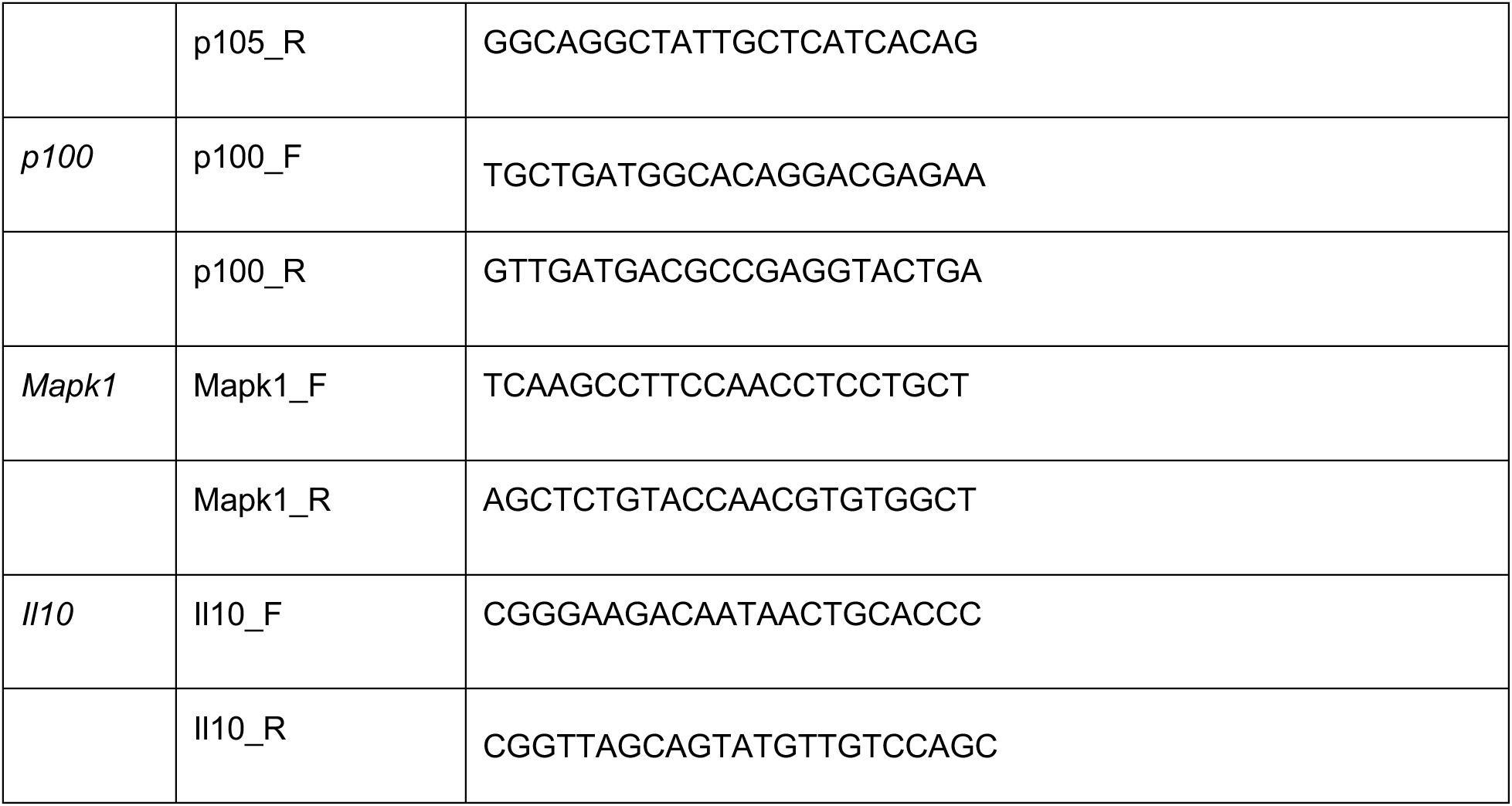
List of primers used in this study

